# A Receptor Like Cytoplasmic Kinase evolved in *Aeschynomene* legumes to mediate Nod-independent rhizobial symbiosis

**DOI:** 10.1101/2024.07.10.602847

**Authors:** Natasha Horta Araújo, David Landry, Johan Quilbé, Marjorie Pervent, Nico Nouwen, Christophe Klopp, Julie Cullimore, Djamel Gully, Laurent Brottier, Carole Pichereaux, Martin Racoupeau, Maëlle Rios, Frédéric Gressent, Clémence Chaintreuil, Clare Gough, Eric Giraud, Benoit Lefebvre, Jean-François Arrighi

## Abstract

Many plants interact symbiotically with arbuscular mycorrhizal (AM) fungi to enhance inorganic phosphorus uptake, and legumes also develop a nodule symbiosis with rhizobia for nitrogen acquisition. Establishment and functioning of both symbioses rely on a common plant signaling pathway activated by structurally related Myc- and Nod-factors. Recently, a SPARK Receptor-like-Kinase (RLK)/Receptor-like Cytoplasmic Kinase (RLCK) complex was shown to be essential for AM in both monocot and dicot plants. Here, we show that in *Aeschynomene* legumes the RLCK component of this receptor complex has evolved following a gene duplication event and mediates a unique nodule symbiosis that is independent of rhizobial Nod factors. In *Aeschynomene evenia*, *AeRLCK2* is crucial for nodule initiation but not for AM. Additionally, AeRLCK2 physically interacts with and is phosphorylated by the Cysteine-rich RLK, AeCRK, also required for nodulation. This work reveals a novel evolutionary origin of this Nod-independent symbiosis from AM.

## Introduction

Plants have evolved a range of mutualistic partnerships with soil-dwelling microorganisms to enhance their nutrient uptake. The oldest and most widespread symbiosis is the association with Glomeromycotina fungi, referred to as arbuscular mycorrhizal (AM) fungi. AM fungi develop extensive hyphal networks to take up inorganic phosphorus (Pi) and other nutrients from the soil. They also colonize plant roots and form intracellular branched strucures called arbuscules, through which they provide these nutrients to plants^(Rich-2021)^. Plant species within the nitrogen (N)-fixing clade are able to develop an additional symbiosis with diazotrophic bacteria that are hosted in root nodules^(vanVelzen-2019)^. This interaction allows the plants to access the abundant atmospheric N by converting it into ammonium^(Roy-2020)^. By providing Pi and N to the plants, these symbioses are central to the functioning of natural ecosystems and the productivity of agro-systems.

Genetic studies of *Oryza sativa* (rice) and model legumes (such as *Lotus japonicus* and *Medicago truncatula*), establishing a symbiosis with AM fungi and/or rhizobia, have demonstrated that the evolution of nitrogen-fixing symbiosis has co-opted perception, signalling, and infection mechanisms essential for the establishment of AM ^(Radhakrishan-2020)^. Chitooligosaccharides (COs) and lipochitosaccharides (LCOs) produced by AM fungi and LCOs produced by rhizobia (known as Nod factors) are perceived by distinct plasma membrane receptor-like kinases of the LysM-RLK subfamily^(Feng-2019;Buendia-2018)^. These signals are then transduced by SymRK (Symbiosis Receptor Kinase), a RLK belonging to the LRR-RLK subfamily, which initiates a common symbiosis signalling pathway leading to transcriptional reprogramming^(Roy-2020;Gobbato-2015)^.

This transcriptional reprogramming drives infection by both AM fungi and rhizobia, culminating in their intracellular accomodation either into the plant root or into nodules^(Roy-2020)^. Another RLK belonging to the SPARK-RLK subfamily, KIN3 (KINASE 3), is crucial for the arbuscule formation during AM^(Leng-2023;Irving-2022)^. Recently, KIN3 has been shown to interact with two receptor-like cytoplasmic kinase (RLCK) paralogs, AMK8 and AMK24, which also play a key role in arbuscule formation in *L. japonicus*^(Leng-2023)^. This RLK/RLCK complex was found to have a conserved role in AM in rice^(Leng-2023;Montero-2021;Roth-2018;Bravo-2016)^. Interestingly, *KIN3* orthologs are only found in plants able to form AM and *KIN3* is dispensable for rhizobial symbiosis in *M. truncatula*^(Irving-2022)^. This suggests that the KIN3-interacting RLCKs may also be not required for nodulation.

We challenged this view through the genetic analysis of nodulation in *Aeschynomene evenia*. This legume species has emerged as a model of choice for the study of a unique N-fixing symbiosis with photosynthetic *Bradyrhizobium* strains that do not produce Nod factors^(Chaintreuil-2016;Arrighi-2012)^. While the molecular mechanisms underlying the activation of this so-called Nod-independent symbiosis remain poorly understood, progress has been made in recent years, thanks to the availability of a reference genome and a collection of EMS nodulation mutants for *A. evenia*^(Quilbé-2022;Quilbé-2021)^. It is now known that many components of the common symbiotic signalling pathway are conserved in *A. evenia.* But that this pathway is likely activated by other type of receptor proteins. Notably, a Cysteine-rich RLK, AeCRK, was recently shown to be required to establish the Nod-independent symbiosis^(Quilbé-2022;Quilbé-2021).^

In this work, we identified *AeRLCK2*, a *RLCK* homolog to *L. japonicus AMK8*, as required for the Nod-independent symbiosis. Through the genetic, phenotypic and molecular characterization of 12 allelic mutants, we show that *AeRLCK2* is essential for nodulation but dispensable for AM. Furthermore, we provide evidence that *AeRLCK2* originated from a recent tandem duplication event and offer clues to explain how it has evolved for a role in the Nod-independent symbiosis. Finally, we demonstrate that AeRLCK2 physically interacts with and serves as a phosphorylation substrate for AeCRK. These findings shed new light on the evolution of RLCKs in plant symbioses and further elucidate the mechanisms by which the Nod-independent nodulation process is triggered in *Aeschynomene* legumes.

## Results

### Mutant-based identification of AeRLCK2, a symbiosis receptor-like cytoplasmic kinase

To identify novel genes involved in establishment of the Nod-independent symbiosis, we analyzed a set of 12 uncharacterized nodulation mutants obtained from screening in greenhouse conditions of an ethyl methane sulfonate (EMS)-mutagenized population of *A. evenia* CIAT22838 (Supplementary Table 1)^(Quilbé-2021)^. The common characteristic of these mutants is that most plants in each mutant line had a Nod^-^ phenotype similar to *ccamk-2* mutant plants^(Quilbé-2022)^, while a few plants formed one or few enlarged nodules, when inoculated with the photosynthetic *Bradyrhizobium* strain sp. ORS278, (Fig. 1a). This type of nodulation was qualified as a Big Nodule (BN) phenotype. For these mutants, Nod^-^ plants displayed typical nitrogen starvation symptoms (reduced plant growth and yellow leaves) whereas BN plants were better developed with green leaves, indicating that the formed BN nodules had nitrogen-fixing activity (Fig. 1b).

**Fig. 1.**
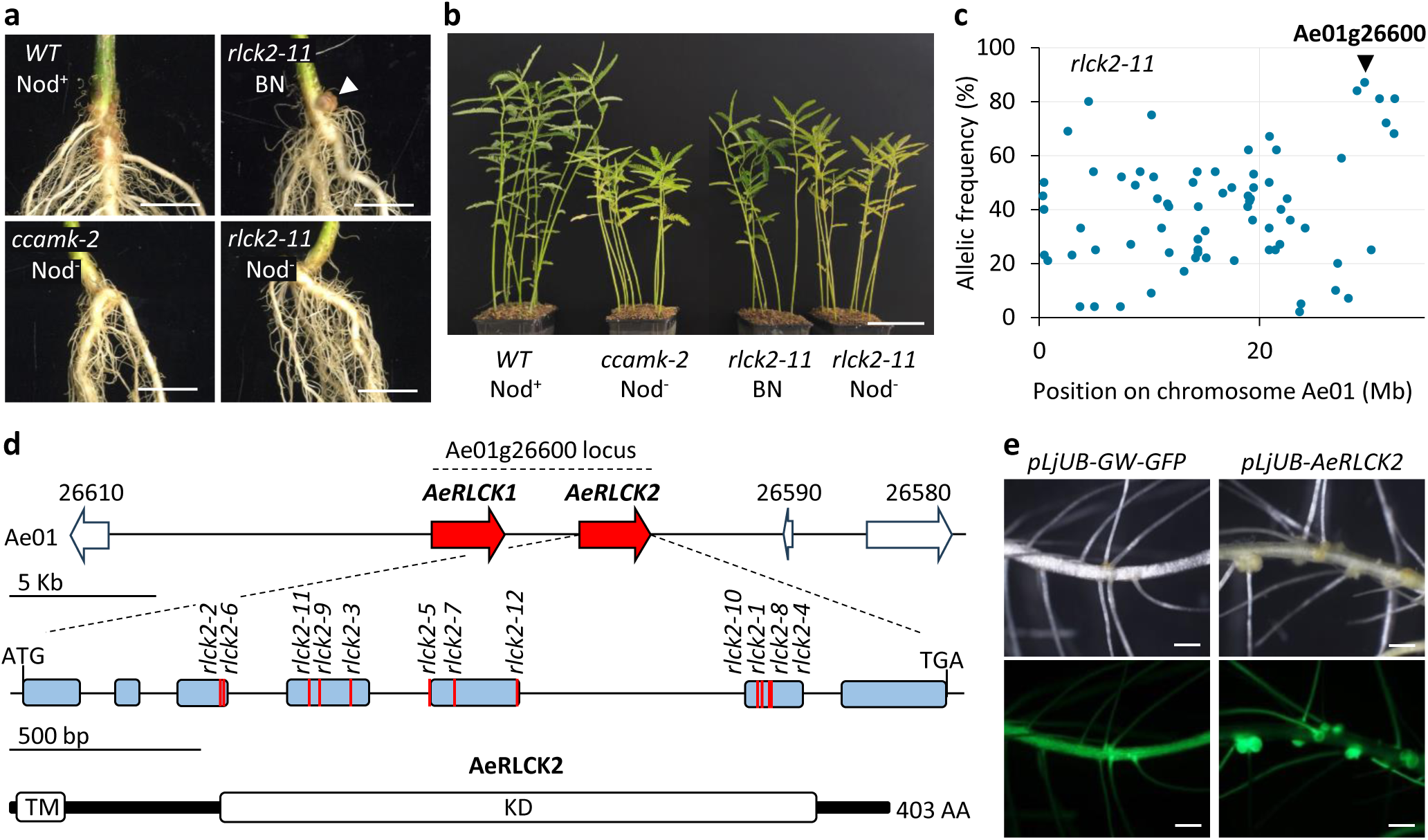
Mutant-based identification of *AeRLCK2* as required for the Nod-independent symbiosis. **a** Root phenotypes of WT, *ccamk-2* and *rlck2-11* plants at 28 days post-inoculation (dpi) with *Bradyrhizobium* ORS278 strain and grown in greenhouse conditions. Nod^+^: presence of WT nodules, Nod^-^: Absence of nodules, BN: Big Nodule (white arrowhead). Scale bar: 1 cm. **b** Aerial phenotypes of the same plants grown for 15 additional days in greenhouse conditions after analysis of their root nodulation. Scale bar: 10 cm. **c** Frequency of EMS-induced mutant alleles in pools of Nod^-^ backcrossed F2 plants derived from the *rlck2-11* mutant using mapping-by-sequencing. The SNP corresponding to the putative causal mutation in *rlck2-11* is marked with a black arrowhead. **d** *AeRLCK2* gene and protein structure. Upper panel: genomic region of chromosome Ae01 containing the Ae01g26600 locus. Red filled arrows indicate *RLCK* genes. Middle panel: gene structure of AeRLCK2. Blue boxes represent exons and red lines indicate the positions of the EMS mutations in the *rlck2* mutants. Bottom panel: domain structure of the predicted AeRLCK2 protein. White boxes indicate the positions of the predicted domains: TM for transmembrane domain and KD for kinase domain. **e** Functional complementation of *A. evenia rlck2-11.* Hairy roots of *rlck2-11* transformed with either the empty vector (left images) or containing the *AeRLCK2* CDS under the control of pLjUb (right images) at 14 dpi with *Bradyrhizobium* ORS278. GFP (Green Fluorescent Protein) was used as a plant transformation marker. Scale bar: 500µm.

Phenotypic analysis of F_2_ progeny generated from crosses between the WT line and the 12 nodulation mutants revealed that they segregated in a 3:1 ratio of plants with a WT-like nodulation phenotype (with numerous and normal sized pink nodules) to plants with either a Nod^-^ or BN phenotype, respectively (Supplementary Table 2). These data confirmed the dual mutant phenotype and demonstrated the monogenic and recessive nature of each mutation. To identify the gene(s) responsible fo this dual nodulation phenotype, we performed a mapping-by-sequencing approach on bulked F_2_ mutant plants for eight mutants. Linkage mapping repeatedly identified the same locus near the end of the Ae01 chromosome, with all mutations located in the Ae01g26600 gene (Fig. 1c, Supplementary Fig. 1 and Supplementary Table 2). Given these results, we amplified and sequenced the Ae01g26600 gene in the remaining 4 mutants and all had mutations (Supplementary Table 1). Further allelism tests between the 12 mutants showed that they belonged to the same complementation group, clearly establishing the Ae01g26600 mutations as responsible for the dual Nod^-^/BN phenotype (Supplementary Table 3).

Blast analysis revealed that Ae01g26600 codes for a RLCK. However, we observed that the coding sequence (CDS) of Ae01g26600 was twice as long as that of other *RLCK* genes. Consistent with this, *A. evenia* RNAseq datasets showed that there are actually two different *RLCK* genes at the Ae01g26600 locus^(Quilbé-2021;Chaintreuil-2016)^. This suggested that the Ae01g26600 gene is misannotated in the current *A. evenia* reference genome. We corrected this by manually delineating the two genes organized in tandem and named them *AeRLCK1* and *AeRLCK2*. As all 12 identified mutations were in the *AeRLCK2* gene, we numbered the corresponding allelic mutants *rlck2-1* to *rlck2-12* (Fig. 1d and Supplementary Table 2).

To validate our curated annotation, the *AeRLCK2* CDS was amplified from WT *A. evenia* cDNAs and cloned downstream a *L. japonicus* ubiquitin promoter (pLjUb). Using *Agrobacterium rhizogenes*-mediated hairy root transformation, the construct was introduced into the roots of the strong allele mutant *rlck2-11*, characterized by a nonsense mutation. WT-like nodulation was restored in *rlck2-11* (Fig. 1e and Supplementary Table 4). Functionnal annotation of the 403 amino acid AeRLCK2 predicted the presence of a transmembrane domain followed by a serine/threonine kinase domain, where all mutations were identified (Fig. 1d). We speculated that AeRLCK2 encodes a symbiosis plasma membrane-localized RLCK whose kinase domain integrity is essential to mediate signal transduction.

### AeRLCK2 perfoms a key function in rhizobial symbiosis

To characterize in more detail the role of *AeRLCK2* in nodulation, we performed *in vitro* growth chamber nodulation assays with *Bradyrhizobium* ORS278 on four mutant lines, *rlck2-1*, *rlck2-5*, *rlck2-10* and *rlck2-11*, using the WT line and *ccamk-2* mutant as controls. Nodulation kinetics revealed that in the WT line, nodules were mature at 10 dpi while in the *rlck2* mutants the first BN nodules emerged at 14 dpi (Supplementary Fig. 2). At 21 dpi, the nodulation frequency of the plants was 100% for WT, with each plant containing numerous nodules, and 0% for the *ccamk-2* mutant plants. In contrast, 30 to 80% of the *rlck2* mutant plants were devoid of nodules, while the others contained one or a few BNs (see below) (Fig. 2a). This points to an important role of *AeRLCK2* in nodule formation.

**Fig. 2.**
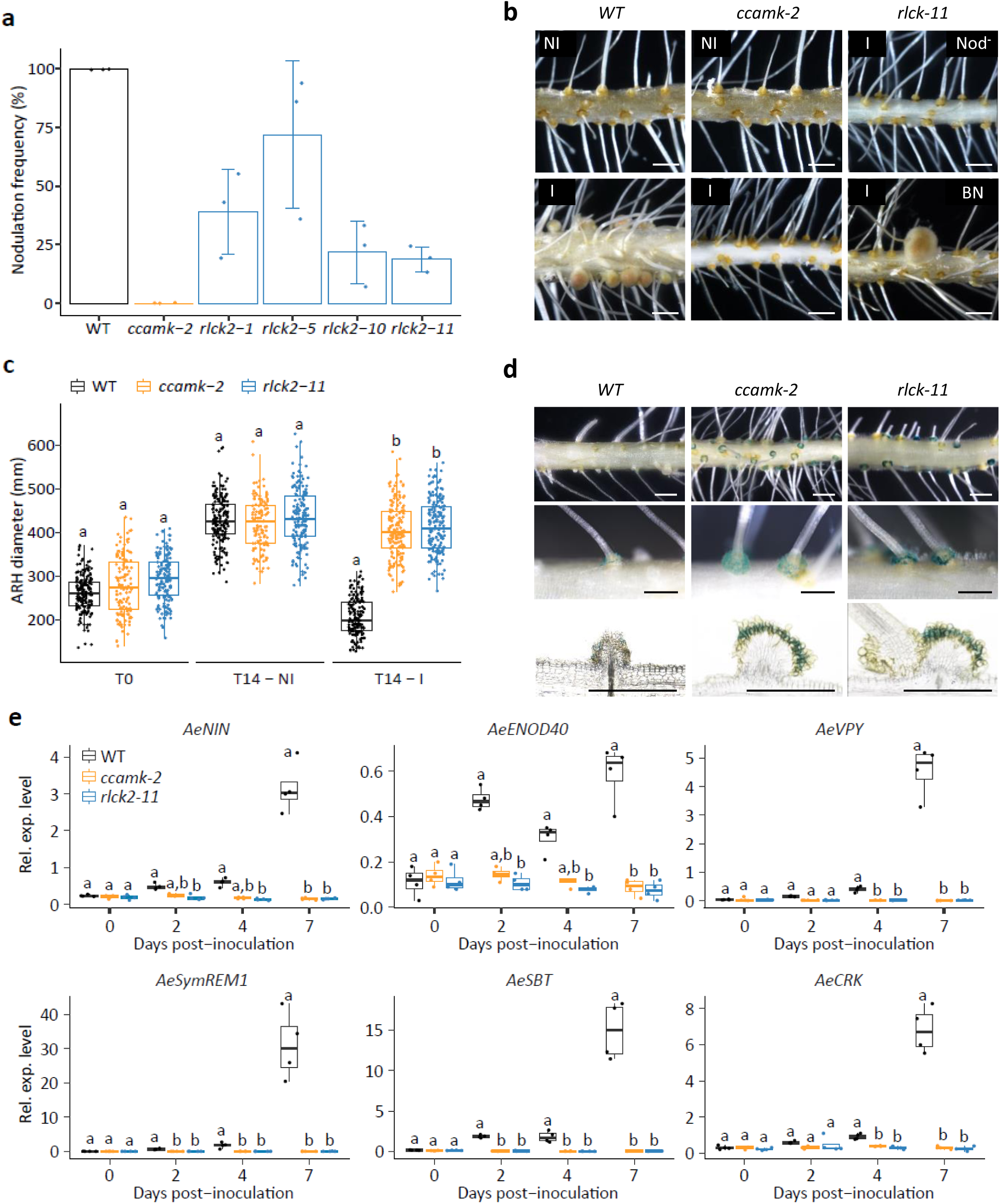
*Bradyrhizobium* infection and symbiotic signaling in *rlck2* mutants. **a** Frequency of nodule occurrence at 21 days post-inoculation (dpi) in WT, *ccamk-2*, *rlck2-1*, *rlck2-5*, *rlck2-10* and *rlck2-11* plants. **b** Comparison of root nodulation phenotypes in WT, *ccamk-2* and *rlck2-11*, under non-inoculated (NI) or *Bradyrhizobium* ORS278 inoculated (I) plants at 21 dpi. Note the presence of either a Nod^-^ or an BN phenotype in *rlck2-11* inoculated roots. Scale bar: 1mm. **c** Axillary root hair (ARH) diameter in WT, *ccamk-2* and *rlck2-11* at different time points in non-inoculated (NI) and inoculated (I) plants with *Bradyrhizobium* ORS278. T0: time 0. T14: time 14 days after inoculation or not. **d** Axillary root hair colonization of WT, *ccamk-2* and *rlck2-11* plants at 21 dpi with GUS-tagged ORS278, observed on whole roots (upper and middle panels) and root sections (lower panels). Scale bars: 1 mm (upper panels) and 0.5 mm (middle and lower panels). **e** Expression of nodulation-induced gene in WT, *ccamk-1* and *rlck2-11* plants. Relative expression levels (Rel. exp. level) of *AeNIN*, *AeSymREM1*, *AeENOD40*, *AeSBT*, *AeVPY* and *AeCRK* were measured by RT-qPCR in plant roots at 0, 2, 4 and 7 dpi. The results were normalized against *AeEF1a* and *Ubiquitin* housekeeping genes. Data presented in boxplots correspond to four biological independent replicates, with five plants per line in each replicate. Different letters indicate significant differences between conditions as determined by analysis of variance (Kruskal-Wallis) and post-hoc analysis (Dunn’s test), p<0,05.

Interestingly, inoculated roots of both Nod^-^ and BN *rlck2* plants displayed well-developped crowns of axillary root hairs (ARHs) at lateral root bases, similar to non-inoculated plants and the *ccamk-2* mutant (Nod^-^ strict), whereas on inoculated WT roots these ARHs were small (Figure 2b). In *A. evenia*, these ARHs are the first colonisation sites of bradyrhizobia and their development is tightly controlled by the nitrogen status of the plant^(Quilbé-2022)^. Kinetics of ARH development showed that they develop over time in uninoculated WT plants, whereas their development is suppressed in inoculated WT plants (Fig. 2c). No such *Bradyrhizobium*-induced repression of ARH development was observed in *rlck2* and *ccamk-2* mutants. This observation led us to analyse at which stage infection is blocked in *rlck2* mutants. X-Gluc staining of ORS278-GUS inoculated *rlck2-11* mutant roots at 21 dpi revealed intense blue staining on the surface and between the ARHs, but no penetration into the inner root cortex was observed (except for the BN), similar to the *ccamk-2* mutant (Fig. 2d). In contrast, *crk* mutants, which develop small bumps containing infection pockets after inoculation^(Quilbé-2022)^, showed reduced ARH development in the presence of ORS278 (Supplementary Fig. 3). RT-qPCR analysis of six symbiosis-induced marker genes, *AeNIN*, *AeVPY*, *AeSymREM1*, *AeENOD40*, *AeSBT* and *AeCR*K^(Quilbé-2022)^, showed no induction of their expression in *rlck2-11* and *ccamk-2* inoculated with the ORS278 strain (Fig. 2e). Taken together, these results indicate that the mutations in *AeRLCK2* block early symbiosis responses.

Additionally, *rlck2* mutant plants showed a drastic decrease in nodule numbers. WT plants contained an average of 35 nodules per root at 21 dpi, while roots of most nodulated *rlck2* mutant plants exhibited only one or two BN nodules (Fig. 3a). These BNs had an average diameter that was twice as large as that of WT nodules (Fig. 3b). We interpret this as a compensatory mechanism for their very few numbers in *rlck2* mutants. Roots of *rlck2* mutant plants containing BN nodules had nitrogenase enzyme activity, as measured by the acetylene reduction assay (ARA), and accordingly these plants carried green leaves (Fig. 3c, Supplementary Fig. 4). Light microscopy analysis of BN nodule sections from *rlck2-11* plants showed that they had a similar structure to WT nodules, with a central tissue infected by bacteria and a peripheral uninfected nodule cortex containing vascular bundles (Fig. 3d). However, very often these BN nodules also contained small necrotic zones and brown spots. These areas had an intense yellow/red fluorescent appearance when a FITC filter was used, suggesting the presence of polyphenolic compounds. Confocal microscopic analysis of the same nodule sections showed that, in contrast to WT nodules, BN nodules contained unevenly infected plant cells. In general, the infected plant cells of BN nodules contained spherical bacteroids, but in some cases, rod-shaped undifferentiated bacteria were also observed (Fig. 3e). These observations highlight an important role of *AeRLCK2* in nodule infection and bacterial differentiation.

**Fig. 3.**
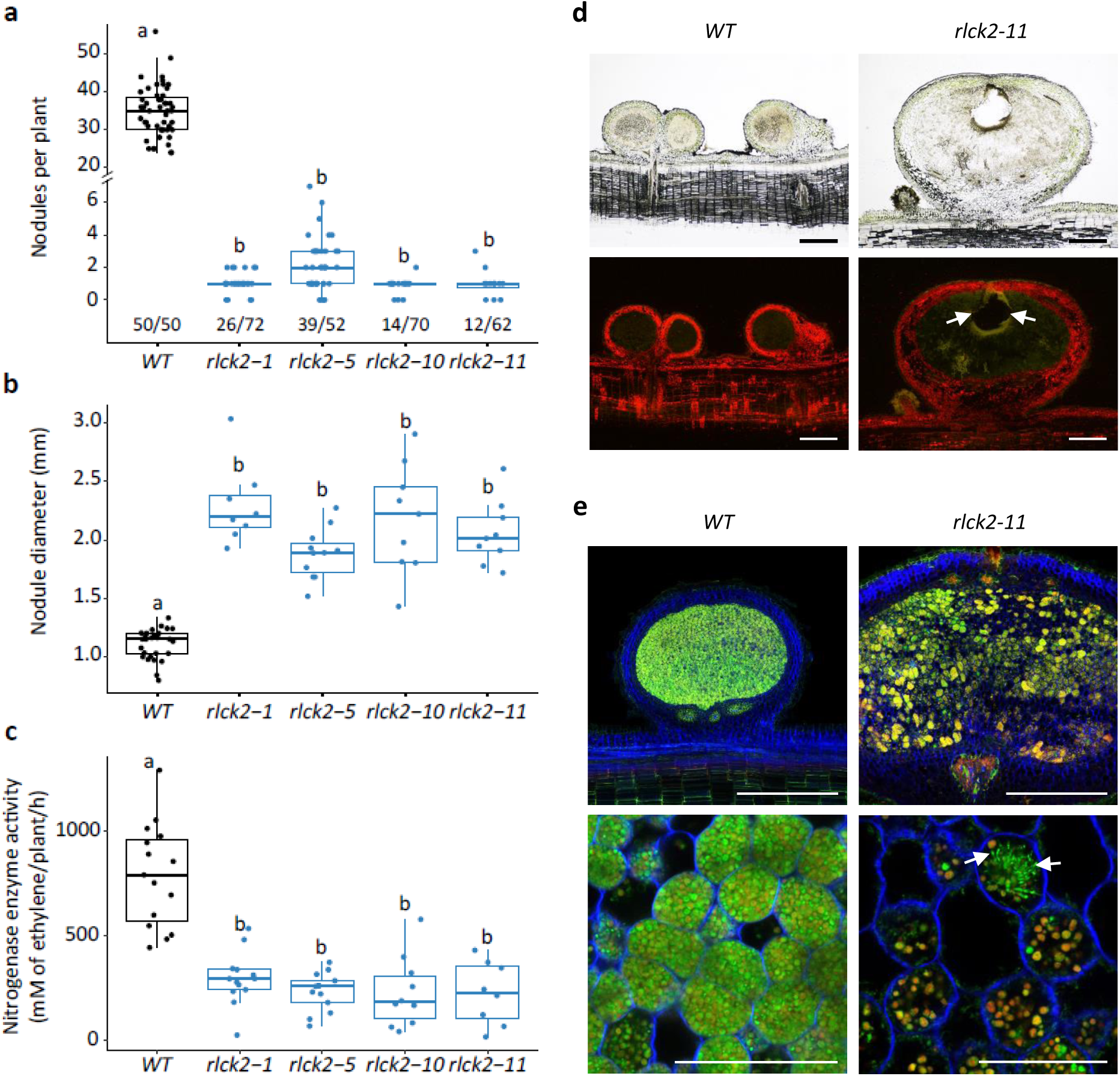
Nodule development and colonisation by *Bradyrhizobium* in *rlck2* mutants. **a** Number of pink nodules formed on nodulated plants in WT, *rlck2-1, 5, 10* and *11* plants at 21 days post-inoculation (dpi) with *Bradyrhizobium* ORS278. Numbers below the boxplots indicate the number of nodulated plants relative to the total number of inoculated plants. **b** Nodule diameter and **c** nitrogenase enzyme activity measured by acetylene reduction assay (ARA) (n≥3 nodulated root per line and biological replicate) from the same plants as in (**a**). Data in (**a**) to (**c**) correspond to 3 biological independent replicates. Letters indicate significant differences between conditions, as determined by analysis of variance (Kruskal-Wallis) and post-hoc analysis (Dunn’s test), p<0,05. **d** Cross-sections of WT and *rlck2-11* nodules observed under brightfield (top) or FITC filter (bottom). White arrows indicate the occurrence of defense-like responses within the nodule. Scale bar: 500 μm. **e** Cytological analysis of nodule cross-sections from WT and *rlck2-11* plants using a confocal microscope after staining with SYTO9 (green, live bacteria), propidium iodide (red, infected plant nuclei and dead bacteria or bacteria with a compromised membrane) and calcofluor (blue, plant cell wall). White arrows show elongated bacteria. Scale bars: 500 μm (top), 50 μm (bottom).

### AeRLCK2 physically interacts with and is phosphorylated by AeCRK

Recently, we showed that a Cysteine-rich RLK-coding gene, AeCRK, is essential for the establishment of the N-fixing symbiosis in *A. evenia* ^(Quilbé-2021,2022)^. Since RLCKs are known to interact with RLKs to mediate downstream signaling^(Liang-Zhou-2018;Lin-2013)^, we investigated the hypothesis that AeRLCK2 and AeCRK form a plasma membrane-bound complex. For this, we first examined the subcellular localisation of AeCRK and AeRLCK2 in *Nicotiana benthamiana* leaves, by generating a translational fusion with the Yellow Fluorescent Protein (YFP). Transient overexpression of AeCRK-YFP induced cell death in *N. benthamiana* leaves five days after *Agrobacterium tumefaciens*-mediated transformation. In contrast, an engineered dead-kinase version (AeCRK^G359E^) with a mutation in the glycine-rich loop did not trigger cell death and was therefore used (Supplementary Fig. 5). The YFP fusion constructs were transiently expressed in *N. benthamiana* leaves in combination with the plasma membrane marker MtLYK3^(Klaus-Heisen-2011)^ fused with Cyan Fluorescent Protein (CFP) (Fig. 4a). AeCRK^G359E^ and AeRLCK2 co-localized with MtLYK3, confirming their targeting to the plasma membrane. Since, AeRLCK2 is atypical in harboring a predicted transmembrane domain (TM), we also tested an N-terminal truncated version of AeRLCK2 (AeRLCK2^ΔTM^-YFP). For this latter, the signal was observed in the nucleus and cytoplasmic threads, indicating that the predicted TM is important for the protein anchoring to the plasma membrane (Fig. 4a).

**Fig. 4.**
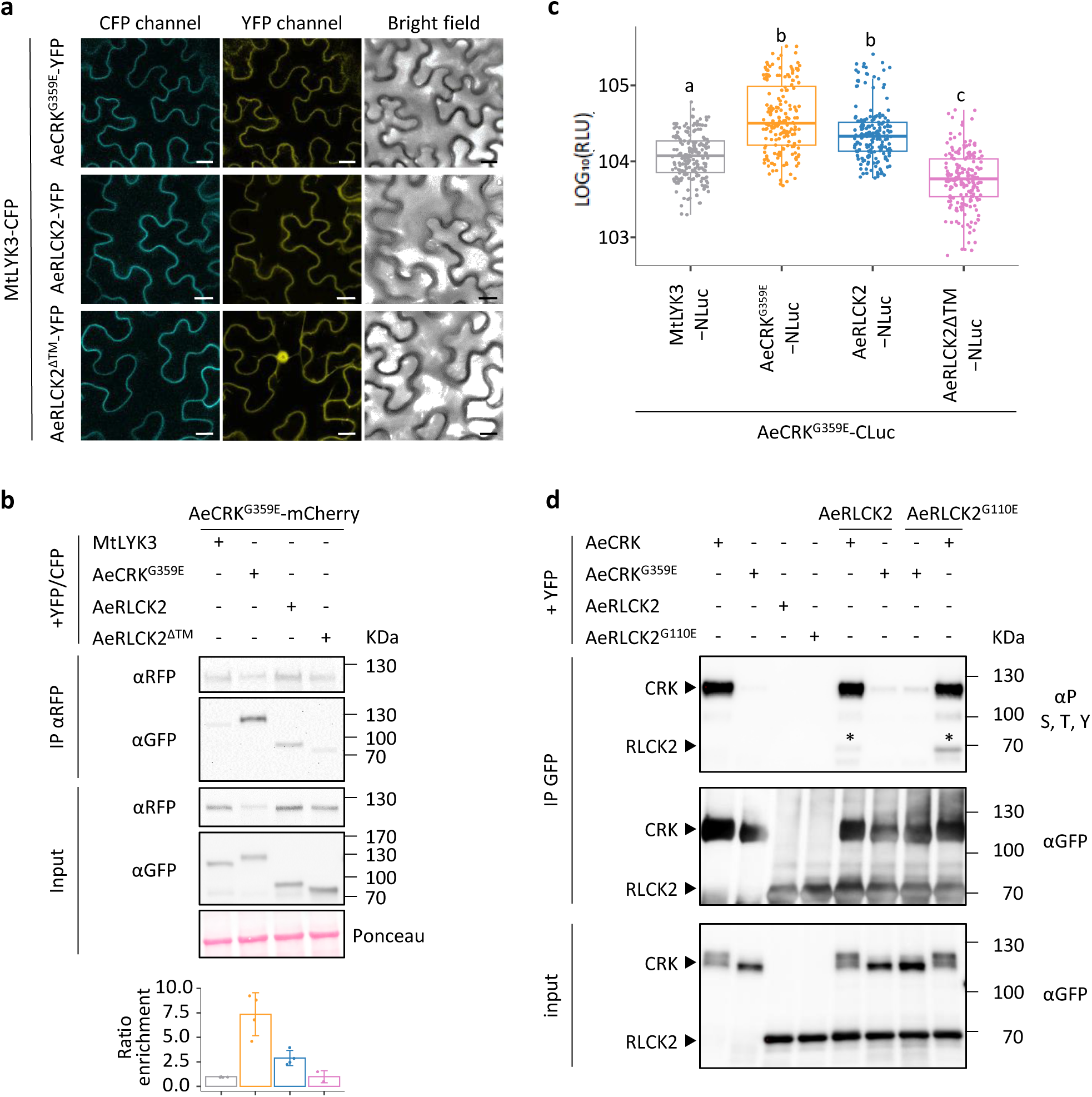
AeRLCK2-AeCRK interaction and kinase assays. **a** Confocal microscopy observations of *Nicotiana benthamiana* leaf cells showing plasma membrane localisation of AeCRK^G359E^-YFP, AeRLCK2-YFP, and nucleo-cytoplasmic distribution of the truncated transmembrane version of RLCK2 (AeRLCK2^ΔTM^). MtLYK3-CFP was used as a plasma membrane marker. Scale bar: 20 µm. **b** Co-immunoprecipitation assay showing interaction of AeCRK^G359E^-mCherry with AeCRK^G359E^-YFP and AeRLCK2-YFP. Proteins were immunoprecipitated with αRFP magnetic agarose beads and co-purified proteins were detected with αGFP antibodies (upper panel). Input (middle panel) and band intensities were calculated and normalized to the negative control MtLYK3 (bottom panel, ranging from 2 to 4 biological replicates). Ponceau staining was used as loading control. **c** Split-luciferase assays showing the interaction of AeCRK^G359E^-CLuc with AeCRK^G359E^-NLuc or AeRLCK2-NLuc. Boxplots represent bioluminescence intensity from seven independent replicates. Expression levels of 3Flag-CLuc and 3HA-NLuc fusions were assessed by Western blot (Supplementary Fig. 6). Bioluminescence intensities were normalized to protein expression and data were Log-transformed (Log_10_). Letters indicate significant differences between samples, as determined by analysis of variance (Kruskal-Wallins) and post-hoc analysis (Dunn’s test), p<0,05. RLU: Relative luminescence unit. **d** Kinase activity assay showing transphosphorylation of AeRLCK2 by AeCRK in *Nicotiana benthamiana* leaf cells. Full-length YFP-tagged proteins were immunoprecipitated with αGFP magnetic agarose beads. Phosphorylation status was analyzed after SDS-PAGE and detected with anti-S, -T and -Y antibodies. Asterisks indicate the phosphorylation status of AeRLCK2-YFP (top). Input (bottom).

Next, we tested AeCRK-AeRLCK2 interactions by co-immunopurification (IP) assays. The mCherry-tagged AeCRK^G359E^ was co-expressed with YFP-tagged AeCRK^G359E^, RLCK2, RLCK2^ΔTM^ and the negative control MtLYK3-CFP in *N. benthamiana* leaves. After IP of AeCRK^G359E^, the co-purified proteins were detected with αGFP antibodies (Fig. 4b). AeCRK^G359E^-YFP and AeRLCK2-YFP were enriched 7.37-fold and 2.9-fold, respectively, compared to the negative control MtLYK3-CFP (Fig. 4b). Conversely, no significant enrichment was observed for AeRLCK2^ΔTM^-YFP (Fig. 4b). To assess these pairwise interactions, we performed split-luciferase assays by fusing AeCRK^G359E^ with the C-terminal part of the luciferase (3Flag-CLuc) and the potential interactors with the N-terminal part of the luciferase (3HA-NLuc). Combination of -NLuc and -CLuc fusion proteins were co-expressed in *N. benthamiana* leaves. The corresponding bioluminescence was measured following luciferin infiltration, and the data were normalized by the expression level of the -NLuc fusion proteins (Supplementary Fig. 6). Co-expression of AeCRK^G359E^-CLuc with AeCRK^G359E^-NLuc or AeRLCK2-NLuc, but not with RLCK2^ΔTM^-NLuc, resulted in significantly higher bioluminescence intensities compared to MtLYK3-NLuc (Fig. 4c and Supplementary Fig. 6). Thus, these findings consistently showed that AeCRK can form homodimers and physically interact with AeRLCK2. The N-terminal domain of AeRLCK2, which confers plasma membrane localisation, is essential for this interaction.

AeCRK and AeRLCK2 have typical Ser/Thr kinase domains, suggesting that their interaction may involve phosphorylation events. We therefore investigated their kinase activities. The kinase domains (KD) of AeCRK and AeRLCK2 were translationally fused to a GST-tag and expressed in *Escherichia coli*. After purification, their autophosphorylation activity was studied *in vitro* using radiolabelled ATP (^32^P-ATP). Autoradiography revealed a robust autophosphorylation activity for AeCRK^KD^, whereas AeRLCK2^KD^ was much less efficient (Supplementary Fig. 7). The AeRLCK2^KD-G110E^ mutant, containing a mutation in the glycine-rich loop, found in *rlck2-*6, failed to incorporate ^32^P-ATP indicating a lack of kinase activity. Transphosphorylation studies showed that AeCRK^KD^ phosphorylated the dead kinase AeRLCK2^KD-G110E^ but not free GST (Supplementary Fig. 7). These results were confirmed by *in planta* experiments with the full-length YFP-tagged proteins transiently expressed in *N. benthamiana* leaves. After IP, the phosphorylation status of the proteins was assessed using an antibody that recognizes phosphorylated threonine, tyrosine and serine residues (Fig. 4d). AeCRK was highly phosphorylated whereas AeCRK^G359E^ showed either no or low levels of phosphorylation. Surprisingly, no phosphorylation was observed for AeRLCK2 in *planta*. To investigate the possibility of trans-phosphorylation, AeCRK-YFP was co-expressed with AeRLCK2-YFP or AeRLCK2^G110E^-YFP. In both cases, AeRLCK2 was phosphorylated, whereas this was not observed when using the AeCRK^G359E^ version (Fig. 4d). Finally, the phosphorylation sites of AeRLCK2 targeted by the kinase activity of AeCRK were searched by comparative LC-MS/MS analysis of *in planta* immunopurified AeRLCK2 produced alone or together with AeCRK or AeCRK^G359E^. Five phosphorylation sites, corresponding to three threonines (T66, T90 and T106) and two serines (S133 and S321) were specifically identified in the kinase domain of AeRLCK2 in the presence of AeCRK (Supplementary Fig. 8 and 9). Consistently, AeCRK transphosphorylation of AeRLCK2 was also found using antibodies that recognize only phosphorylated threonine residues (Supplementary Fig. 8). Taken together, these results demonstrated that AeCRK and AeRLCK2 have distinct kinase activities and that AeCRK transphosphorylate AeRLCK2 on specific residues.

### *AeRLCK2* arose from a duplication of a mycorrhiza-conserved gene in *Aeschynomene*

Since *AeRLCK2* is tandemly organized with *AeRLCK1* in the *A. evenia* genome, we investigated whether this *RLCK* gene tandem is present in other legumes. Synteny analysis based on genome sequence comparisons revealed the presence of a single *RLCK* homolog at the same locus in the analyzed legume species (Supplementary Fig. 10). To specify the relationships between *AeRLCK1*, *AeRLCK2* and *RLCK* homologs, we analysed the genome sequences of 15 legume species (13 Papilionoideae and 2 Caesalpinoideae) and 3 non-legume species (Supplementary Data 1). We also included in this analysis unpublished RNAseq data from *Aeschynomene afraspera*, a close relative of *A. evenia* that uses a Nod-dependent symbiosis^(Bonaldi-2011)^ (Supplementary Table 5). This search retrieved *RLCK* homologs for each analysed plant species except *Arabidopsis thaliana* and lupin sp, two species unable to form AM, consistently with previous phylogenomic studies that predicted their conservation for AM (Supplementary Table 6)^(Bravo,2016)^. In rice and *L. japonicus*, the homologous genes *OsRLCK171*, *LjAMK8* and *LjAMK24* were demonstrated to be essential for AM ^(Leng-2023)^. Phylogenetic reconstruction, based on protein sequences, placed the *RLCK* genes present in non-legume and Caesalpinoideae species, such as *OsRLCK171*, in separate clades and revealed that the *RLCK* homologs present in Papilionoideae legume species were distributed in two sister clades, one containing *LjAMK8* and the other *LjAMK24* (Fig. 5a). These clades probably originated from the ancient whole genome duplication in the Papilionoideae subfamily. In *A. afraspera*, *AaRLCK_O* and *AaRLCK_P* also corresponds to the paralogous Papilionoid gene pair. In contrast, the two *A. evenia RLCK* genes, *AeRLCK1* and *AeRLCK2*, clustered together in the clade harbouring *LjAMK8* and *AaRLCK_O,* whereas the expected paralog was missing (Fig. 5a). This suggests a recent *AeRLCK1*-*AeRLCK2* duplication accompanied by the loss of the paralogous gene in *A. evenia*.

**Fig. 5.**
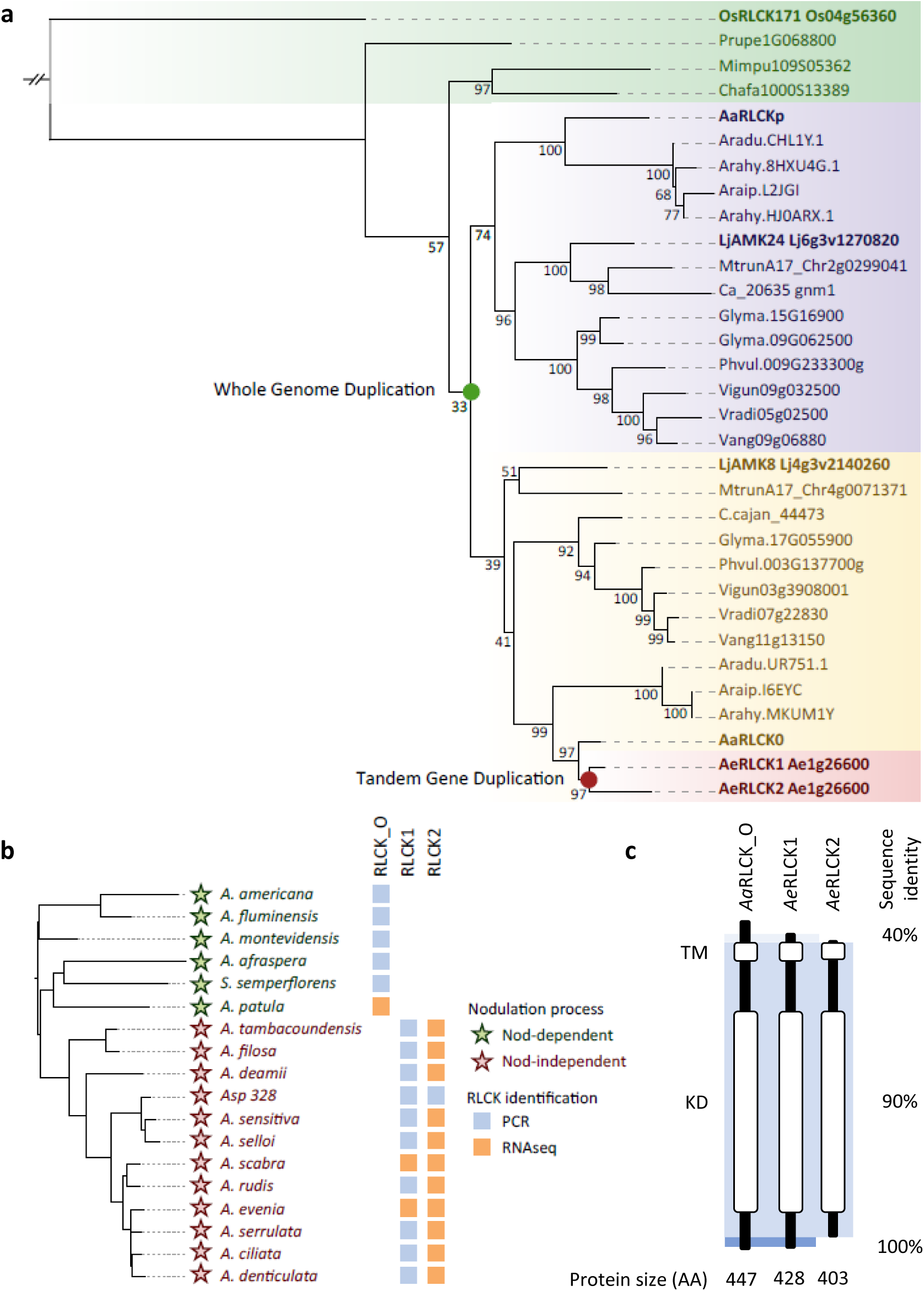
Phylogeny of legume *RLCK* genes and evolution in *Aeschynomene* species. **a** Maximum likelihood (ML) phylogenetic reconstruction of the orthogroup containing AeRLCK2. Color coding indicates non-papilionoid RLCKs (green), the two papilionoid RLCK clades (purple and yellow) putatively originating from the 58-MA Whole Genome Duplication (WGD) event (green dot), and the two RLCK copies present in *A. evenia* (red), which are derived from a recent tandem duplication (red dot). **b** Detection of different *RLCK* gene versions in *Aeschynomene* species and the closely related species *Soemmeringia semperflorens.* The ML phylogenetic tree was constructed using concatenated *ITS* (Internal Transcribed Spacer) and *matK* sequences. Green stars indicate a Nod-dependent symbiosis and red stars indicate a Nod-independent symbiosis. The RLCK_O, RLCK1 and/or RLCK2 copies were identified in available RNAseq data (orange square) and by PCR amplification on genomic DNA (blue square). (**a**) and (**b**) support values were determined using 100,000 iterations of the ultrafast bootstraps approximation (UFboot). **c** Domain structure of *Aa*RLCK_O, *Ae*RLCK1 and *Ae*RLCK2 and sequence similarities between the proteins. White bars indicate predicted domains. TM: transmembrane domain, KD: kinase domain, AA: amino acids. Intensities of blue shaded backgrounds delineate zones with different level of sequence identity. All domains are to scale.

To clarify when these gene changes occurred in *Aeschynomene*, we searched for *RLCK* orthologs among RNAseq data previously generated from roots and nodules for 11 *Aeschynomene* species in the Nod-independent clade^(Quilbé-2021)^. We found an *AeRLCK2* ortholog for each of these species and an *AeRLCK1* ortholog only for *A. scabra,* but no putative *RLCK* paralog was detected (Fig. 5b, Supplementary Data 2). We completed this analysis by experimental investigation of their presence in *Aeschynomene* species^(Brottier-2018)^. To this end, we designed primers matching conserved or specific regions to *AaRLCK_O*, *AeRLCK1* or *AeRLCK2* copies and screened by PCR amplification followed by amplicon sequencing in *Aeschynomene* species and the allied species, *Soemmeringia semperflorens*. Sequences similar to *AeRLCK1* and *AeRLCK2* were identified in the 11 Nod-independent *Aeschynomene* species as for *A. evenia*, whereas a single *RLCK* sequence was recovered in the 5 Nod-dependent species as in *A. afraspera* (Fig. 5b). Thus, there is a perfect correlation between the Nod-independent symbiosis and the *RLCK* gene duplication in *Aeschynomene* legumes. Interestingly, no additional gene tandems or clusters specific to the Nod-independent *Aeschynomene* lineage could be evidenced in the set of 138 genes predicted to be required for AM (Supplementary Table 6)^(Bravo,2016)^.

To substantiate the changes associated with *AeRLCK2*, we compared the type of *RLCK* genes present in *Aeschynomene* species. Protein sequence alignment and 3D modelling showed that they have the same general structure and share a highly conserved sequence (AeRLCK2 has 86% and 80% amino acid identity with AeRLCK1 and AaRLCK_O, respectively) (Fig. 5c and Supplementary Fig. 11). However, RLCK2 proteins are shorter at both the N- and C-terminus compared to the other RLCK proteins. We next assessed the extent of gene structural variation using *AaRLCK_O* cDNA and *AeRLCK1*/*AeRLCK2* genomic sequences. Significant differences were observed in their 5’- and 3’-UTR regions as well as in their flanking exons (Supplementary Fig. 12). We also identified a sequence in the promoter region of *AeRLCK2* that is highly similar to a downstream gene, Ae01g26580 (Supplementary Fig. 13). Our interpretation is that both the ancestral *RLCK* and the downstream genes underwent a gene tandem duplication in the ancestor of the Nod-independent *Aeschynomene* species. Subsequently, complex rearrangements occurred in the promoter region and gene extremities of the *RLCK2* copy (Supplementary Fig. 13). These data support the hypothesis that the creation of *AeRLCK2* has participated in enabling the evolution of the Nod-independent symbiosis.

### AeRLCK2 is dispensable for arbuscular mycorrhiza

To determine whether *AeRLCK2* is important for AM, as its homologs in rice (*OsRLCK171*) and *L. japonicus* (*LjAMK8* and Lj*AMK28*)^(Leng-2023)^, we tested four *rlck2* mutants together with WT plants and the *ccamk-2* mutant line. In contrast to the completely mycorrhiza-free *ccamk-2* mutant, roots of both WT and the four *rlck2* mutants contained fungal hyphae, arbuscules and vesicles, 6 weeks after inoculation with *Rhizophagus irregularis* spores (Fig. 6a). Quantitative assessement of mycorrhization levels using the Trouvelot method further showed that the *rlck2* mutants were colonized similary to the WT plants (Fig. 6b, Supplementary Fig. 14).

**Fig. 6.**
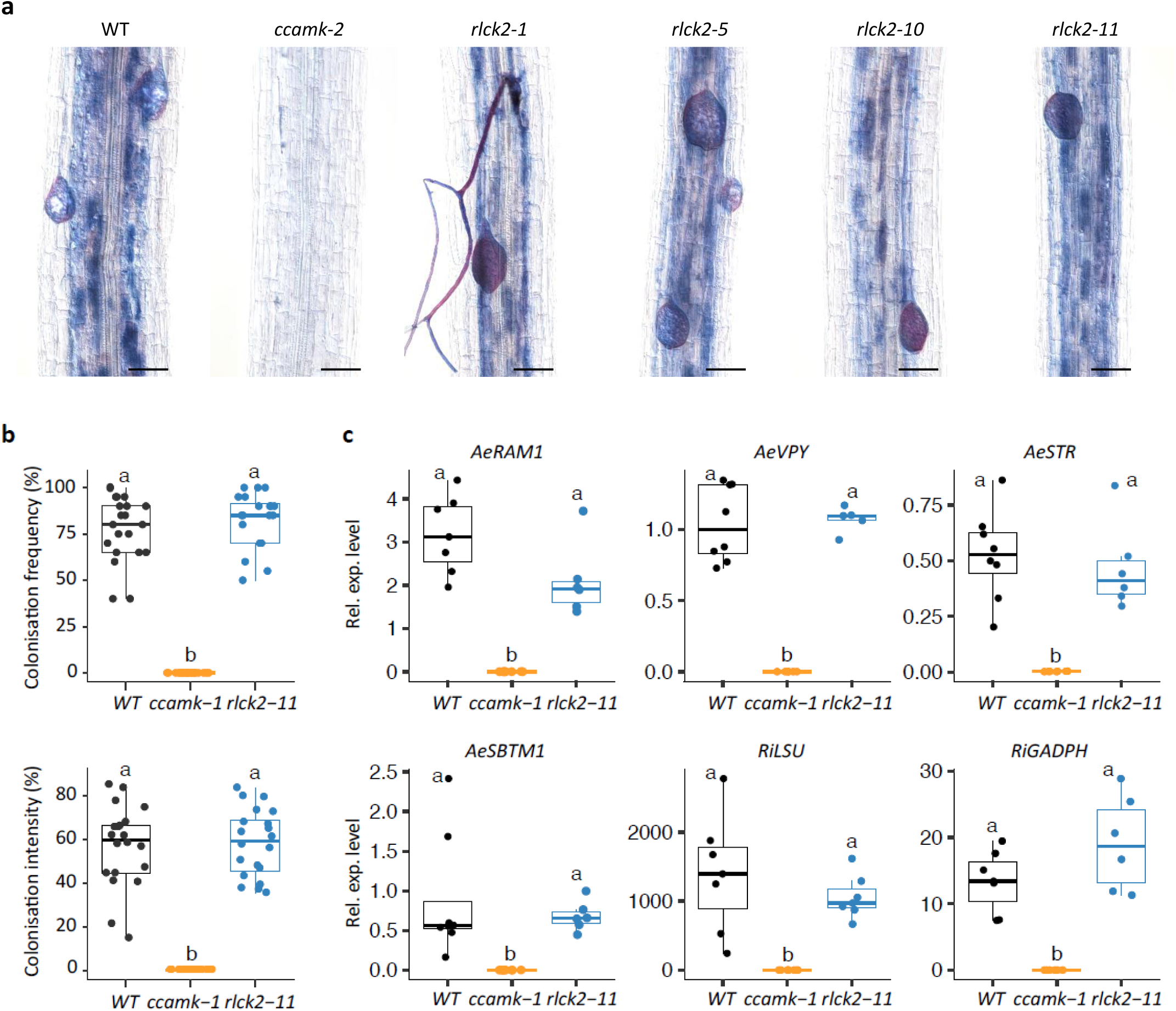
Arbuscular mycorrhizal (AM) root colonization in *rlck2* mutants. **a** Microscopy images of *R. irregularis* colonization of WT, *ccamk-2* and *rlck2* mutants at 6 weeks post-inoculation (wpi), stained with Sheaffer skrip ink. Scale bars: 50 µm. **b** Box plots show the colonisation frequency and intensity, both expressed as percentages, in 6 wpi WT, *ccamk-2* and *rlck2-11* plants. **c** Analysis of AM-induced gene expression in WT, *ccamk-2* and *rlck2-11* plants. Relative expression levels (Rel. exp. level) of plant *AeRAM1*, *AeVPY*, *AeSTR*, *AeSBTM1* and fungal *RiLSU*, *RiGADPH* genes were measured by RT-qPCR in roots of 6 wpi plants. The results were normalized against *AeEF1a* and *Ubiquitin*. The data represent four biological replicates, with five plants per line in each replicate. Letters indicate significant differences between lines, as determined by analysis of variance (Kruskal-Wallis) and post-hoc analysis (Dunn’s test), p<0,05.

To deepen the analysis, we focused on *rlck2-11*. At 6 wpi, there was again no difference in both the frequency and intensity of colonisation between the *rlck2-11* mutant and the WT line (Fig. 6b). In parallel, quantification of the fungal *RiLSU* and *RiGADPH* gene expressions, as markers of fungal biomass in the root tissues was performed by RT-qPCR analysis^(Quilbé-2022)^. Expression levels of these fungal genes in *rlck2-11* roots were equivalent to those in WT roots, indicating that *R. irregularis* colonisation in *A. evenia* roots is not affected by mutation in *AeRLCK2* (Fig. 6e). We also determined the expression level of plant AM-induced genes, *AeRAM1*, *AeVPY*, *AeSTR* and *AeSBTM1* by RT-qPCR analysis^(Quilbé-2022)^ in *rlck2-11*. In this mutant, the induction levels of all genes tested were similar to those in WT plants (Fig. 6e). Therefore, the *rlck2* mutants appeared to develop functional AM.

The lack of any detectable mycorrhizal phenotype could either reflect an absence of involvement in AM of *AeRLCK2* or a potential functional redundancy with *AeRLCK1* in the AM symbiosis. As *LjAMK8*, *LjAMK24* and the functionally related *LjKIN3* are AM-induced genes in *L. japonicus*^(Leng-2023)^, we investigated whether such an upregulation of expression occurs for *A. evenia* homologs. To enable fine expression analysis, we generated RNaseq data for WT *A. evenia* inoculated or not with *R. irregularis* (Supplementary Table 7). As expected^(Quilbé-2022)^, the AM-marker genes *AeRAM1*, *AeVPY*, *AeSTR* and *AeSBTM1* were well induced during mycorrhization (Supplementary Fig. 15). We observed a strong induction during AM for *AeKIN3* (Ae06g09820), the putative ortholog of *LjKIN3*, whereas the induction level of *AeRLCK1* was weaker and expression of *AeRLCK2* itself appeared to be unaffected (Supplementary Fig. 15).

### AeRLCK2 shows adaptations to the Nod-independent symbiosis

To question how the Nod-independent *Aeschynomene*-specific *RLCK* gene duplication may have led to the involvement of *AeRLCK2* in nodulation, we compared the functionality of *AeRLCK2* with *AeRLCK1*, the duplicated gene in *A. evenia,* and with *AaRLCK_O*, the corresponding single copy gene in *A. afraspera*. First, the three *RLCK* genes fused to *YFP* were expressed in *N. benthamiana* leaves. In contrast to *AeRLCK2*, cell death was observed in leaves expressing *AaRLCK_O* and *AeRLCK1*, at 8 dpi (Supplementary Fig. 16a). Confocal microsocpic analysis and kinase assays performed at 3 dpi, i.e. before the onset of cell death, showed that both AaRLCK_O and AeRLCK1 proteins were localized at the cell periphery but showed no autophosphorylation activity, as for AeRLCK2 (Supplementary Fig. 16b,c).

To further test whether AaRLCK_O, AeRLCK1 and AeRLCK2 are functionally equivalent, we performed cross-complementation studies. We used the *AeRLCK2* upstream region (∼ 2.5 kb including the 5’-UTR) to drive the expression of the *AaRLCK_O*, *AeRLCK1* and *AeRLCK2* CDS in the *rlck2-11* mutant line. Using *A. rhizogenes* root transformation and *Bradyrizobium* ORS278 inoculation, we found full complementation of the *rlck2-11* mutant phenotype with *AeRLCK2* at 3 wpi, both in terms of aerial plant development and nodule number, validating the functionality of the *AeRLCK2* promoter (Fig. 7a,b; Supplementary Table 8). Unexpectedly, in contrast to *AeRLCK1*, *AaRLCK_O* was also able to rescue the *rlck2-11* mutant phenotype (Fig. 7a,b; Supplementary Table 6). Microscopy analysis of *AaRLCK_O* and *AeRLCK2* complemented roots revealed WT sized nodules with cells well-filled with bacteria (Fig. 7c). Similar results were also obtained when expressing the three *RLCK* genes under the constitutive Ubiquitin promoter in *rlck2-11* (Supplementary Fig. 17, Supplementary Table 8).

**Fig. 7.**
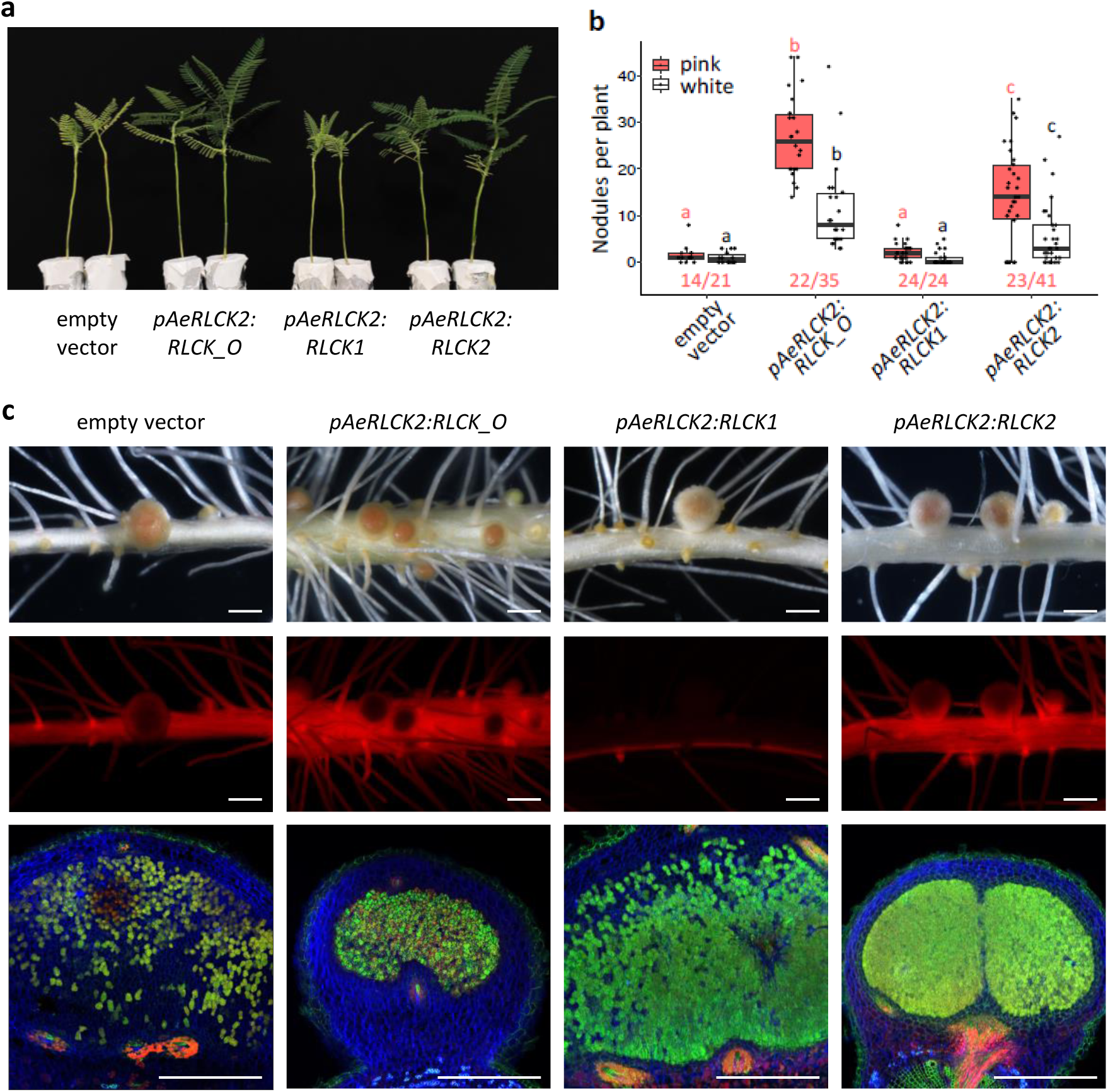
*A. evenia rlck2* mutant cross-complementation of root nodulation. Hairy roots of *A. evenia rlck2-11* plants were transformed with the empty vector (EV) containing the DsRed marker, or the same vector containing *pAeRLCK2:RLCK0*, *pAeRLCK2:RLCK1* or *pAeRLCK2:RLCK2* and their nodulation phenotype was evaluated 21 days post-inoculation with *Bradyrhizobium* ORS278. Observations were made on two biological replicates. Representative root nodulation phenotypes are shown here and detailed in Supplementary Table 8. **a** Plant aerial phenotype. **b** Number of pink and white nodules formed on plants expressing the indicated constructs. Dots represent individual plants. Red numbers below the boxplots indicate the number of plants with pink nodules, relative to the total number of transformed plants. Letters indicate significant differences between constructs, as determined by analysis of variance (Kruskal-Wallis) and post-hoc analysis (Dunn’s test), p<0,05. **c** Nodule analysis on *rlck2-11* roots transformed with the indicated constructs. Top and middle panels: microscopy observations of whole nodules under brightfiled and red fluorescence using a DsRed filter, respectively. Bottom panels: cross-sections of nodules stained with SYTO 9, propidium iodide and calcofluor, and observed with a confocal microscope. Scale bars: 1 mm (top and middle panels), 0.5 mm (bottom panel).

We next investigated whether AeRLCK2 behaves similarly to other related *RLCK* genes at the transcriptional level. Based on RNAseq data during nodulation that are available for *A. evenia*^(Quilbé-2021;Gully-2018)^ and obtained in the present study for *A. afraspera* (Supplementary Table 4). *AeRLCK1* and *AeRLCK2* were found to be expressed in roots and nodules but *AeRLCK2* was at least 10 folds more expressed that *AeRLCK1* in both organs (Fig. 8a). In contrast, *AaRLCK_O* and *AaRLCK_P* showed clear induction of their expression level during nodulation (Fig. 8a). This behaviour is similar to that described in *L. japonicus* for their respective orthologs, *LjAMK8* and *LjAMK24*^(Leng-2023)^. We also analyzed the expression levels of *AeCRK* and its ortholog *AaCRK*, identified by BLAST search in the *A. afraspera* transcriptome. Expression of both *CRK* genes was found to be induced during nodulation (Fig. 8a). To better understand these contrasting expression behaviours, we monitored the spatio-temporal expression profile of *AeRLCK2* and *AeCRK* in WT roots transformed by *A. rhizogenes* with promoter-GUS fusions (Fig. 8b,c). For *AeRLCK2*, a weak GUS staining was detected at the base of lateral roots before inoculation. After inoculation with *Bradyrhizobium* ORS278, increased GUS staining was observed at the base of lateral roots and in nodule primordia (2 and 4 dpi). When nodules emerged from the lateral root base (7 dpi), GUS staining was predominant at the nodule base and the vascular bundles of the adjacent lateral root. Finally, in mature nodules (14 dpi), GUS staining persisted at the nodule base and and in the cell layers surrounding the central nitrogen-fixation zone. For *AeCRK*, no expression was detected before inoculation. At early stages of the interaction *AeCRK* expression mimicked that of *AeRLCK2* in nodule primordia (4 dpi). But then, the expression of *AeCRK* was observed in the central infected tissue of mature nodules (7 and 14 dpi. It is noteworthy that in *L. japonicus*, *LjAMK8* and *LjAMK24*, are expressed in the central infected tissue of mature nodules^(Leng-2023)^. These observations support the distinctness of the *AeRLCK2* expression pattern observed in *A. evenia*.

**Fig. 8.**
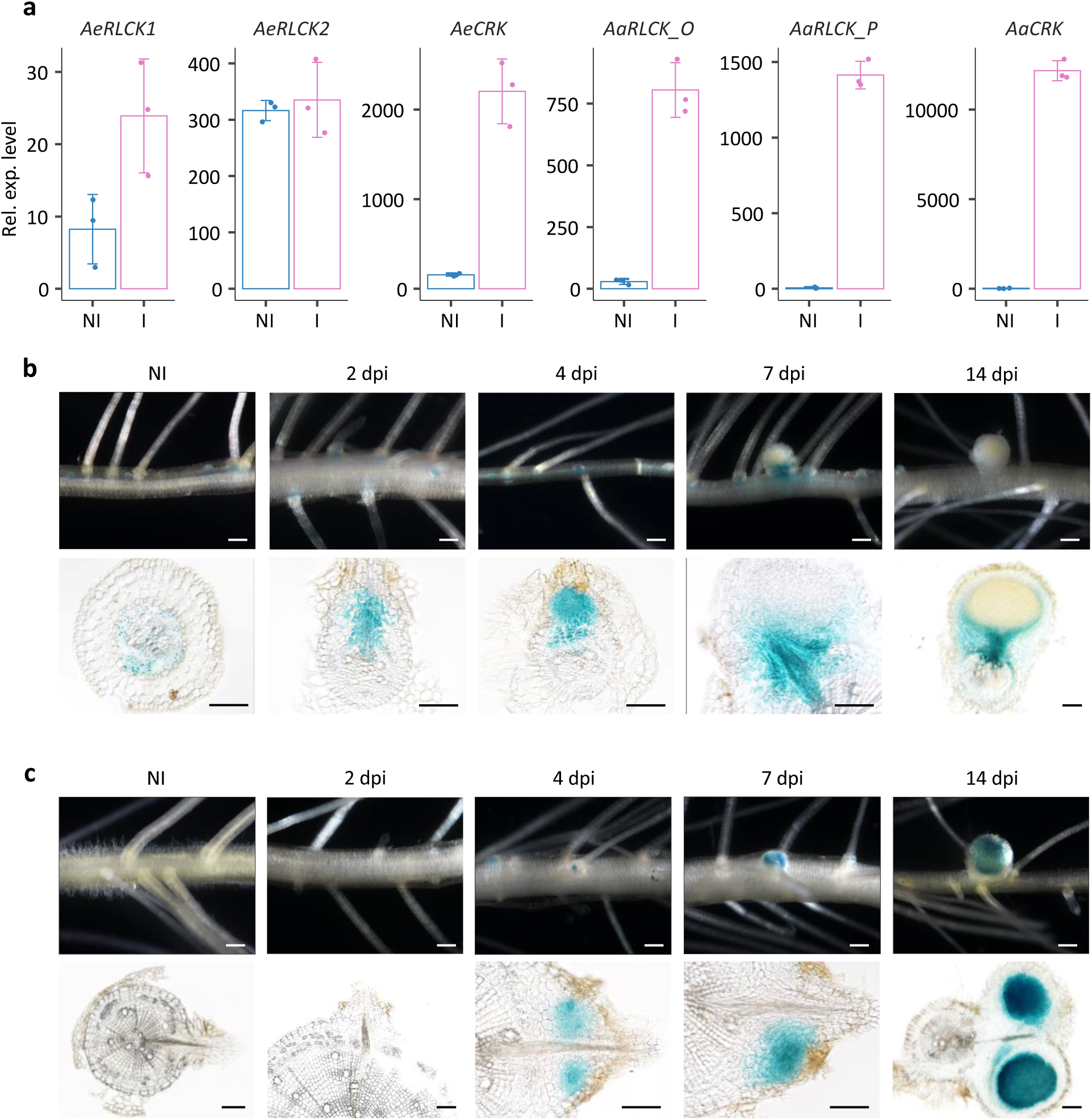
Expression pattern of *AeRLCK2* and comparison with other *Aeschynomene RLCK* and *CRK* genes. **a** RNAseq-based gene expression levels of *AeRLCK1*, *AeRLCK2, AeCRK*, *AaRLCK_O, AaRLCK_P* and *AaCRK* in roots of *A. evenia* and *A. afraspera*, non-inoculated (NI) and inoculated (I) with compatible *Bradyrhizobium* strains. **b** and **c** Histochemical localisation of GUS activity in hairy roots of WT *A. evenia* transformed with *pRLCK2:GUS* (**b**) and *pCRK:GUS* (**c**) during nodulation with *Bradyrhizobium* ORS278. NI: non-inoculated, dpi: days post-inoculation. Top panels: whole roots observed under a light stereomicroscope. Bottom panels: sections of roots and nodules observed by microscope. Scale bars: 1 mm (upper panels), 0.1 mm (bottom panels).

## Discussion

*A. evenia* shares with a few other *Aeschynomene* species a nitrogen-fixing symbiosis with photosynthetic *Bradyrhizobium* strains that is unique among legumes in that its initiation does not depend on the perception of rhizobial Nod factors^(Giraud-2007)^. The molecular processes underlying this Nod-independent symbiosis are still largely unknown. Recently, a forward genetic approach *in A. evenia* identified signalling components that are conserved in other legumes and led to the discovery of AeCRK, a Cysteine-rich RLK^(Quilbé-2021,2022)^. Here, using our mutant-based approach, we have identified a second novel symbiosis actor, AeRLCK2, which corresponds to a Receptor-like Cytoplasmic Kinase. This is another step forward in understanding the Nod-independent symbiosis signalling pathway in *A. evenia*. The 12 *rlck2* mutants show a dual Nod^-^/BN phenotype, indicating a drastic reduction in the ability to initiate nodules. These mutants also lack early responses to rhizobial inoculation such as repression of ARH development, which is the first site of *Bradyrhizobium* colonization, and induction of symbiosis gene expression. In contrast to genes of the conserved symbiosis signalling pathway, for which most mutants are completely Nod*^-^*, all *rlck2* mutants occasionally develop few enlarged nodules. This singular phenotype appears to be inherent to mutations in *AeRLCK2* but it remains to be clarified whether the presence of BN nodules indicates a non-total genetic penetrance for *AeRLCK2* (as for *NOOT* in *M. truncatula*^(Gouzigou-2012)^), the existence of a partial functional redundancy (*e.g*. *RINRK1* in *L. japonicus*^(Li-2019)^) or a function that is distinct from the conserved symbiosis signalling pathway (*e.g*. the infection receptor gene *EPR3* in *L. japonicus*^(Kawaharada-2015)^).

RLCKs lack extracellular ligand-binding domains, but they often functionally and physically associate with plasma membrane localised RLKs to transduce intracellular signals^(Liang-Zhou-2018;Lin-2013)^. The AeRLCK2 protein is unusual among RLCKs in that it has a transmembrane domain. This is essential for its localisation at the plasma membrane. Although its cytoplasmic domain corresponds to a typical Ser/Thr kinase, this activity was weak under *in vitro* conditions and not detected *in planta*. It cannot be ruled out that AeRLCK2 has a kinase activity *in planta* that was not detectable with the antibodies used. But it is also possible that some specific conditions (*e.g*. the RLCK BIK1 is activated by phosphorylation when bacterial flg22 binds to the FLS2-BAK1 complex in *A. thaliana*^(Lee-2017)^) or the presence of interacting partners (*e.g*. several RLCKs have been shown to be strongly and specifically activated by Rop GTPases in *A. thaliana* and *M. truncatula*^(Dorjgotov-2009;Molendijk-2008)^) may be required for AeRLCK2 kinase activity. Protein-protein interaction assays revealed the association of AeRLCK2 and AeCRK *in vitro* and *in vivo*. In contrast to AeRLCK2, AeCRK showed a strong kinase activity and was able to trans-phosphorylate AeRLCK2 both *in vitro* and *in vivo*. This interaction is reminiscent of that between CRK36 and the RLCK BIK1, which is part of the the FLS2-BAK1 receptor complex that perceive bacterial flg22 in *A. thaliana*^(Lee-2017)^. When activated, CRK36 increases BIK1 phosphorylation, leading to increased flg22 signalling and immunity. For AeRLCK2, the residues phosphorylated by AeCRK were identified but it remains to be investigated how these phosphorylation events may regulate AeRLCK2 activity and control nodulation. Additionnally, the biological relevance of the AeCRK-RLCK interaction is supported by the co-expression of *AeCRK* and *AeRLCK2* in nodule primordia infected by *Bradyrhizobium*. Otherwise, their tissular expression patterns are distinct under non-symbiotic conditions and in mature nodules. Furthermore, the nodulation phenotypes of the *crk* and *rlck2* mutants both include early blocks in symbiosis establishment, although these blocks are different^(Quilbé-2021,2022)^. A likely explanation is that AeCRK and AeRLCK2 have overlapping but not identical functions during symbiosis. Therefore, we hypothesize them to have other interacting partners, to form one or more receptor complexes that mediate the Nod-independent symbiosis.

Many RLCKs have been characterized for their involvement in plant development, abiotic stress or immune responses^(Liang-Zhou-2018;Lin-2013)^. However, this analysis is very *A. thaliana*-centered, leaving out RLCKs that have no equivalent in this model plant^(Vij-2008)^. This is the case for the OsRLCK171/LjAMK8/LjAMK28 orthogroup to which AeRLCK1/AeRLCK2 belongs and for which a role in AM has only recently been uncovered^(Leng-2023)^. In *L. japonicus*, LjAMK8 and LjAMK24 interact with the RLK KIN3 and their counterparts in rice, OsRLCK171 and OsARK1, form a similar receptor complex, suggesting that this receptor complex has been evolutionary converved in plants for AM^(Leng-2023)^. Although the expression of *LjAMK8* and *LjAMK24* in nodules suggests that they might also play a role in the rhizobial symbiosis, this remains elusive^(Leng-2023)^. In this respect, AeRLCK2 differs strikingly from its close *L. japonicus* homologs, since in *A. evenia*, *rlck2* mutants are strongly impaired for nodulation but unaltered for mycorrhization. This change in symbiosis involvement correlates perfectly with a tandem gene duplication event that is specific to the the Nod-independent lineage within the genus *Aeschynomene*. In the resulting duplicate *RLCK* genes, *AeRLCK1* is structurally conserved and *AeRLCK2* is more divergent, the latter having a novel promoter sequence. This gene duplication and divergence may have well facilitated the acquisition of the Nod-independent signaling. *AeRLCK1* failed to complement an *rlck2* mutant in *A. evenia*, indicating that it is functionally divergent from *AeRLCK2*. However, *AaRLCK_O*, the *RLCK* homolog in the Nod-dependent *A. afraspera*, was able to rescue the nodulation phenotype of *rlck2* mutant plants. This suggests that AaRLCK_O can still functionally replace AeRLCK2. Based on the available data, the expression pattern of *AeRLCK2* appears to differ from the *RLCK* homologs in *A. afraspera* and *L. japonicus*^(Leng-2023)^. The lack of a genome sequence for *A. afraspera* currently precludes the analysis of tissular gene expression in this species. Despite this, it is likely that the functional specialization of *AeRLCK2* is based on the evolution of promotor specificity and on divergence of protein function with *AeRLCK1*.

From our work and most recent studies^(Leng-2023)^, we propose a model for distinct RLCK involvements in AM through interaction of LjAMK8 and LjAMK24 with LjKIN3 in *L. japonicus* and in the Nod-independent rhizobial symbiosis through interaction of AeRLCK2 with AeCRK in *A. evenia* (Fig. 9). A more comprehensive view of the function of *RLCK* genes in *Aeschynomene* species and the search for other occurrences of Nod-independent specific gene duplications should help us elucidate how the Nod-independent symbiosis evolved. The present advances also pave the way for the identification of additional molecular players that could be involved in the formation of receptor complex(es) with AeCRK and/or AeRLCK2 and mediate the Nod-independent symbiosis pathway in *Aeschynomene* legumes.

**Fig. 9.**
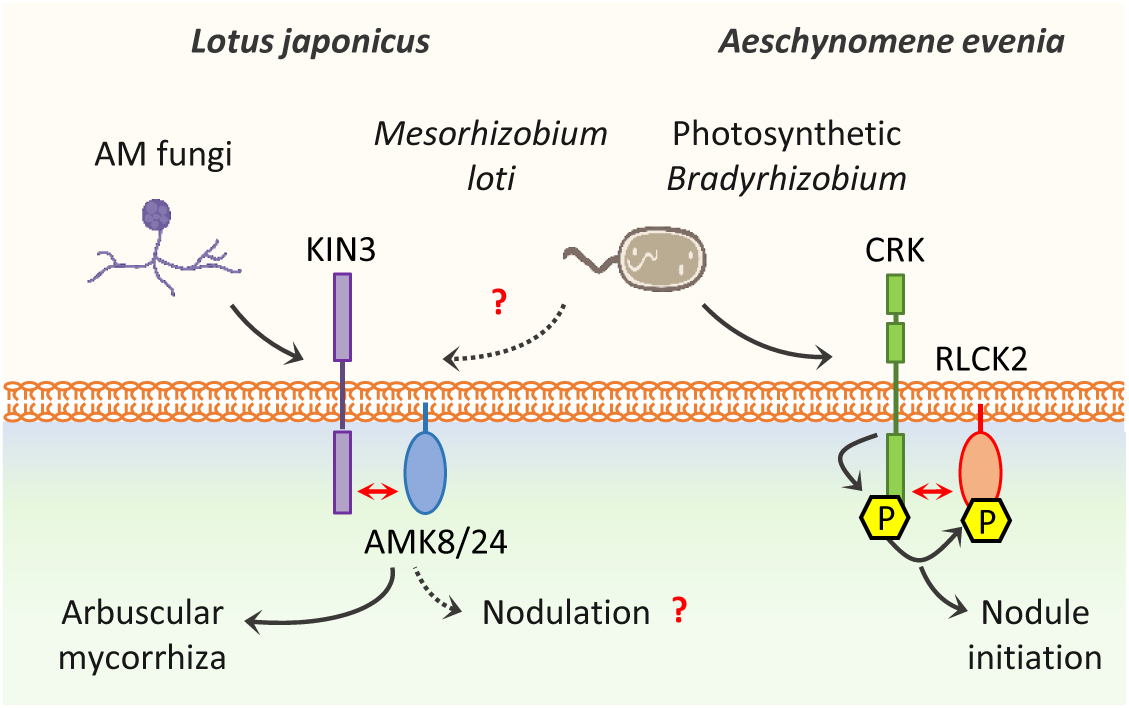
Model of RLCK functions in the AM symbiosis and the Nod-independent symbiosis in legumes. During AM in *L. japonicus*, the paralogs AMK8 and AMK24 interact with KIN3 at the periarbuscular membrane. Autophosphorylation and transphosphorylation events in this RLCK-RLK complex is linked mediate downstream AM responses. In contrast to *LjKIN3*, *LjAMK8* and *LjAMK24* are also expressed during nodulation, but their putative role in the rhizobial symbiosis is not known yet. In *A.evenia*, the LjAMK24 counterpart is absent, while two proteins, AeRLCK1 and AeRLCK2, are closely related to LjAMK8. The symbiotic role of AeRLCK1 is currently unkown whereas AeRLCK2 is central to mediate the Nod-independent symbiosis with photosynthetic bradyrhizobia. One of its fucntions is to interact with and be phosphorylated by AeCRK at the plasma membrane. The upstream signal and downstream signalling components remain to be elucidated.

## Methods

### Plant material and growth conditions

The *A. evenia* lines studied here include the CIAT22838 reference line, mutants derived from this reference line as obtained from a nodulation screen of an EMS-mutagenized population^(Quilbé-2021;Quilbé-2022)^, and the other WT accession PI225551^(Chaintreuil,2018)^ (Supplementary Tables 1 and 2). A selection of *Aeschynomene* species, which use either a Nod-dependent or -independent symbiosis process was also selected^(Brottier-2018)^ (Supplementary Table 1). Seeds were scarified with 96% v/v sulfuric acid for 25-40 min with agitation, and rinsed with distilled water. Scarfied *A.evenia* seeds were incubated overnight with 0,01% (v/v) ethrel (BAYER) to induce germination^(Chaintreuil-2016)^. Plant growth during *in vitro* and in greenhouse conditions according to the protocols established for *Aeschynomene* sp.^(Chaintreuil-2016)^.

### Genetic characterization and sequencing of nodulation mutants

Genetic analyses, consisting of genetic determinism and allelism tests, were performed on *rlck2* mutants following the methodology described previously^(Quilbé-2021)^. Without *a priori* gene identification by mapping-by-sequencing was performed on F_2_ mutant plants obtained from mutant x WT crosses^(Quilbé-2021)^. Illumina sequencing of the F2 mutant DNA pools was performed by the Norwegian Sequencing Center (CEES, Oslo, Norway) and the GeT-PlaGe platorm (INRAE, Toulouse, France). A targeted search for mutations in *AeRLCK2* was performed by PCR-amplification and followed by sequencing for the *de novo* mutation identification in mutant lines or co-segregation analysis in F_2_ mutant plants^(Quilbé-2021)^. The genetic characteristics of the mutants are listed in Supplementary Table 2 and the primers used for *AeRLCK2* sequencing are listed in Supplementary Table 9.

### Plant nodulation

Nodulation assays on *A. evenia* WT CIAT22838 and nodulation mutants were performed using *Bradyrhizobium* ORS278 as inoculum. To analyse the infection process, plants were inoculated with the derivative strains ORS278-GUS and ORS278-GFP^(Giraud-2007;Bonaldi-2011)^. Plant culture, inoculation, determination of the nitrogenase enzyme activity (by measuring the acetylene reduction activity (ARA)), and macoscopic and microscopic observations were performed as described in detail in Supplementary Note 1. Nodulation kinetic experiments with *A. evenia* PI225551 and *A. afraspera* LSTM1 were carried out under standard *in vitro* culture conditions, by inoculating plants with *Bradyrhizobium* ORS285^(Giraud-2007)^ and collecting plant material at 0, 4 and 8 dpi (day-post inoculation).

### Plant mycorrhization

Mycorrhiza studies were performed by inoculating 5 day-old *A. evenia* seedlings with spores of *Rhizophagus irregularis* DAOM197198 (Agronutrition, Carbonne, France) and growing them for 6 weeks^(Nouwen-2024)^. Roots were stained with Sheaffer skrip ink, and fungal colonisation was assessed on 20 root fragments per plant, with 6 plants per line, using the Myco-Calc method as described^(Quilbé-2022)^. Plant mycorrhization was analysed using a Nikon AZ100 stereomicroscope (Champigny-sur-Marne, France) and images were taken using the Nikon Advanced software.

### RNA sequencing analysis and real-time quantitative PCR

RNA was extracted from root material using the RNeasy Plant Mini Kit (Qiagen), treated with DNAse I (RNAse-Free DNAse set, Qiagen) and purified using the RNeasy Min Elute Cleanup Kit (Qiagen), according to the supplier’s protocol. For RNAseq analysis, RNA material was prepared in biological triplicates of 5 plants/replicate for *A. afraspera* at 0, 4 and 8 dpi with *Bradyrhizobium* ORS285, and *A. evenia* CIAT22838 inoculated or not with *R. irregularis* DAOM197198 at 6 wpi (Supplementary Tables 5 and 7). Sequencing libraries were prepared using the TruSeq Stranded mRNA Kit and Illumina sequences generated on the MGX platform (Montpellier Genomix, Institut de Genomique Fonctionnelle, Montpellier France) and the GeT-PlaGe platorm (INRAE, Toulouse, France). *A. afraspera* Illumina RNA-seq datasets were *de novo* assembled using DRAP^(Cabau-2014)^, and those of *A. evenia* were mapped to the *A. evenia* reference genome using nf-core/rnaseq pipeline^(Patel-2024)^. Gene expression levels were normalized using the Diane pipeline^(Cassan-2021)^. For expression analysis, root material was generated in four biological replicates for the *A. evenia* CIAT22838 WT line and nodulation mutants at 0, 2, 4, and 7-dpi with *Bradyrhizobium* ORS278 (3 plants/line/replicate) and at 6 wpi with *R. irregularis* DAOM197198 (5 plants/line/replicate).

RT-qPCR was performed using the Takyon SYBR®Master Mix dTTP Blue kit(Eurogentec) in a 96-well plate format and the Stratagene MX3005P thermocycler (Agilent Technologies). The amplification protocol consisted of the following cycle: 3 minutes at 95°C + 40 cycles of (10 seconds at 95°C + 30 seconds at 60°C + 60 seconds at 95°C + 30 seconds at 60°C) + 30 seconds at 95°C. MXPro software was used to analyze the results based on the cycle threshold (CT) value. The gene expression level was obtained using the formula N=10 ((CT-b)/a) - where a and b vary according to the efficiency of each primer pair. The housekeeping genes *AeEF1α* and *AeUbi* were used for subsequent normalization of expression levels. Primers used for quantification of gene expression are listed in Supplementary Table 10.

### Sequence collection and *in silico* gene analysis

The Ae01g26600 gene is misannotated in the *A. evenia* genome v1. Based on *A. evenia* RNASeq data, it was manually curated to delineate *AeRLCK1* and *AeRLCK2* (Supplementary File 1). Microsynteny analysis was performed using the Legume Information System with the Genome Context Viewer (https://legumeinfo.org/lis_context_viewer) to visualize the gene collinearity in syntenic regions. RLCK protein domains were identified and annotated using InterProscan (http://www.ebi.ac.uk/interpro/) and DeepTMHMM (https://dtu.biolib.com/DeepTMHMM).

AeRLCK2 homologs were identified in legume species by mining the orthogroup database generated with OrthoFinder during the previous *A. evenia* genome project^(Quilbé-2021)^. RLCK sequences were also obtained by BLASTP searches in lupin genomes where *RLCK* genes are present but not annotated and in the *A. afraspera* transcriptome generated in this study. The dataset was completed by searching for additional RLCK proteins in *A. thaliana*, *O. sativa* and *P. persica* in the Arabidopsis Information Resource (https://www.arabidopsis.org), the Rice Genome Annotation Project (http://rice.uga.edu) and the Phytozome (https://phytozome-next.jgi.doe.gov/) databases, respectively. A total of 32 RLCK protein sequences were retrieved from a set of 18 plant species and used for phylogenetic reconstruction (Supplementary Data 1; Supplementary File 1). Identified homologous proteins were aligned using MAFFT v7^(Katoh-2019)^ with the auto strategy, allowing for gapped regions. To optimize the alignments and select the most appropriate approach for Maximum Likelihood analysis, trimAL v1.4.1 was used with the automated-1 option^(Capella-Gutiérrez-2009)^. The resulting alignments were used for phylogenetic analysis using IQ-tree v2.2.0.3^(Minh-2020)^ with the recommended best-fit Model from ModelFinder^(Kalyaanamoorthy-2017)^. Support values were determined with 100,000 iterations of ultrafast bootstrap approximation (UFboot)^(Hoang-2018)^. Tree visualization and annotation was performed using iTOL v6 ^(Letunic-2024)^.

For the analysis of *RLCK* copy number in the genus *Aeschynomene*, additional protein sequences were retrieved in the OrthoFinder-derived RLCK orthogroup for species with available transcriptomes^(Quilbé-2021)^. For an extended set of *Aeschynomene* species (Supplementary Data 2), DNA was extracted using the CTAB method and served as matrix for PCR amplication using different pairs of primers designed as general or copy-specific to amplify an *RLCK* gene fragment in *Aeschynomene* species (Supplementary Table 9). The amplicons were amplified by Sanger technology. Transcriptome and PCR-derived sequences were translated into protein sequences and aligned to AaRLCK_O, AeRLCK1 and AeRLCK2 in Multalin (http://multalin.toulouse.inra.fr/multalin/multalin.html) for comparison (Supplementary Data 2; Supplementary File 2). To resconstruct the phylogeny of *Aeschynomene* species, previously published nuclear *ITS* (Internal Transcribed Spacer) and chloroplast *matK* sequences^(Brottier-2018)^ were concatenated and processed using the same methods described above. The symbiosis type and the presence of the different *RLCK* gene copies were added to the species tree.

### Cloning and plasmid construction

For the initial complementation assay of the *rlck2-11* mutant, the 1212 nucleotide *AeRLCK2* coding sequence (CDS) was PCR-amplified from *A. evenia* cDNAs, and cloned into the CR8/GW/TOPO entry vector, to generate pCR8-*AeRLCK2*. It was then transferred into the pUB-GW-GFP vector via the LR reaction (Invitrogen) to generate the p*Ubi*-*AeRLCK2*-*GFP* construct, in which the *GFP* gene is used as a fluorescent marker for plant transformation.

To analyse *AeRLCK2* and *AeCRK* expression in hairy roots, the *AeRLCK2* promoter (2537 pb upstream of the start codon) and the AeCRK promoter (1381 pb upstream of the start codon) were synthesized and cloned into the Puc57-BSAI-free plasmid by GeneCust (www.genecust.com). The cloned promoters were subsequently fused to the *GUS* gene by GoldenGate cloning, using the pCambia2200-DsRed vector^(Fliegmann-2016)^.

For cross-complementation tests on the *rlck2* mutant, the *AaRLCK_O*, *AeRLCK1* and *AeRLCK2* CDS were PCR-amplified from *A. afraspera* and *A. evenia* cDNAs, respectively, and cloned in the pMiniT 2.0 vector (NEB PCR Cloning Kit, New England Biolabs). Using the GoldenGate cloning method^(Fliegmann-2016)^, they were placed downstream of both the p*LjUbi* and the p*AeRLCK2* promoters in the pCambia2200-DsRed vector, where the *DsRed* gene is used as a fluorescent marker for plant transformation.

For gene expression in *N. benthamiana* leaves and subsequent protein production, the *AaRLCK_O*, *AeRLCK1*, *AeRLCK2* and *AeCRK* CDS without ending stop codon were cloned into pGEMT plasmids. An *AeRLCK2* ^ΔTM^ version, corresponding to the AeRLCK2 protein without the N-ter transmembrane domain (TM) was also produced. Golden gate assembly was performed in the pCambia2200 vector as described^(Fliegmann-2016)^.

For the split-luciferase assay, the N-terminal and C-terminal parts of the luciferase were fused with a triple hemaglutin3HA) or a triple Flag (3Flag) tag, respectively, and flanked by compatible Golden gate extensions. The luciferase modules were synthetized by Azenta (www.azenta.com), and assembled with the CDS to be tested as targets and placed under the control of the p*LjUbi* promoter in a modified pCambia2200 vector by GoldenGate cloning ^(Fliegmann-2016)^.

For *in vitro* assays, the predicted kinase domain of AeRLCK2 (G74 – S403) was amplified using the AeRLCKkinEcoF3 and AeRLCKkinNotR primers and cloned into a modified pCDFDuet-1 vector (Novagen), as described for the cloning of AeCRK kinase^(Quilbé-2022)^. Site-directed mutagenesis was performed to generate the AeCRK^G359E^ (inactive kinase mutation) and AeRLCK2^G110E^ (*rlck2-6* mutant allele mutation) variants using the Q5® Site-Directed Mutagenesis Kit (New England Biolabs). The primers used were AeCRKkin_Mut_G359E_F and AeCRKkin_Mut_G359E_R and the AeRLCK2mutL42F and AeRLCKmutL42R, respectively.

All PCR amplifications were performed using high-fidelity DNA polymerase Taq Phusion (New England Biolabs) or PrimeSTAR Max DNA polymerase (Takara). All constructs were verified by restriction enzyme digestion followed by sequencing with the Sanger technology. Different *E. coli* strains, Dh10b (ThermoFisher Scientific), TOP10 (ThermoFisher Scientific), XL10-Gold (Agilent) and Rosetta/De3 (Novagen Sigma-Aldrich) were used for molecular cloning. Final constructs were electroporated into *Agrobacterium rhizogenes* Arqua1 cells for transformation of *A. evenia* hairy roots^(Quilbé-2021)^ or into *Agrobacterium tumefaciens* LBA4404 VirGN54D for transient expression in *Nicotiana benthamiana* leaves^(Voinnet-2003)^. All primers are listed in Supplemental Table 9.

### Analysis of promoter-GUS and complementation of *A. evenia* transformed hairy-roots

*Agrobacterium rhizogenes* Arqua1 strains carrying the indicated constructs were used to transform roots of WT *A. evenia* CIAT22838 line and *rlck2-11* mutant. Transformation of hairy roots was carried out as previously described^(Quilbé-2021)^. Briefly, two-day-old seedlings with freshly cut radicles were directly inoculated with *A. rhizogenes* Arqua1 carrying the desired plasmid. They were grown on solid MS (Murashige and Skoog basal salt mixture) at 20°C in the dark for five days and then transferred to solid MS medium containing 300µg/mL cefotaxime. Plants bearing transgenic hairy roots were transferred to covered glass tubes containing liquid buffered nodulation media supplemented with 1 mM KNO ^-^. Seven days after transfer, plants were cultivated and inoculated with *Bradyrhizobium* ORS278 according standard procedures. The *A. evenia* WT hairy roots transformed with p*AeRLCK2:GUS* and p*AeCRK:GUS* were stained with X-Gluc to vizualize gene expression at the indicated time-points. For the *A. evenia rlck2-11* complementation assays, nodule formation was monitored 21 days after inoculation.

### Subcellular localisation in *Nicotiana benthamiana* leaves

*N. benthamiana* plants were grown in a controlled environment chamber under the following conditions: 19-21°C with a 16h light/8h dark photoperiod. Four-week-old plants were used for *A. tumefaciens*-mediated transformation to achieve transient protein expression. *A. tumefaciens LBA4404 VirGN54D* strains containing the desired constructs were grown overnight in liquid LB medium, centrifuged at 7 000g for 3 min, and washed twice with agroinfiltration buffer (10 mM MES-KOH pH 5.6, 10 mM MgCl_2_ and 150 µM acetosyringone). The optical density (OD_600_) was measured and adjusted to OD_600_=0.5. *A. tumefaciens* expressing the P19 protein (RNA silencing supressor) was added to the *A. tumefaciens* solutions (OD_600_=0.2) to enhance protein expression^(Voinnet-2003)^. *N. benthamiana* leaves were agroinfiltrated with a needleless syringe and leaves were harvested 3 days later. Subcellular localisation was assessed 72 hours after infiltration with a x25 water immersion objective lens (confocal microscope, SPE8 Leica). MtLYK3-CFP was used as a plasma membrane marker^(Klaus-Heisen-2011)^ and co-expressed with YFP fusion proteins. The excitation/emission filter sets for CFP and YFP were 458 nm/463-512 nm and 514 nm/525-580 nm, respectively.

### Protein-protein interaction assays

Different construct combinations (MtLYK3-CFP, AeCRK^G359E^-YFP, AeRLCK2-YFP, AeRLCK2^ΔTM^-YFP and AeCRK^G359E^-mCherry) were agroinfiltrated into *N. benthamiana* leaves together with p19. Leaf material was collected 3 days after infiltration, and proteins were extracted for co-immunoprecipitation assays as previously published^(Ding-2024)^. YFP fusion bands were quantified using Image Lab 6.0 (volume function). Enrichment ratios (signal IP αGFP / signal input αGFP) were normalized using the negative control MtLYK3-CFP (signal αRFP).

The split-luciferase assay was performed as previously described^(Landry-2023)^. Briefly, *A. tumefaciens LBA4404 VirGN54D* strains containing the indicated NLuc and CLuc plasmids were co-infiltrated into 4- week-old *N. benthamiana* leaves. After 72 h, 4 mm leaf discs were placed in a 96-well plate, washed twice with water, and incubated with 1 mM luciferin substrate (Xenolight, PerkinElmer). Light emission was then quantified using a luminometer (VICTOR Nivo, PerkinElmer). Protein expression levels were assessed from 7 mm leaf discs, which were previously used to quantify luciferase activity. After protein extraction and Western blotting, 3HA-NLuc fusion bands were quantified using the volume function in Image Lab 6.0. The corresponding data were used to calculate a ratio normalized to the negative control AeCRK^G359E^-Cluc/MtLYK3-Nluc. Raw data were normalized using the previously calculated ratio to account for variation in protein expression.

For Western blotting, leaf samples were ground in liquid nitrogen using a Retsch mixer mill (MM400). Proteins were solubilized in Laemmli buffer, heat-denatured at 95°C for 5 min and separated by SDS-PAGE on home-made gels or 4-15% precast polyacrylamide gel (Bio-rad).

Proteins were transferred to a nitrocellulose membrane using the Transblot-Turbo system (Bio-rad) according to the manufacturer’s instructions. The nitrocellulose membrane was blocked for one hour at RT (or overnight at 4°C) with 5% non-fat milk or 3% BSA solution in Tris-saline buffer (TBS) supplemented with 0.1% Tween-20 (TBS-T). As a loading control, Ponceau S staining solution was used to visualize Rubisco protein. The membrane was then incubated with the appropriate antibodies for one hour at RT (or overnight at 4°C). The following antibodies were used for protein biochemistry experiments: αHA-Hrp (12013819001, Roche, 1/5000), αFlag-Hrp (A8592, Sigma-Aldrich, 1/5000), αPhospho-Ser-thr-tyr (61-8300, Invitrogen, 1/333), αPhospho-tyr (1/2000, Zymed), αRabbit-Hrp (12-348, Millipore, 1/20000), αGFP (11814460001, Roche, 1/3000), goat αMouse-Hrp (1706516, Bio-Rad, 1/10000), rabbit αRFP (1/5000)^(Lefebvre-2012)^. Hrp bioluminescence was detected using Clarity Western ECL substrate (Biorad) and observed using the ChemiDoc imager (Biorad).

### *In vitro* and *in planta* phosphorylation assays

For *in vitro* assays, *AeRLCK2-kin* and its mutant form *AeRLCK2-kin^G110E^*, and *AeCRK-kin* and its mutated form *AeCRK-kin*^G359E^, were cloned into a modified pCDFDuet-1 vector (Novagen) and expressed as fusion proteins at 16°C. Proteins were purified using Glutathione-Sepharose4B (Amersham Biosciences) as described^(Klaus-Heisen-2011)^. AeCRKkin was released from the resin using PreScission Protease (GE27-0843-01, Sigma Aldrich, Germany). For kinase assays, proteins were incubated for 50 min at 30°C in 10 mM HEPES-HCl pH 7.4 containing 5 mM MgCl2, 5 mM MnCl2, 20 mM ATP and 5 mCi 32P-ATP. Reactions were analysed by SDS-PAGE, followed by Coomassie staining and Phosphor imaging. Bands were quantified using the volume function in Image Lab 6.0 (Bio-Rad). Protein purifications and kinase assays were repeated at least twice.

For *in planta* phosphorylation assays, proteins were extracted (w/v, 0.2/1) with protein extraction buffer (50 mM Tris-HCl 7.5, 150 mM NaCl, 10 mM EDTA, Triton X-100 1%, DTT 2 mM, supplemented with protease inhibitor cocktail (Sigma) and phosphatase inhibitor cocktail 3 (Sigma)). Proteins were solubilized for 30 min at 4°C and then centrifuged at 20 000g for 5 min at 4°C. The supernatant was filtered through Miracloth and incubated for two hours at 4°C with αGFP magnetic agarose beads (Chromotek). The beads were washed three times with protein extraction buffer. Proteins were solubilized in Laemmli 2X buffer, heat denaturated at 95°C and subsequently separated by SDS-PAGE. The phosphorylation status was assessed using α-Phospho-serine-threonine-tyrosine and αphosphorylated threonine. Identification of phosphorylated sites by LC-MS/MS analysis was performed as described in detail in Supplementary Note 2.

### Statistical analysis and graphs

Data analysis and visualization were performed using R with theggplot2 package^(RCoreteam-2020; Whickham-2016)^. The Kruskal-Wallis test, followed by Dunn’s post hoc test for multiple comparisons, was used for all statistical analysis^(Kruskal-1952; Dunn-1964)^.

### Data availability

All data are available in the main text or the supplementary material. Source Data are provided with this paper. The datasets, plasmid constructs and plant materials generated and analyzed in this study are available on request from the corresponding authors.

## Acknowledgements

We thank Robin Duponnois (LSTM Laboratory, IRD) for assistance with the characterization of the *A. evenia* nodulation mutants and for kindly providing fungi spores for mycorrhization experiments. We would also like to thank Virginie Gasciolli, Céline Vicedo and Léandre Bouat (LIPME Laboratory, INRAE) for their technical assistance. Illumina sequence data were produced by the MGX platform (https://www.mgx.cnrs.fr/), the Norwegian Sequncing Centre (http://www.sequencing.uio.no) and the GeT-PlaGe platform (https://get.genotoul.fr/la-plateforme/get-plage/). Computing was performed thanks to the GenoToul bioinformatics facility (http://bioinfo.genotoul.fr/). The project also benefited from the expertise of the Proteomics French Infrastructure (https://www.profiproteomics.fr/) and France-BioImaging Infrastructure (https://france-bioimaging.org/) located at the Agrobiosciences, Interactions and Biodiversity Research Federation (https://www.fraib.fr/). This study was supported by four grants from the French National Research Agency (ANR-SymWay-21-CE20-0011-01, ANR-DUALITY-20-CE20-0017, ANR-AeschyNod-14-CE19-0005-01 and ANR-BugsInaCell-13-BSV7-0013-02).

## Author contributions

J.F.A. and B.L. conceived the whole project and supervised data analyses. J.Q. performed the genetic and molecular analysis of the *rlck2* mutants, to which J.F.A. contributed. N.H.A conducted the phenotypic characterization of the *rlck2* mutants and RT-qPCR analyses relative to the nodulation and AM tests, to which J.Q. and M.P. contributed. N.H.A., D.L., J.Q. and C.G. conducted phylogenetic and evolutionary analyses. N.H.A., J.Q. and M.R. generated molecular constructs and conducted functional experiments on *AeRLCK2*. D.L. performed the biochemical characterization of AeRLCK2 and AeCRK, to which J.C. and C.P. contributed. M.P., F.G., and D.G. produced plant material and RNA material. LB screened the *A. evenia* mutagenized population to isolate *rlck2* mutants. C.K. analyzed the sequence data for the Mapping-by-Sequencing analysis of the *rlck2* mutants as well as for the RNAseq data. M.P., F.G., C.C., N.N. and E.G. contributed to different experiments and provided their assistance for the achievement of the project. J.F.A., N.H.A. and D.L. wrote the manuscript. N.H.A produced the figures, to which D.L. contributed. All authors critically commented on and approved the manuscript.

## Additional information

Correspondence and requests for materials should be addressed to J.-F.A. and B.L.

