## Supplemental Notes, Figures, Tables for "A Receptor Like Cytoplasmic Kinase evolved in *Aeschynomene* legumes to mediate Nod-independent rhizobial symbiosis"

### Supplementary Notes

#### Supplementary Note 1. Plant nodulation with *Bradyrhizobium* ORS278

##### *In vitro* nodulation tests

For the *in vitro* nodulation test, one-day-old seedlings were transferred to 0.8% agar-water plates at 37°C for 24 hours to achieve at least 1 cm of radicle growth. They were then transferred to covered glass tubes containing liquid buffered nodulation medium (BNM) supplemented with 0,5 mM $KNO_3^-$, as described in detail[(Arrighi-2012)]. Seven days after transfer in tubes, plants were inoculated with 1 mL of bacterial culture per plant, adjusted to an OD600=1 using a spectrophotometer (Varian, UV-visible spectrophotometer Cary 50 scan). At 7, 10, 14, 17 and 21 dpi, the number of red and white nodules was assessed and counted using a binocular loupe. Other plant analyses were carried out on samples collected at 21 days post-inoculation (dpi). Nodule and axillary root hair diameters were measured using ImageJ (version 2.14.0/1.54f, http://imageJ.nih.gov/ij), while the number of lateral roots per plant was evaluated using the Optimas software (version 6.1, Media Cybernitics, Silverspring, MD, USA). Nitrogenase enzyme activity is assessed by analysizing the reduction acetylene to ethylene (ARA - Acetylene Reduction Assay) on plants with nodules, as described[(Arrighi-2012)].

##### Greenhouse nodulation studies

For greenhouse experiments, scarified seeds were left overnight under gentle agitation (80 rpm) to induce radicle emergence. The next day they were transplanted into plastic trays containing attapulgite (Dry Oil, US Sorbix Special). Plants were inoculated at transplantating with 150 mL of ORS278 culture (OD600=1) per plastic tray (50x40 cm, 70 to 150 plants per tray) and grown for 4 weeks before root observation.

##### Macroscopic and microscopic observations

All root samples from non-inoculated and inoculated roots were visually inspected using a Nikon AZ100 stereomicroscope (Champigny-sur-Marne, France) and imaged using the Nikon Advanced software. Where needed, fresh 42 micron thick section of roots and nodules were made using a Leica VT1000s vibratome. For bacterial infection analysis (ORS278-GUS) and promoter activity studies (Prom-GUS), Whole plant roots or sectioned samples were stained with X-Gluc[(Fabre-2015)]and then analysed using a Nikon macroscope or microscope. Gene expression was observed at different time points in both young and older non-inoculated plants, and at 2, 4, 7, 10, 14 and 21 dpi for plants inoculated with ORS278. To investigate bacterial infection with ORS278, freshly sectioned nodules were incubated with the live/dead reagent (Syto9 /propidium iodide) and then stained with calcofluor white, as previously described[(Nouwen-2024)]. Samples were analysed using a confocal laser-scanning microscope (Carl Zeiss LSM 700, Jena, Germany). Calcofluor was excited at 405 nm and emitted light collected between within 405–470 nm, while SYTO 9 and propidium iodide were excited at 488 nm and 555 nm, respectively, with emissions collected between 490–522 nm and 555–700 nm. Images were acquired using the ZEN 2008 software (Zeiss, Oberkochen, Germany).

#### Supplementary Note 2. Identification of AeRLCK2 phosphorylation sites

##### Sample preparation and LC-MS/MS spectrometric analysis

To identify the phosphosites of AeCRK and AeRLCK2, the YFP-tagged constructs were transiently expressed in *N. benthamiana* leaves*.* Samples were harvested 3 days after agro-infiltration. Leaves were crushed in a mortar with liquid nitrogen. Proteins were solubilized with protein extraction buffer (1:5 (w/v), 150 mM NaCl, 50 mM Tris-HCl, 10 mM EDTA, 1% Triton X-100, 2 mM DTT, 5 µM MG132, supplemented with commercial phosphatase and protease inhibitor cocktail (Roche)). Samples were centrifuged at 16 000g for 20 minutes at 4°C. The supernatant was filtered through miracloth and proteins were immunoprecipitated using magnetic agarose GFP-beads (Chromotek). Proteins were solubilized in L2X and boiled at 95°C. They were separated by SDS-PAGE and stained with Instant Blue. The bands were then cut and prepared for mass spectrometry analysis. To remove staining, the bands were washed 5 times alternately with 100 mM $NH_4HCO_3$ and 50 mM $NH_4HCO_3$/Acetonitrile (CAN) (1:1). The bands were incubated with 100% ACN for 10 min. Proteins were reduced for 45 min at 56°C by adding 10 mM DTT prepared in 100 mM $NH_4HCO_3$. The alkylation step was conducted by adding 55 mM iodoacetamine in 100 mM $NH_4HCO_3$. Bands were washed twice with 50 mM $NH_4HCO_3$ and peptides were digested overnight with trypsin enzyme (10 ng/uL, sequence grade Promega). Peptides were extracted with 1% ACN/ 1% formic acid and dried using a speed vac. Peptides were resuspended with 15 µL of 2% Acetonitrile, 0.05% TFA, vortexed and sonicated for 10 min before injection. Peptide mixtures were analysed by nano-LC-MS/MS using nanoRS UHPLC system (Dionex, Amsterdam, The Netherlands) coupled to a Q-Exactive Plus mass spectrometer (Thermo Fisher Scientific, Bremen, Germany). Five microlitres of each sample were loaded on a C18 pre-column (5 mm × 300 µm; Thermo Fisher) at 20 µL/min in 5% acetonitrile, 0.05% trifluoroacetic acid. After 5 min of desalting, the pre-column was switched on line with the analytical C18 column (15 cm × 75 µm; Reprosil C18 in-house packed) equilibrated in 95% of solvant A (5% acetonitrile + 0.2% formic acid in water) and 5% of solvant B (80% acetonitrile + 0.2% formic acid in water). Peptides were eluted using a 5-50% gradient of B for 105 min at a flow rate of 300 nL/min. The Q-Exactive Plus was operated in data-dependent acquisition mode using the Xcalibur software. Survey scan MS spectra were acquired in the Orbitrap in the 350-1500 m/z range with a resolution of 70 000. The ten most intense ions per survey scan were selected for HCD fragmentation, and the resulting fragments were analysed in the Orbitrap at a resolution of 17 500. Dynamic exclusion was used within 30s to avoid repetitive selection of the same peptide. Acquired MS and MSMS data were searched using Mascot (version 2.8.0.1, http://matrixscience.com) against a custom database of all protein sequences of interest and contaminant protein sequences. The search included methionine oxidation, N-ter acetylation, S, T and Y phosphorylation as variable modifications and carbamidomethylation of cysteine as a fixed modification. Trypsin was chosen as the enzyme and 2 missed cleavages were allowed. The mass tolerance was set to 10 ppm for the precursor ions and to 20 mmu for fragment ions. Raw MS signal extraction of identified peptides was performed across different samples. Validation of identifications was performed by a false-discovery rate at 1% at the protein and peptide-sequence match level, determined by target-decoy search using the in-house-developed software Proline[(Bouyssié-2020)] software version 2.1 (http://proline.profiproteomics.fr/). The mass spectrometry proteomics data have been deposited to the ProteomeXchange Consortium with the PRIDE partner repository under the dataset identifier PXD053561.

**Alphafold prediction**

The AlphaFold server was used for the AlphaFold prediction, [(Abramson-2024)]. The highest-ranked model was selected. For AeRLCK2, the prediction metrics indicating the accuracy of the prediction showed a pTM score of 0.74. The protein structure was analyzed and annotated using ChimeraX software version 1.7.1.

### Supplementary Figures

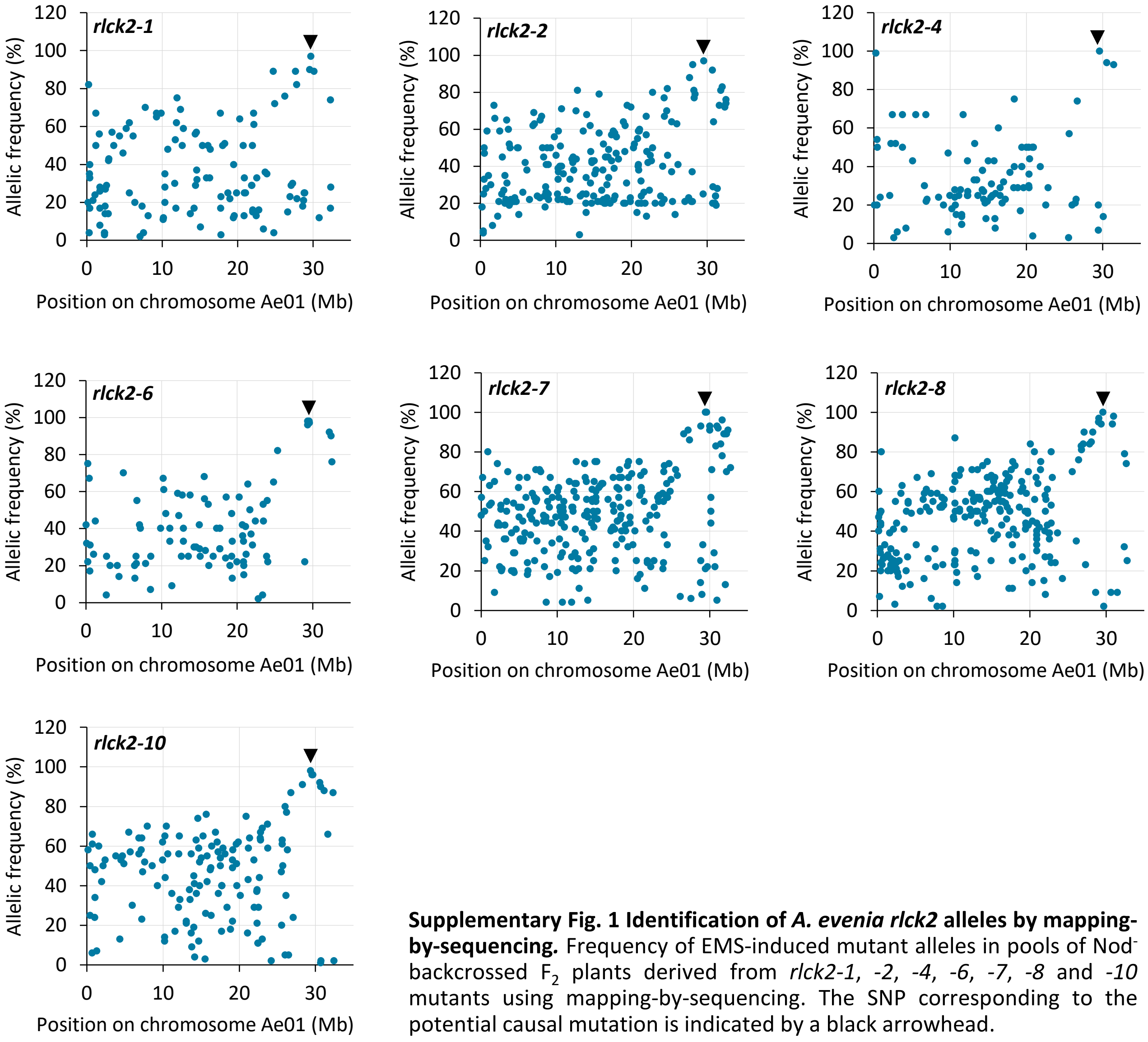


**Supplementary Fig. 1 Identification of *A. evenia rlck2* alleles by mapping-by-sequencing.** Frequency of EMS-induced mutant alleles in pools of Nod$^-$ backcrossed $F_2$ plants derived from *rlck2-1*, *-2*, *-4*, *-6*, *-7*, *-8* and *-10* mutants using mapping-by-sequencing. The SNP corresponding to the potential causal mutation is indicated by a black arrowhead.

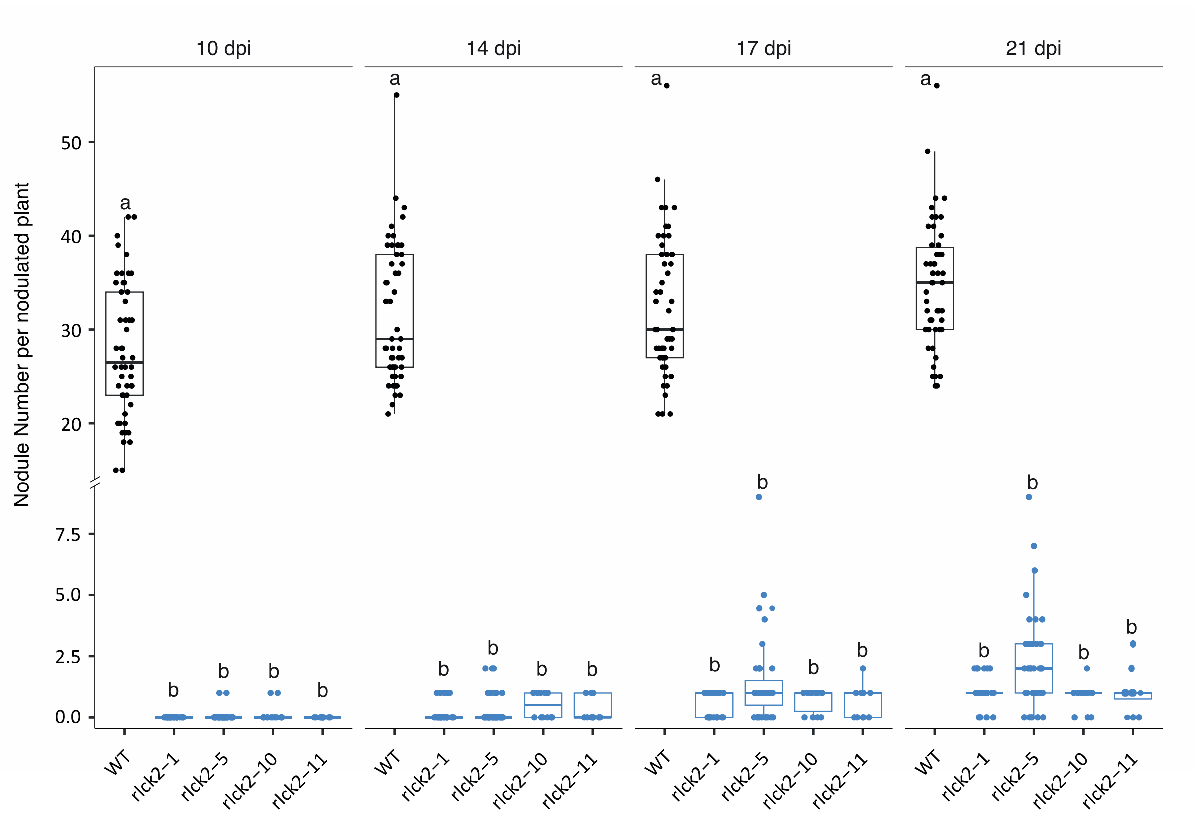


**Supplementary Fig. 2 Nodulation kinetics of WT and *rlck2* mutants.** Pink nodule numbers at 10 to 21 days post-inoculation with *Bradyrhizobium* ORS278. Analysis of variance (Kruskal-Wallis) and post-hoc analysis (Dunn's test), different letters indicate significant difference: p<0,05, with 3 biological replicates. Nodulated plants from 3 independent experiments.

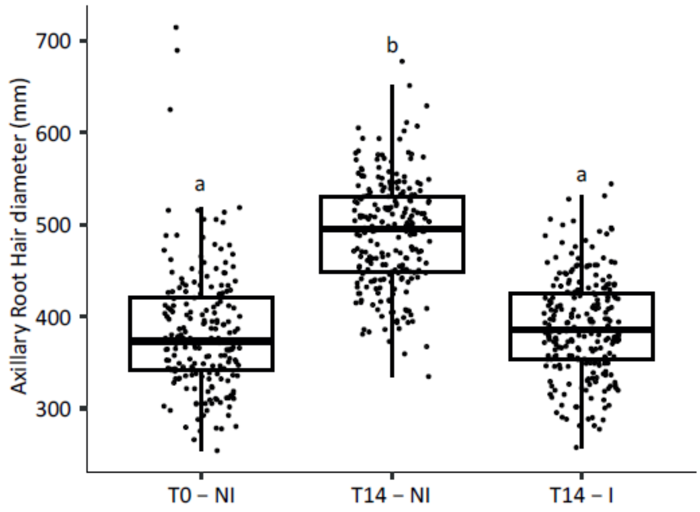


**Supplementary Fig. 3 Development of axillary root hairs in the *crk-1* mutant.** Diameter of axillary root hair crowns at lateral root bases at different time points in non-inoculated (NI) and inoculated (I) plants with *Bradyrhizobium* ORS278. T0: time of inoculation. T14: 14 days after inoculation. Different letters above the box borders indicate significant differences between samples. Analysis of variance (Kruskal-Wallis) and post-hoc analysis (Dunn's test), p<0,05.

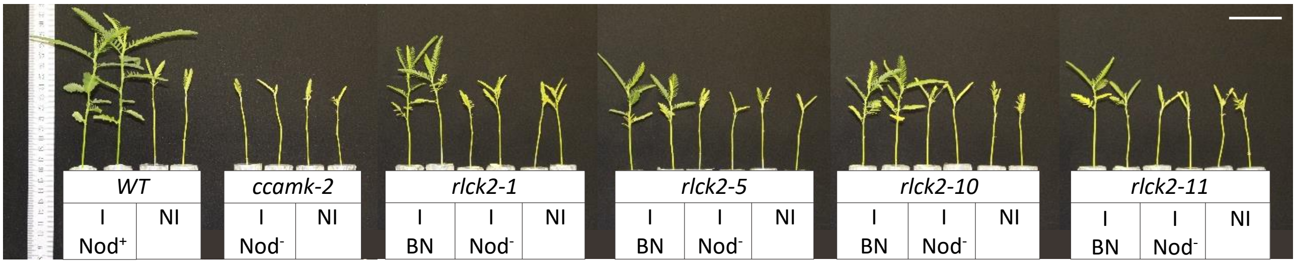


**Supplementary Fig. 4 Aerial phenotype of WT and *rlck2* plants grown in growth chamber.** Images showing the aerial phenotype of non-inoculated (NI) and inoculated (I) plants with *Bradyrhizobium* ORS278 at 21 dpi. $Nod^{+}$: nodules, $Nod^{-}$: no nodules, BN: big nodules. Scale bar: 5 cm.

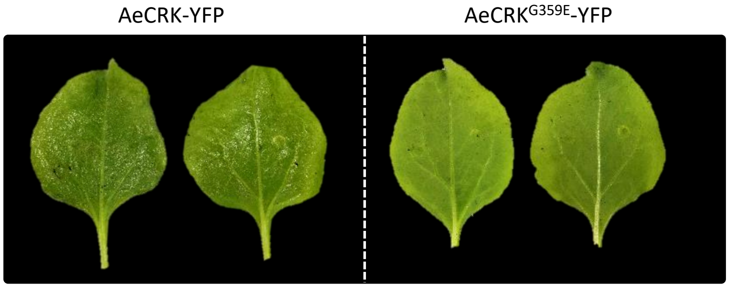


**Supplementary Fig. 5 AeCRK induces a cell death in *Nicotiana benthamiana* leaves.** Images showing cell death induced by the transient expression of *AeCRK-YFP* in *N. benthamiana* leaves. In contrast, the kinase version mutated in the glycine-rich loop motif (G359E) did not induce cell death. Leaved were imaged five days post-infiltration.

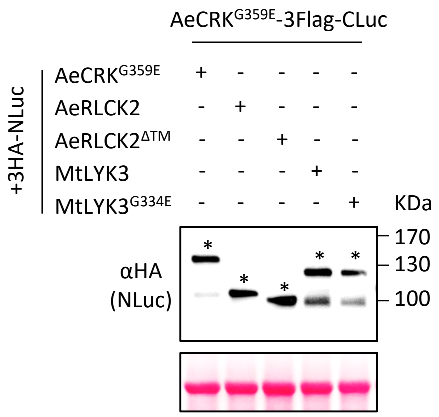


**Supplementary Fig. 6 Expression of 3HA-NLuc fusion proteins.** For each split-luciferase replicate, the expression levels of the 3HA-Nluc fusion proteins were assessed by Western blotting using αHA antibodies. Asterisks indicate the corresponding NLuc fusion proteins. These Western blot analyses were used to quantify the expression levels of NLuc constructs and to normalize luminescence levels. Ponceau staining was used as a loading control.

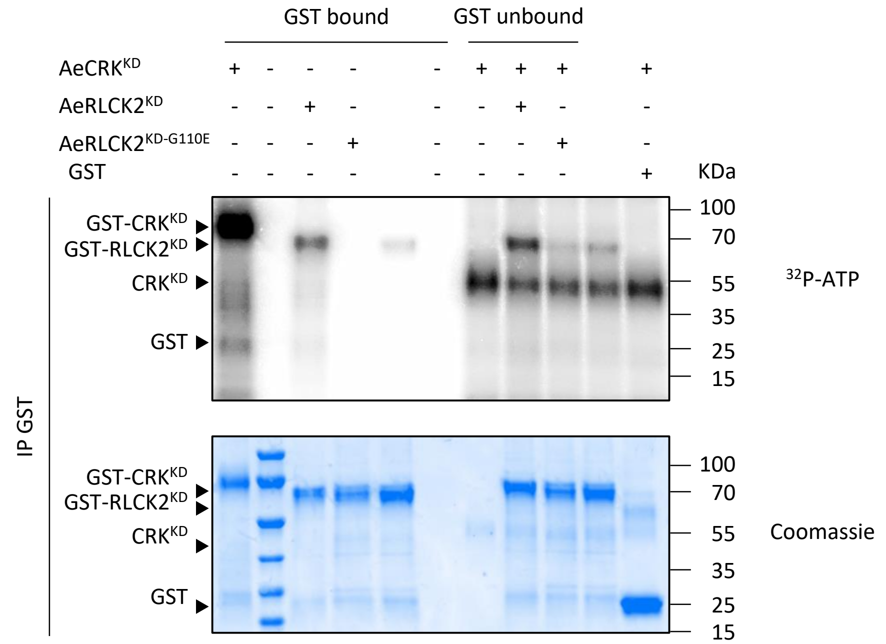


**Supplementary Fig. 7 AeCRK kinase activity and AeRLCK2 transphosphorylation *in vitro.*** Autoradiography showing the potent kinase activity of AeCRK and its ability to transphosphorylate AeRLCK2. The kinase domains (KD) were produced in *Escherichia coli* and immunopurified using glutathione agarose beads. *In vitro* kinase activity was assessed using radiolabeled ATP ($^{32}$P-ATP). For the transphosphorylation assay, the GST tag of AeCRK$^{KD}$ was removed using PreScission protease. Coomassie staining served as the loading control.

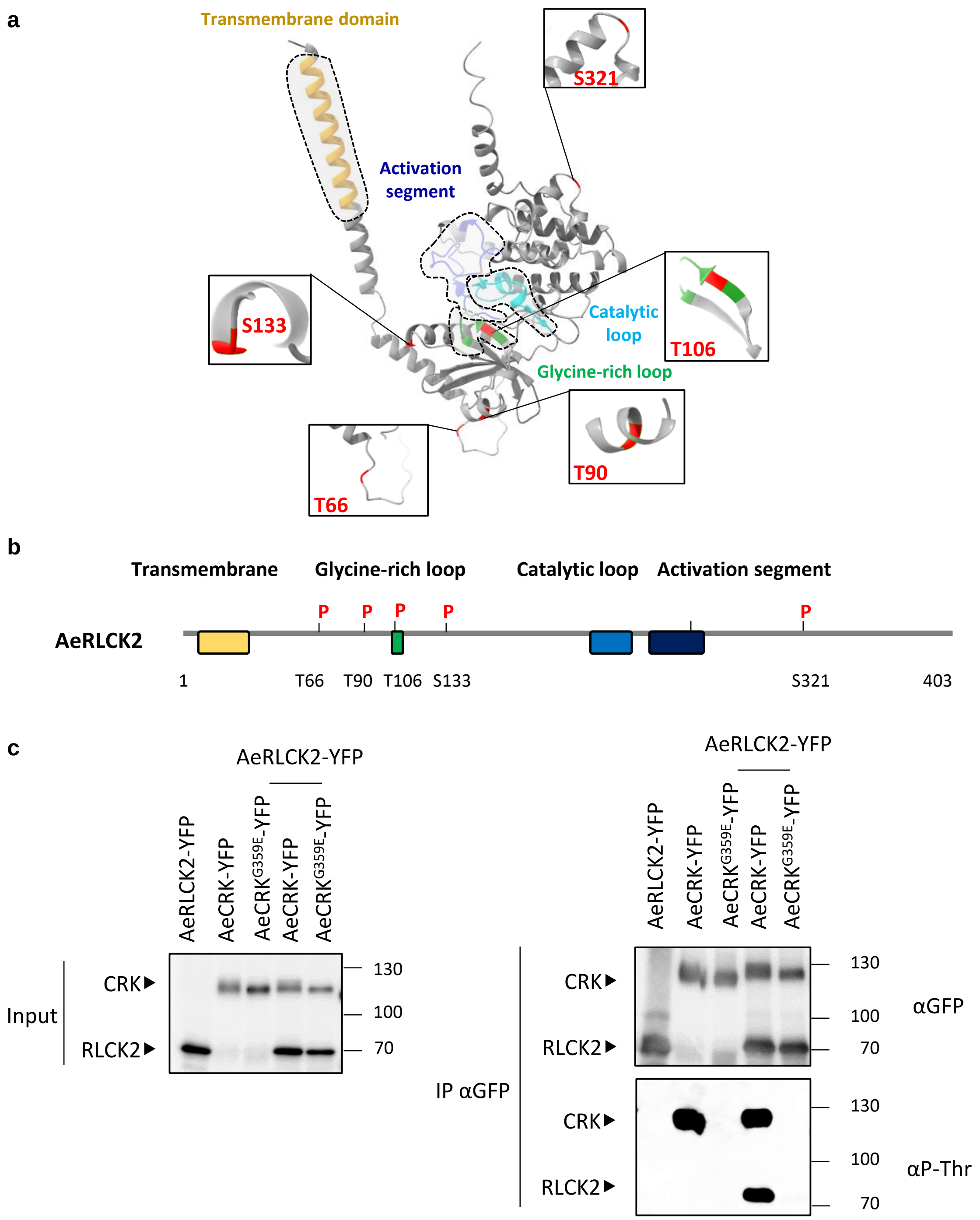


**Supplementary Fig. 8 The phopshorylation sites of AeRLCK2 specifically targeted by AeCRK. a** The 3D structure of AeRLCK2 was obtained using Alphafold server prediction. The top-ranked prediction is displayed, with kinase subdomains and the transmembrane domain highlighted by a grey circle and distinct colours. Phosphosites of AeRLCK targeted by the kinase activity of AeCRK are marked in red. A total of five residues were identified accross different kinase subdomains. Residue T106 is located in the glycine-rich loop. **b** The protein schematic structure of AeRLCK2 with phosphorylation sites targeted by AeCRK and kinase domains are shown. **c** AeRLCK2 is specifically phosphorylated by AeCRK mostly at threonine residues. The YFP-tagged AeRLCK2, AeCRK and AeCRK$^{G359E}$ were transiently expressed in *N. benthamiana* leaves. Total proteins were extracted, and YFP-tagged proteins were immunopurified. The phosphorylation status of immunopurified AeRLCK2, AeCRK and AeCRK$^{G359E}$ was assessed using an antibody recognizing phosphorylated threonine (αP-Thr). The right panel shows the input, while the left panel corresponds to immunopurified proteins.

**Residue T66**

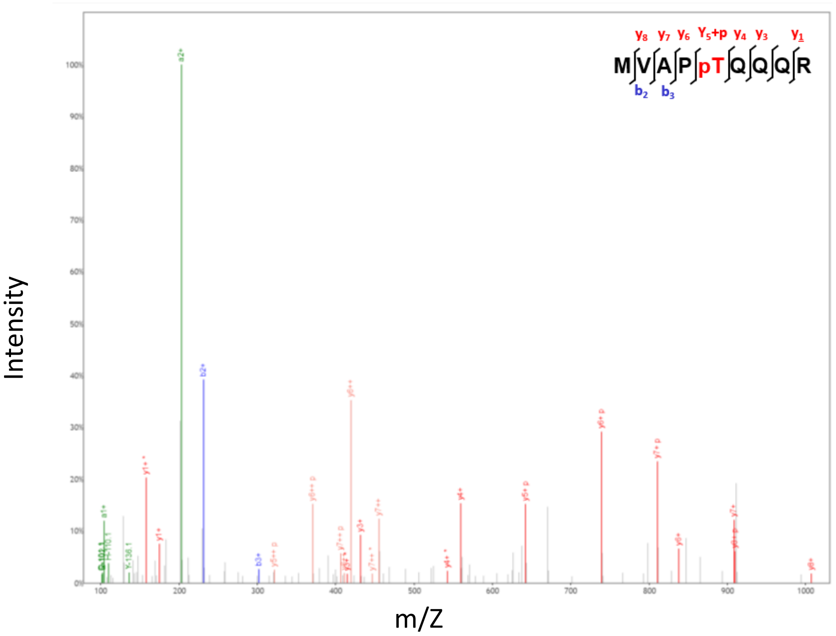


**Residue T90**

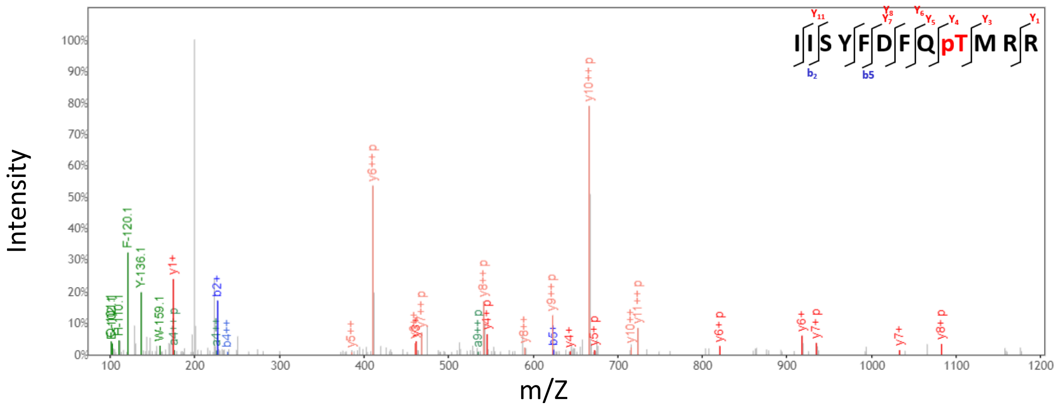


**Residue T106**

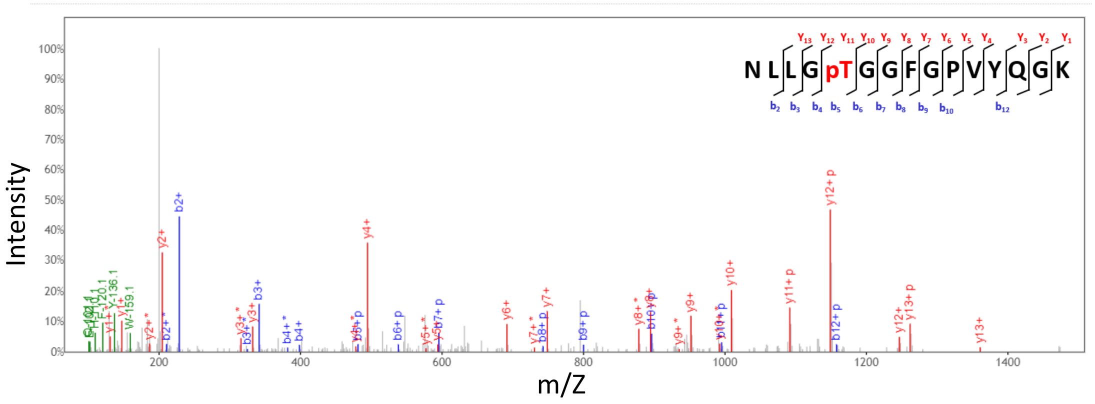


**Supplementary Fig. 9 MS/MS fragmentation spectra supporting the Supplementary Fig. 8.** The original raw fragmentation spectrum for each peptide targeted by the kinase activity of AeCRK is shown.

**Residue S133**

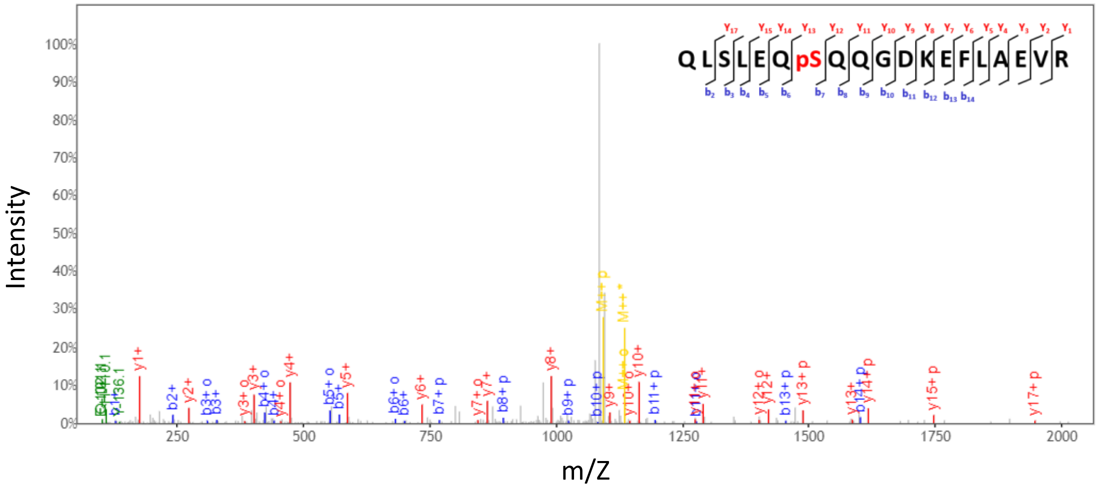


**Residue S321**

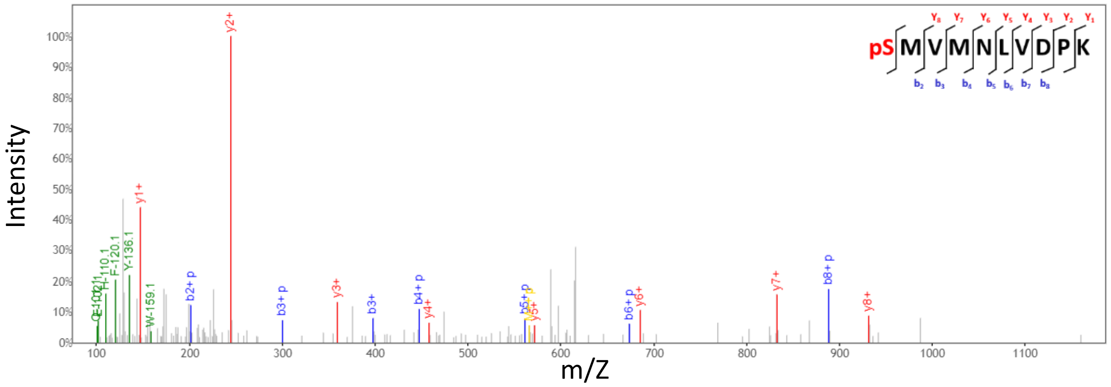


**Supplementary Fig. 9 following MS/MS fragmentation spectra supporting the Supplementary Fig. 8.** The original raw fragmentation spectrum for each peptide targeted by the kinase activity of AeCRK is shown.

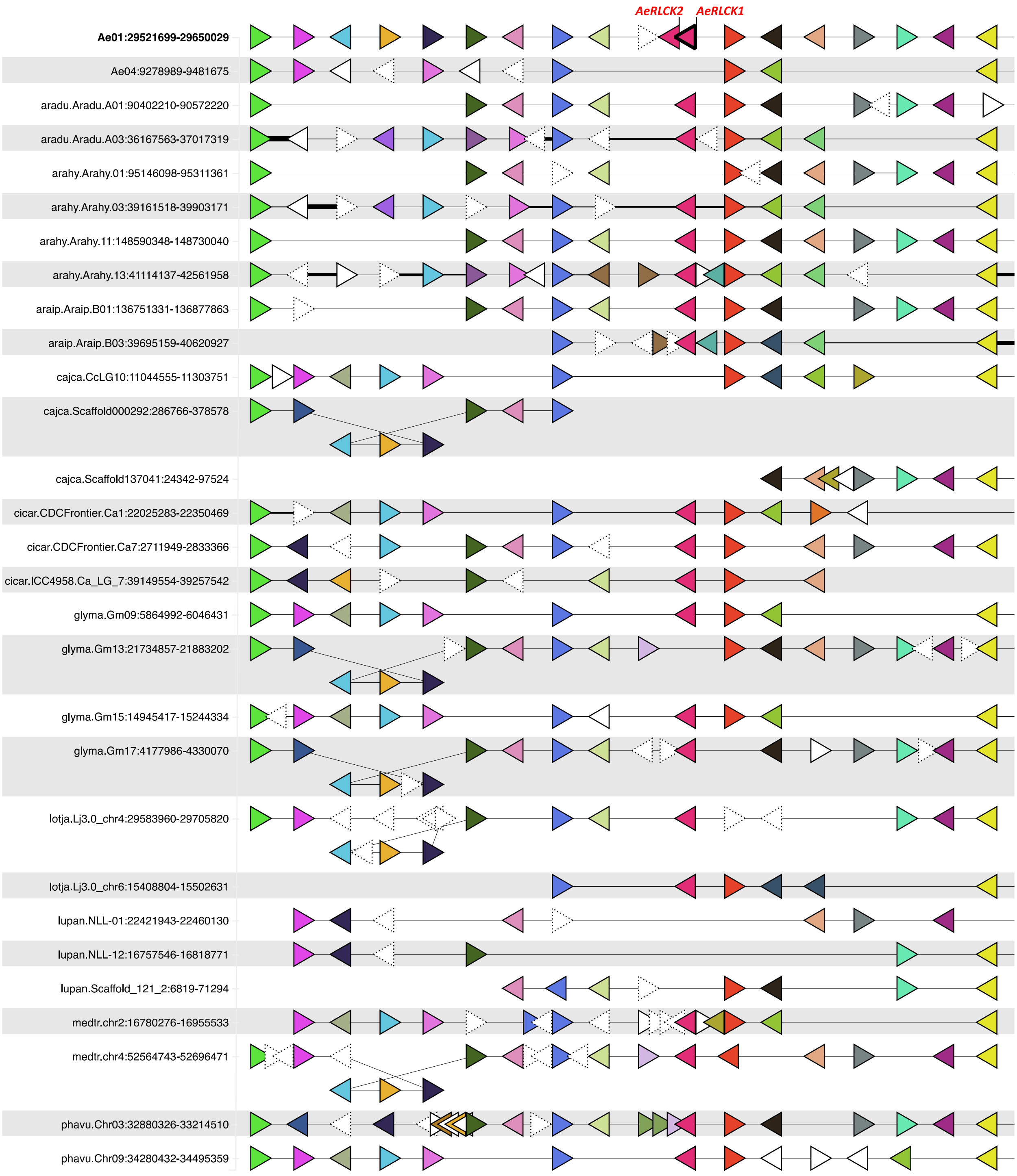


**Supplementary Fig. 10 Microsynteny analysis of the *AeRLCK2* locus.** Schematic representation of the conservation of gene colinearity at the *RLCK* locus. The same color code indicates homologous gene pairs. Note in *A. evenia*, the presence of the *RLCK1-RLCK2* gene tandem on chromosome Ae01 and the absence of a *RLCK* paralog on chromosome Ae04. These two features are not found in other legume species. Data were obtained from the Legume Information System (https://legumeinfo.org/) using the genome context viewer.

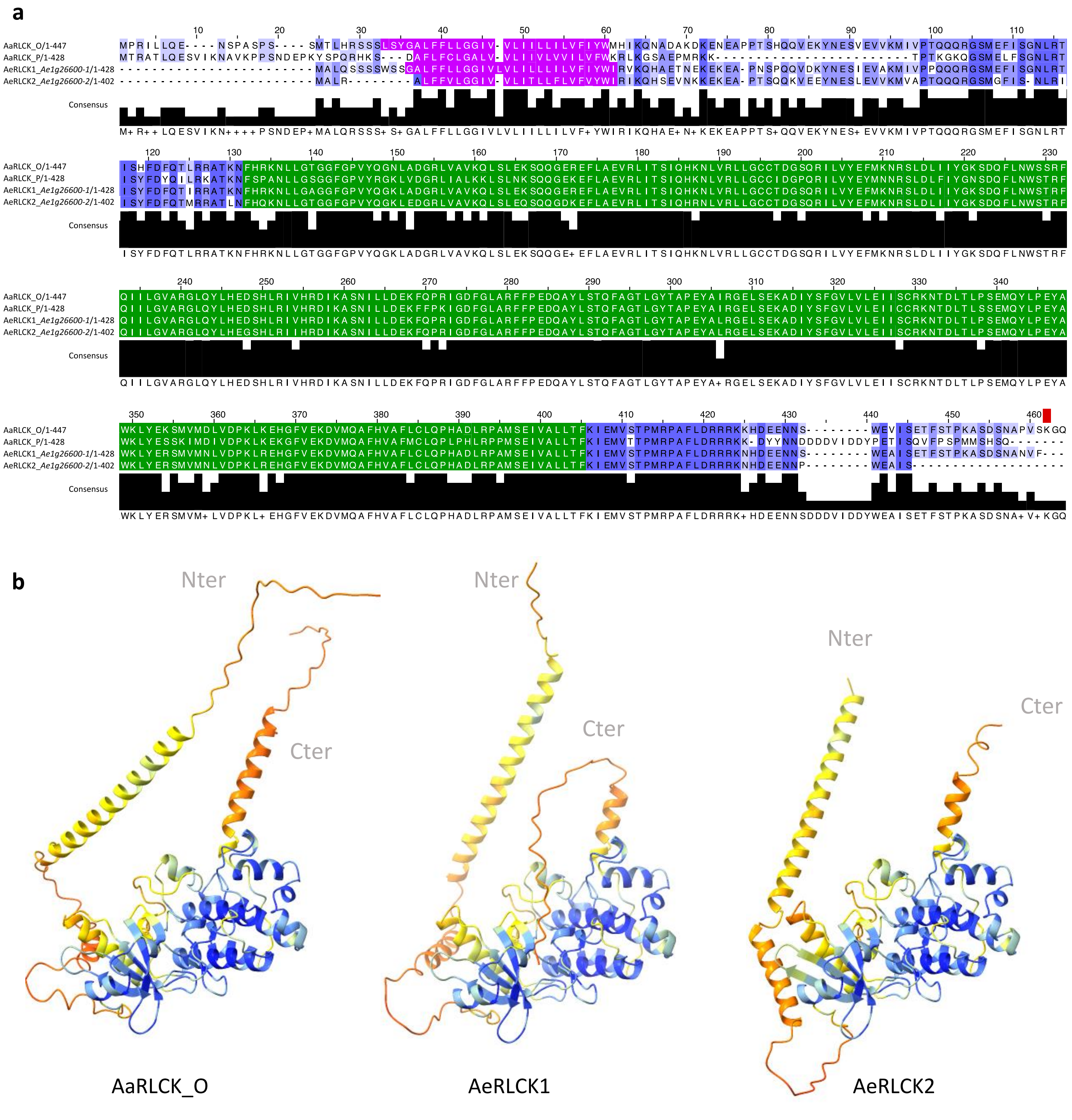


**Supplementary Fig. 11 Alignment and structure of *Aeschynomene* RLCK proteins. a** MAFFT alignment of RLCK protein sequences from *A. evenia,* RLCK1 and RLCK2, and *A. afraspera*, RLCK_P and RLCK_O. Predicted transmembrane and kinase domains are highlighted in pink and green, respectively. The level of consensus identity is represented by varying intensities of blue shading. Differences in blue intensity indicate the level of conservation. A light blue indicates low conservation while a dark blue indicates a high degree of homology. **b** Alphafold predicted structure of AaRLCK_O, AeRLCK1 and AeRLCK2. The top-ranked prediction model is displayed. The pLDDT indicates the model's confidence in the local accuracy of the predicted structure. The pLDDT is shown as a range of colours where warmer colours show poorer prediction than cooler colours. Nter and Cter domains were poorly predicted. The kinase domain is well predicted and its structure is conserved between RLCKs variants.

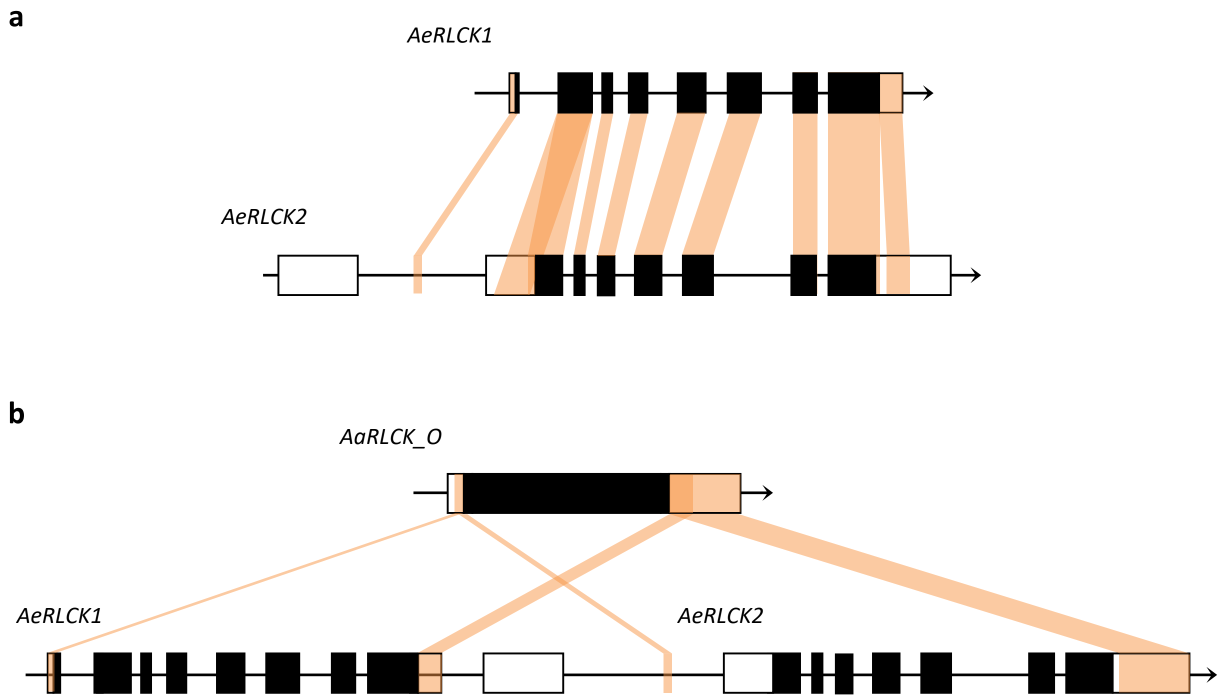


**Supplementary Fig. 12 Comparison of gene structure between *AaRLCK_O, AeRLCK1* and *AeRLCK2.*** **a** Comparison of the genomic structures of *AeRLCK1* and *AeRLCK2* located at the Ae01g26600 locus in *A. evenia*. **b** The *AaRLCK_O* transcript from *Aeschynomene afraspera* is also compared with *A. evenia AeRLCK1* and *AeRLCK2*. Black bars represent exons, while white bars indicate UTR regions. Sequence conservation blocs between the different regions are indicated by orange shaded areas.

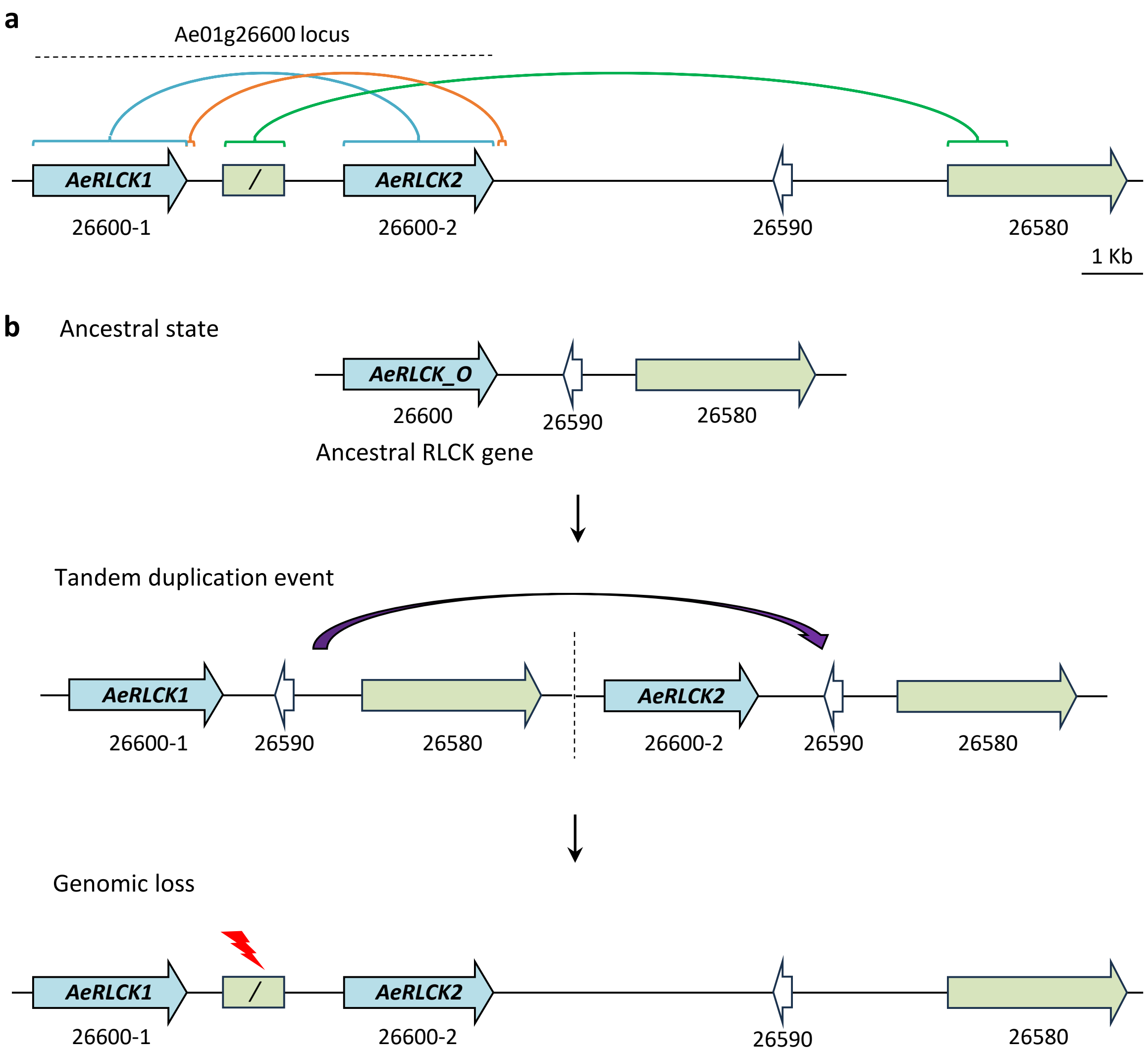


**Supplementary Fig. 13 Model of gene duplication at the *AeRLCK2* locus.** **a** Genomic region of chromosome Ae01 containing the Ae01g26600 locus is shown at scale. Blue, orange and green square brackets indicate that three conserved sequence blocks are present in this locus. **b** Schematic illustration of presumed *AeRLCK* locus evolution. Initial genomic structure showing an ancestral version of Ae01g26600 and two adjacent genes, Ae01g26590 and Ae01g26580, on chromosome Ae01 (top). Transient genomic structure following tandem duplication of the locus (middle). The promoter gene sequence of *AeRLCK2* has undergone rearrangements or accumulated mutations, resulting in significant modifications in the current genomic structure (bottom).

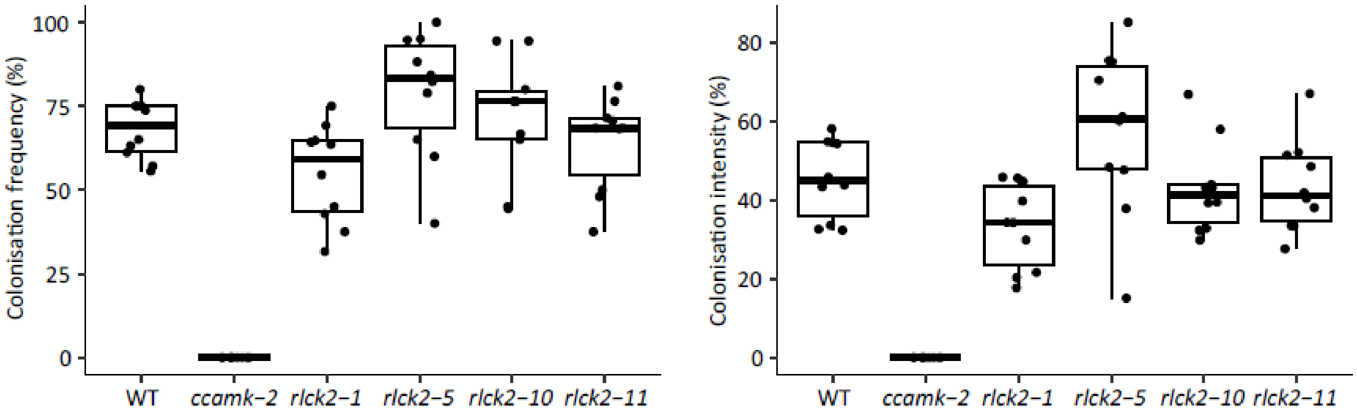


**Supplementary Fig. 14 Mycorrhizal phenotype of the *rlck2* mutants.** Box plots show the colonisation frequency and intensity, both expressed as percentages, in 6 weeks post-inoculation WT, *ccamk-2* and *rlck2* mutant lines. n=10 plants per line.

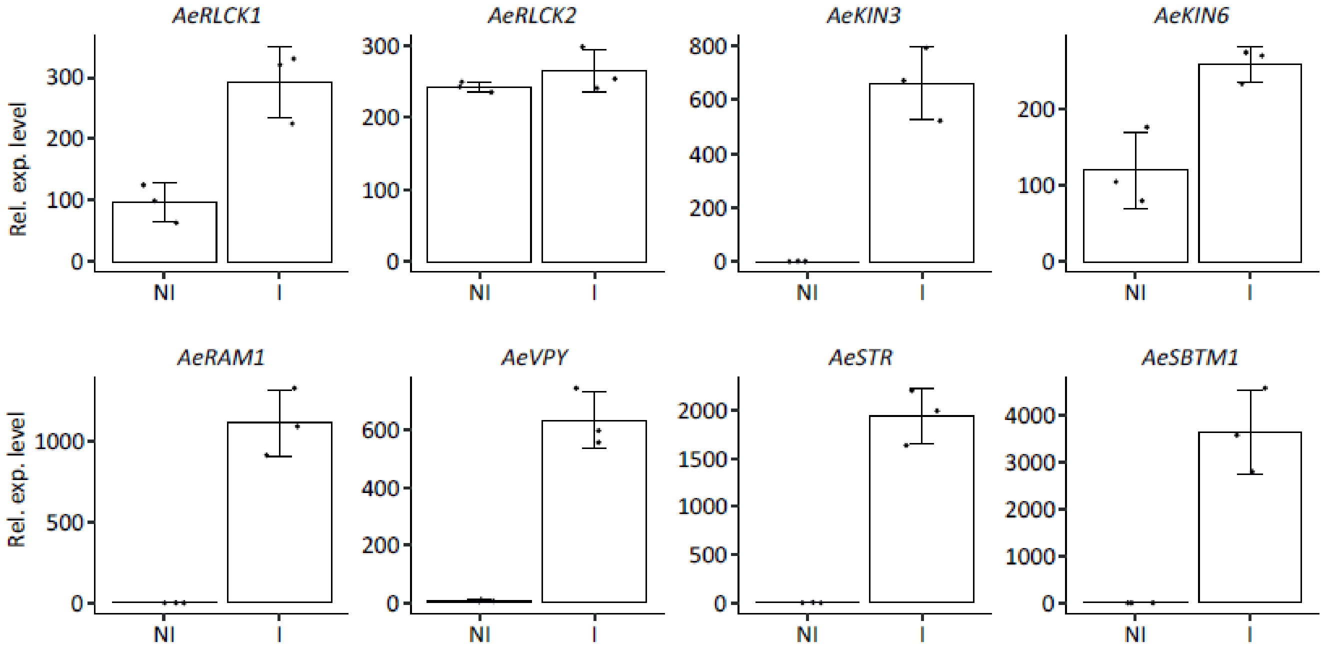


**Supplementary Fig. 15 Gene expression levels during AM in *Aeschynomene evenia*.** RNASeq data of AM-induced gene expression in non-inoculated (NI) and inoculated (I) conditions, 6 weeks after inoculation with spores of *Rhizophagus irregularis*. WT *A. evenia* samples produced in biological triplicates.

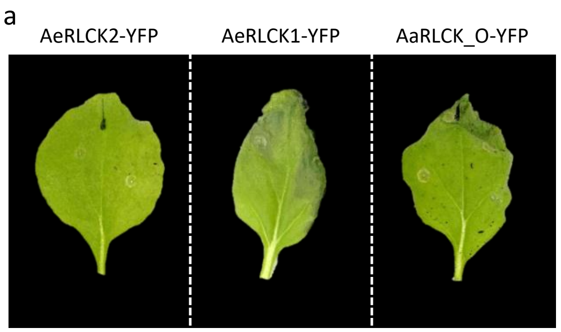


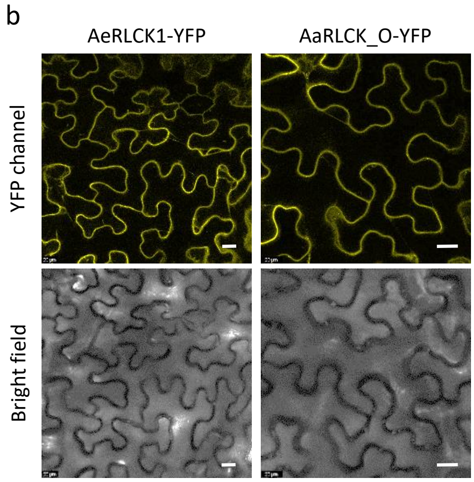


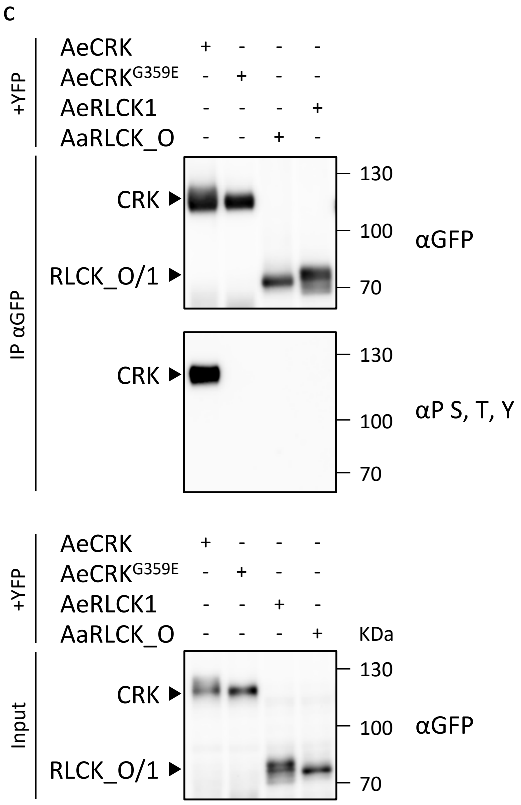


**Supplementary Fig. 15 Comparative analysis of *AeRLCK1* and *AaRLCK_O* in *Nicotiana benthamiana* leaves. a** Transient expression of *AeRLCK1*-YFP and *AaRLCK_O-YFP* triggers cell death in *N. benthamiana leaves* at 8 days post-inoculation (dpi). **b** Confocal microscopy showing the plasma membrane localization of AeRLCK1-YFP and AaRLCK_O-YFP in *N. benthamiana* leaf cells. Scale bar: 20 µm. **c** AeRLCK1 and AaRLCK_O have no autophosphorylation activity *in planta*. The YFP-tagged constructs were transiently expressed in *N. benthamiana* leaves. The proteins were immunopurified with αGFP agarose magnetic beads and the phosphorylation status was assessed using antibodies (αP S, T, Y) that recognize phosphorylated threonine, tyrosine, and serine residues. The upper panel shows the immunopurified proteins and the lower panel shows the input. In this experiment, AeCRK and AeCRK$^{G359E}$ were used as positive and negative controls, respectively.

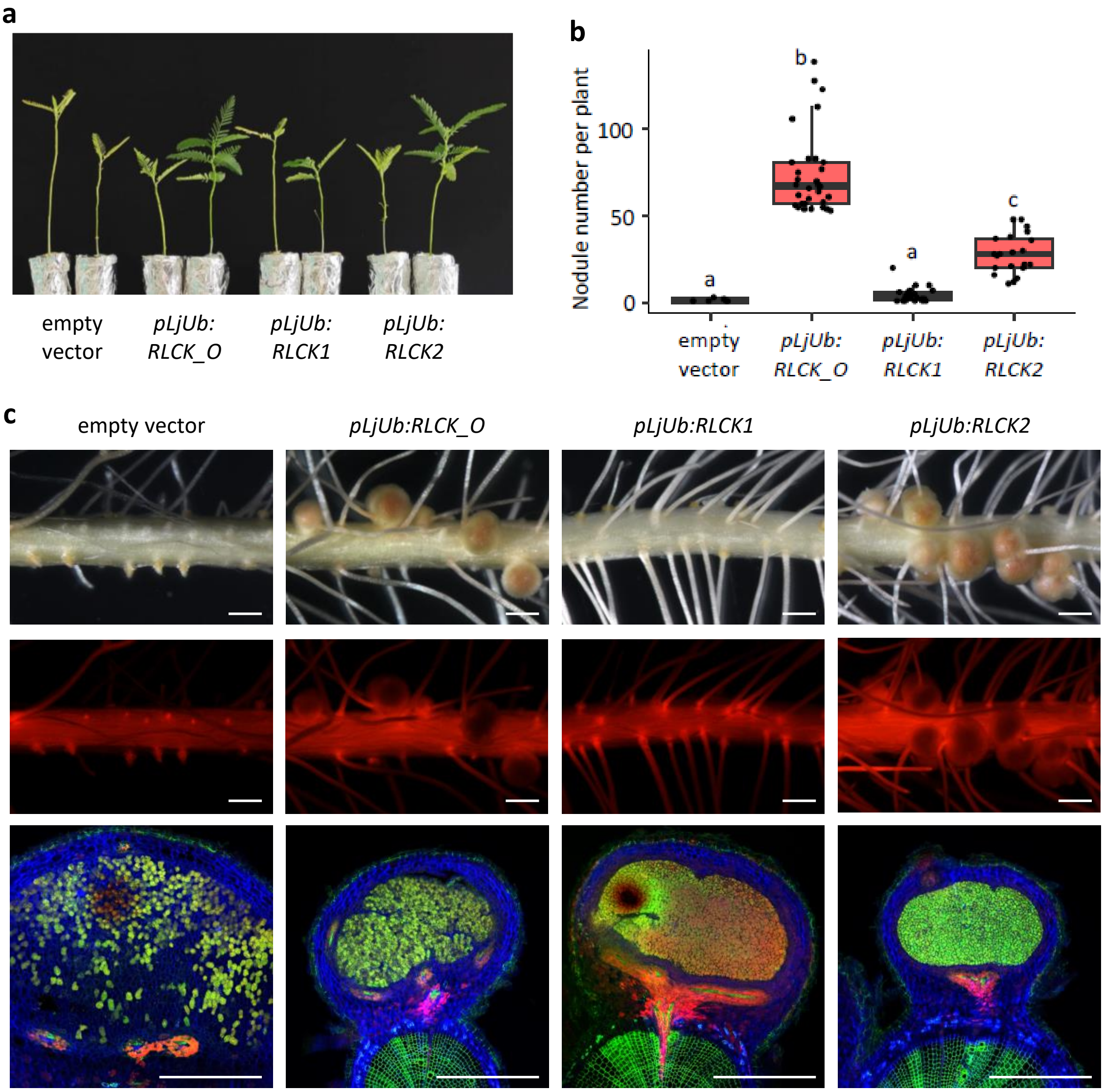


**Supplementary Fig. 16 *A. evenia rlck2-11* mutant trans-complementation of root nodulation using pLjUb**. Hairy roots of *A. evenia rlck2* mutant plant untransformed (UT) or transformed with the empty vector (EV) containing the DsRed marker, *pLjUb:RLCK_O*, *pLjUb:RLCK1* or *pLjUb:RLCK2* were generated and their nodulation phenotype was evaluated 21 days post-inoculation with *Bradyrhizobium* ORS278 strain. **a** Image showing the aerial phenotype of non-inoculated (left) and inoculated (right) plants. **b** Number of pink nodules formed on representative plants expressing specified constructs. Points represent individual plants. Letters indicate significant differences between samples, as determined by analysis of variance (Kruskal-Wallis) and post-hoc analysis (Dunn's test), p<0,05. **c** Roots were observed under brightfield (top) and a DsRed filter (middle) using a stereomicroscope. Cross-sections of nodules stained with SYTO 9, propidium iodide and calcofluor were observed under confocal microscope (bottom). Images represent the main root (top and middle) and nodule (bottom) phenotypes observed. Scale bars: 1 mm (top and middle), 0,5 mm (bottom).

### Supplementary Tables

**Supplementary Table 1. *Aeschynomene* species used in this study.**

| Species name | Accession | Symbiotic group | Ploidy level | Origin | Seedbank |
|---|---|---|---|---|---|
| *A. afraspera* | LSTM1 | ND | 8x | Senegal | LSTM |
| *A. americana* | LSTM281 | ND | 2x | Madagascar | LSTM |
| *A. ciliata* | IRRI 013078 | NI | 2x | Colombia | IRRI |
| *A. deamii* | LSTM24 | NI | 2x | Mexico | LSTM |
| *A. denticulata* | IRRI0 13003 | NI | 2x | Brazil | IRRI |
| *A. evenia* spp. *evenia* | CIAT22838 | NI | 2x | Malawi | CIAT |
| | PI225551 | | | Zambia | USDA |
| *A. evenia* spp. *serrulata* | IRFL6945 | NI | 2x | USA | USDA |
| *A. filosa* | CIAT22466 | NI | 2x | Mexico | CIAT |
| *A. fluminensis* | LSTM17 | ND | 2x | Brazil | LSTM |
| *A. montevidensis* | LSTM131 | ND | 2x | Uruguay | LSTM |
| *A. patula* | CPI 052332 | ND | 2x | Madagascar | AusPGRIS |
| *A. rudis* | Matt Lavin #82 | NI | 2x | USA | LSTM |
| *A. scabra* | LSTM26 | NI | 2x | Mexico | LSTM |
| *A. selloi* | CPI104040 | NI | 2x | Argentina | AusPGRIS |
| *A. sensitiva* | LSTM28 | NI | 2x | Senegal | LSTM |
| *A. sp328* | CIAT8499 | NI | 2x | Brazil | CIAT |
| *A. tambacoundensis* | LSTM60 | NI | 2x | Senegal | LSTM |
| *S. semperflorens* | CIAT9844 | ND | 2x | Brazil | CIAT |

N.B: *S. sempeflorens* is included as the single species of the genus *Soemmeringia* that is allied to the genus *Aeschynomene*.

Symbiotic group: ND for Nod-dependent, NI for Nod-independent.

**Supplementary Table 2. Phenotypic, genetic and molecular data on the *Aeschynomene evenia* nodulation mutants.**

| Gene | Mutant | Allele | Mutation effect | | Genetic determinism | | % of BN phenotype in F2 mutant plants | F2 mutant sequencing | % mutant allele |
|---|---|---|---|---|---|---|---|---|---|
| | | | Nucleotide change | Amino-acid change | F2 Nod+:Nod- | (*, P> 0.05) | | | |
| *AeRLCK2* (Ae01g26600) | A40 | *rlck2-1* | $G_{1932}$ to A | $E_{278}$ to K | F2 458:164 | 3:1* | 14% | MBS | 90% |
| | D22 | *rlck2-2* | $G_{516}$ to A | $G_{108}$ to E | F2 431:162 | 3:1* | 6% | MBS | 97% |
| | E27 | *rlck2-3* | $C_{857}$ to A | $L_{170}$ to F | F2 415:159 | 3:1* | 3% | PCR | - |
| | L10 | *rlck2-4* | $G_{1957}$ to A | $G_{286}$ to E | F2 466:151 | 3:1* | 6,60% | MBS | 100% |
| | L22 | *rlck2-5* | $G_{1066}$ to A | Splice site | F2 418:157 | 3:1* | 4,50% | PCR | - |
| | L24 | *rlck2-6* | $G_{525}$ to A | $G_{110}$ to E | F2 496:163 | 3:1* | 12,30% | MBS | 97% |
| | L31 | *rlck2-7* | $G_{1129}$ to A | $G_{207}$ to R | F2 379:118 | 3:1* | 15% | MBS | 100% |
| | L37 | *rlck2-8* | $G_{1951}$ to A | $S_{284}$ to N | F2 398:138 | 3:1* | 20% | MBS | 100% |
| | N10 | *rlck2-9* | $G_{776}$ to A | $E_{143}$ to K | F2 137:58 | 3:1* | 8% | PCR | - |
| | V21 | *rlck2-10* | $G_{1920}$ to A | $G_{274}$ to R | F2 418:153 | 3:1* | 6,30% | MBS | 96% |
| | X30 | *rlck2-11* | $C_{749}$ to T | $S_{131}$ to STOP | F2 394:155 | 3:1* | 14% | MBS | 87% |
| | AE4 | *rlck2-12* | $G_{1292}$ to A | G261 to E | F2 245:72 | 3:1* | 6% | PCR | - |
| *AeCCamK* (Ae08g13330) | J31 | *ccamk-2* | $G_{2405}$ to A | S341 to STOP | - | - | - | - | - |

Genetic determinism: *Significance according to a $\chi^2$ test (α=5%).
when x2 , 3.84.

BN: Big Nodule, PCR: PCR amplication and sequencing, MBS: mapping-by-sequencing.

-: not determined.

**Supplementary Table 3. Allelism analysis of the *Aeschynomene evenia rlck2* mutants.**

| Gene | Crossing (♂ x ♀) | n F1 plants (n pods) | F1 phenotype |
|---|---|---|---|
| *AeRLCK2* (Ae01g26600) | A40 x L10 | 5(1) | Nod- |
| | A40 x V21 | 6(1) | Nod- |
| | D22 x L10 | 3(1) | Nod- |
| | D22 x M11 | 4(1) | Nod- |
| | E27 x V21 | 18(3) | Nod-/1BN |
| | L10 x V21 | 6(1) | Nod- |
| | L22 x V21 | 18(3) | Nod- |
| | L24 x L10 | 10(2) | Nod- |
| | L24 x V21 | 3(1) | Nod- |
| | L31 x X30 | 7(1) | Nod-/1BN |
| | L37 x V21 | 23(3) | Nod- |
| | N10 x V21 | 12(2) | Nod- |
| | N10 x X30 | 6(1) | Nod- |
| | V21 x E27 | 6(2) | Nod- |
| | V21 x L10 | 9(1) | Nod- |
| | X30 x L10 | 1(1) | Nod- |
| | X30 x L31 | 15(3) | Nod-/2BN |
| | X30 x N10 | 6(1) | Nod- |
| | X30 x V21 | 7(1) | Nod- |
| | X30 x AE4 | 10(2) | Nod- |

Nod-: no nodule phenotype, BN: Big Nodule phenotype.

**Supplementary Table 4. Functional complementation test for nodulation with *AeRLCK2.***

| ***A. evenia* line** | **Transformation construct** | **Transformed plants** | **Nodulated plants** | **Nodules/ nodulated plant*** |
|---|---|---|---|---|
| *rlck2-11* | pUB-GW-GFP | 14 | 0 | 0 |
| *rlck2-11* | pUB-AeRLCK2-GFP | 15 | 9 | 17.4 ± 11.8 |
| WT | not transformed | 5 | 5 | 19.6 ± 5.9 |

* Values correspond to mean nodule number per nodulated plant ± standard deviation.

**Supplementary Table 5. Summary of Illumina transcriptome sequencing and assembly for *Aeschynomene afraspera*.**

| Samples | Total reads | Total length | Number of contigs | N50 | Number of genes |
|---|---|---|---|---|---|
| A. afraspera root #1 | 60,214,658 | 7,526,832,250 | - | - | 49,981 |
| A. afraspera root #2 | 68,854,420 | 8,606,802,500 | - | - | 50,067 |
| A. afraspera root #3 | 94,432,112 | 11,804,014,000 | - | - | 50,045 |
| A. afraspera nodules 4dpi ORS285 #1 | 105,844,382 | 13,230,547,750 | - | - | 49,52 |
| A. afraspera nodules 4dpi ORS285 #2 | 60,283,806 | 15,070,951,500 | - | - | 49,907 |
| A. afraspera nodules 4dpi ORS285 #2 | 117,268,432 | 14,658,554,000 | - | - | 49,858 |
| A. afraspera nodules 8dpi ORS285 #1 | 82,378,552 | 10,297,319,000 | - | - | 50,133 |
| A. afraspera nodules 8dpi ORS285 #2 | 87,745,420 | 10,968,177,500 | - | - | 50,255 |
| A. afraspera nodules 8dpi ORS285 #3 | 78,019,448 | 9,752,431,000 | - | - | 50,261 |
| Total | 755,041,230 | 101,915,629,500 | 50,437 | 1,979 | 50,438 |

#### Supplementary Table 6. Analysis of the list of 138 AM conserved genes.

| Mt Gene v4[a] | Annotation | Gene | Family | Functional category | OrthoGroup[b] | Ae Gene[c] | Gene analysis in legumes[d] |
|---|---|---|---|---|---|---|---|
| Medtr8g022270 | ABC transporter | ABCB12 | ABCB12 | Transporter | OG0011668 | Ae06g23390 | |
| Medtr3g093430 | ABC transporter | ABCB20 | ABCB20 | Transporter | OG0014895 | Ae08g07060 | |
| Medtr3g111900 | AMP-binding enzyme | AMP1 | AMPa | Lipid metabolism | OG0017523 | Ae08g02730 | |
| Medtr2g016730 | AP2 domain protein | AP2c | AP2C | Transcription | OG0014406 | Ae03g26240<br>Ae04g04370 | |
| Medtr8g032610 | AP2/ERF domain transcription factor | AP2d | AP2D | Transcription | OG0013970 | Ae02g03600<br>Ae06g31890 | |
| Medtr1g090420 | phytocyanin | BCP1 | BCP | Redox enzyme | | - | |
| Medtr7g117580 | ion channel protein | CASTOR | CAS | Signaling | OG0002125 | Ae04g09220 (AePOLLUX)<br>**Ae06g11380 (AeCASTOR)** | |
| Medtr6g043700 | citrate-binding protein | CBP1 | CBP | Defense | OG0007822 | Ae05g27930 | |
| Medtr3g110195 | 9-cis-epoxycarotenoid dioxygenase | CCD8b | CCD | Secondary metabolism | OG0003922 | Ae08g12010, Ae08g12020 | Legume tandem gene duplication |
| Medtr1g033360 | cytochrome protein b561 | CYTb1 | CYTB561 | Redox enzyme | OG0018523 | - | |
| Medtr6g034940 | cytochrome P450 | CYT733A1 | CYT733A1 | Secondary metabolism | | Ae05g28140 | |
| Medtr4g097510 | short-chain dehydrogenase/reductase | DHY1 | DHY | Lipid metabolism | OG0018890 | Ae01g04760 | |
| Medtr5g030920 | nodulation receptor kinase | DMI2 | DMI2 | Signaling | OG0005446 | Ae03g06650 (AeSYMRK1)<br>**Ae05g01060 (AeSYMRK2)** | |
| Medtr1g017910 | protein binding protein | EXO70I | EXO70 | Exocytosis | OG0014739 | **Ae07g09130**<br>Ae08g04110 | |
| Medtr1g109110 | acyl-(acyl carrier protein) thioesterase | FatM | FATM | Lipid metabolism | OG0016085 | Ae09g25530 | |
| Medtr1g069725 | GRAS family transcription factor | TF72 | GRAS | Transcription | OG0015322 | Ae02g08200 | |
| Medtr5g076900 | glutathione S-transferase tau | GST1 | GST | Redox enzyme | OG0017182 | Ae03g06920 | |
| Medtr5g020960 | heavy-metal-associated domain protein | HMAD1 | HMAD | Unknown | OG0015100 | Ae05g04750 | |
| Medtr4g104750 | hypothetical protein | HYP3 | HYP3 | Unknown | OG0012602 | Ae01g19590 | |
| Medtr2g104800 | hypothetical protein | HYP5a | HYP5a | Unknown | OG0008590 | Ae06g09830<br>**Ae10g14590** | |
| Medtr8g069400 | nucleotide-diphospho-sugar transferase | HYP5c | HYP5c | Unknown | OG0005988 | Ae03g06090 | |
| Medtr8g040940 | hypothetical protein | HYP6 | HYP6 | Unknown | OG0015844 | Ae08g16290 | |
| Medtr5g018610 | hypothetical protein | HYP7 | HYP7 | Unknown | OG0016132 | - | |
| Medtr2g012790 | kelch repeat protein | KELCH1 | KELCH | Unknown | OG0014388 | Ae04g06260 | |
| Medtr3g104900 | serine-threonine protein kinase | KIN5 | KINF | Signaling | OG0014184 | Ae07g10560 | |
| Medtr5g033490 | LysM type receptor kinase | LYK10 | LYK | Signaling | OG0010887 | -<br>Ae03g06000 | |
| Medtr3g115940 | MLO-like protein | MLO | MLO | Defense | OG0016001 | Ae08g01190 | |
| Medtr3g088855 | protein kinase | KIN4 | kinE1pair | Signaling | OG0013015 | Ae08g08730 | |
| **Medtr6g007690** | protein kinase | KIN6 | kinG1pair | Signaling | OG0011910 | Ae03g05490<br>**Ae05g36220** | |
| **Medtr7g116650** | protein kinase domain protein | KIN3 | kinG2pair | Signaling | OG0015837 | Ae06g09820 | |
| Medtr4g129010 | protein kinase | KIN2 | kinHpair | Signaling | OG0015303 | Ae01g25530 | |
| Medtr0021s0370 | signal peptidase I | TAU | TAU | Redox enzyme | OG0013049 | Ae06g28700 | |
| Medtr1g112940 | protein phosphatase 2A regulatory B subunit | PP2A | PP2A | Signaling | OG0015848 | Ae09g27060 | |
| Medtr1g028600 | inorganic phosphate transporter | PT4 | PT4 | Transporter | OG0015195 | Ae08g08360 | |
| Medtr7g027190 | GRAS family transcription factor | RAM1 | RAM1 | Transcription | OG0016905 | Ae06g18380 | |
| Medtr1g040500 | glycerol-3-phosphate acyltransferase | RAM2 | RAM2 | Lipid metabolism | OG0010966 | Ae09g18120 | |
| Medtr5g020810 | replication factor C subunit 3 | RFCa | RFCa | Unknown | OG0014959 | Ae05g04890 | |
| Medtr3g118160 | replication factor C subunit | RFCb | RFCb | Unknown | OG0015288 | Ae08g00170 | |
| Medtr7g087500 | Sec14p-like phosphatidylinositol transfer | Sec14p | SEC | Exocytosis | OG0015001 | Ae06g14940 | |
| Medtr5g030910 | ABC transporter | STR2 | STR2 | Transporter | OG0014512 | Ae05g01070 | |
| Medtr8g107450 | ABC transporter | STR | STR | Transporter | OG0013556 | Ae05g35200 | |
| Medtr2g088700 | syntaxin | SYN | SYN | Exocytosis | OG0004618 | Ae10g20750 | |
| Medtr4g104020 | GRAS family transcription factor | RAD1 | RAD1 | Transcription | OG0016202 | Ae01g19180 | |
| Medtr6g027840 | penetration and arbuscule morphogenesis protein | VPY | VPY | Exocytosis | OG0007003 | **Ae05g16930**<br>Ae06g15540 | |
| Medtr2g091210 | DUF538 domain protein | | 538 | Unknown | OG0009998 | Ae04g04800 | |
| Medtr2g091215 | DUF538 domain protein | | 538 | Unknown | | **Ae10g22750** | |
| Medtr2g098490 | AMP-dependent synthetase and ligase | | ampB | Lipid metabolism | OG0009981 | Ae04g32270 | |
| Medtr4g066130 | AMP-dependent synthetase and ligase | | ampB | Lipid metabolism | | | |

#### Supplementary Table 6 following. Analysis of the list of 138 AM conserved genes.

| Mt Gene v4[a] | Annotation | Gene | Family | Functional category | OrthoGroup[b] | Ae Gene[c] | Gene analysis in legumes[d] |
|---|---|---|---|---|---|---|---|
| Medtr1g036410 | ammonium transporter | AMT2-5 | amt | Transporter | OG0007212 | Ae04g35250<br>Ae08g17090 | |
| Medtr7g115050 | ammonium transporter | AMT2-4 | amt | Transporter | | Ae06g09140 | |
| Medtr4g082345 | AP2/ERF domain transcription factor | | ap2a | Transcription | OG0018374 | Ae03g03910 | |
| Medtr6g012970 | AP2/ERF domain transcription factor | | ap2a | Transcription | OG0015722 | Ae05g37610 | |
| Medtr7g011630 | AP2 domain protein | | ap2a | Transcription | | | |
| Medtr6g011490 | AP2 domain transcription factor | | ap2b | Transcription | OG0002420 | Ae05g37030 | |
| Medtr7g009410 | AP2 domain transcription factor | ERF1 | ap2b | Transcription | | Ae06g37890 | |
| Medtr7g009430 | AP2 domain transcription factor | | ap2b | Transcription | | | |
| Medtr8g468920 | AP2 domain transcription factor | | ap2b | Transcription | | Ae03g04600 | |
| Medtr2g081600 | CCAAT-binding transcription factor | Cbf1 | cbf | Transcription | OG0014916 | - | |
| Medtr2g081630 | CCAAT-binding transcription factor | Cbf2 | cbf | Transcription | | | |
| Medtr8g092440 | GRAS family transcription factor | DIP1b | dip1 | Transcription | OG0014874 | Ae03g28880 | |
| Medtr8g093070 | GRAS family transcription factor | DIP1 | dip1 | Transcription | | | |
| Medtr2g008520 | DnaJ domain protein | | dnaj | Stress response | OG0004610 | Ae04g08160 | |
| Medtr2g008540 | DnaJ domain protein | | dnaj | Stress response | | | |
| Medtr4g094275 | chaperone DnaJ-domain protein | | dnaj | Stress response | | Ae03g22010 | |
| Medtr8g074560 | GDSL-like lipase/acylhydrolase | | gdsl | Lipid metabolism | OG0009476 | Ae01g01340 | Legume tandem gene duplication |
| Medtr8g074570 | GDSL-like lipase/acylhydrolase | | gdsl | Lipid metabolism | | Ae01g01350 | |
| Medtr8g074580 | GDSL-like lipase/acylhydrolase | | gdsl | Lipid metabolism | | | |
| Medtr2g086620 | germin-like protein | | ger | Redox enzyme | OG0011772 | Ae10g21600 | |
| Medtr2g086630 | germin-like protein | | ger | Redox enzyme | | | |
| Medtr2g086640 | germin-like protein | | ger | Redox enzyme | | | |
| Medtr4g052770 | germin-like protein | | ger | Redox enzyme | | Ae04g28380 | |
| Medtr4g052780 | germin-like protein | | ger | Redox enzyme | | | |
| Medtr1g062970 | heparan-alpha-glucosaminide N-acetyltransferase | | hep | Lipid metabolism | OG0000947 | Ae02g22310 | |
| Medtr7g102840 | protein D8Ertd354e | | hep | Lipid metabolism | | Ae06g02920 | |
| Medtr7g102870 | heparan-alpha-glucosaminide N-acetyltransferase | | hep | Lipid metabolism | | | |
| Medtr3g099200 | hypothetical protein | | hyp | Redox enzyme | OG0004573 | Ae08g23990 (AeEPP1) | |
| Medtr4g102890 | hypothetical protein | | hyp | Redox enzyme | | Ae01g18860 | |
| Medtr4g104060 | hypothetical protein | | hyp | Redox enzyme | | | |
| Medtr3g467150 | hypothetical protein | | hyp2 | Unknown | OG0013890 | Ae01g11360 | |
| Medtr5g075400 | hypothetical protein | | hyp2 | Unknown | | | |
| Medtr1g069620 | hypothetical protein | | hyp4 | Unknown | OG0015808 | Ae02g08320 | |
| Medtr2g022500 | hypothetical protein | | hyp4 | Unknown | - | - | |
| Medtr5g026850 | cyclops protein | IPD3 | ipd3 | Transcription | OG0012745 | Ae05g02230 | |
| Medtr8te071320 * | cyclops protein | IPD3b | ipd3 | Transcription | | | |
| Medtr2g023150 | protein kinase | | kinA | Signaling | OG0015589 | Ae06g24770 | |
| Medtr3g078250 | protein kinase | | kinA | Signaling | OG0004099 | **Ae08g22270 (AeRINRK1)**<br>Ae01g20350 (AeRINRK2) | |
| Medtr1g098300 | cyclin-dependent kinase | | kinD | Signaling | OG0013175 | Ae09g21070 | |
| Medtr8g092290 | cyclin-dependent kinase | | kinD | Signaling | OG0014359 | Ae03g28630 | |
| Medtr2g088610 | late embryogenesis abundant protein | | lea | Unknown | OG0007704 | Ae10g20760 | |
| Medtr2g088680 | late embryogenesis abundant protein | | lea | Unknown | | | |
| Medtr4g046787 | late embryogenesis abundant protein | | lea | Unknown | | | |
| Medtr4g046803 | late embryogenesis abundant protein | | lea | Unknown | | | |
| Medtr4g046830 | transmembrane protein | | lea | Unknown | | Ae04g27020 | |
| Medtr5g005950 | late embryogenesis abundant protein | | lea | Unknown | | | |
| Medtr3g086430 | ABCB transporter , MDR family | | mdr1 | Transporter | OG0008675 | Ae03g13920 | |
| Medtr4g081190 | ABCB transporter , MDR family | | mdr1 | Transporter | | Ae04g24770 | |
| Medtr5g019040 | Nod-factor receptor 5 | NFP | nfp | Signaling | OG0011134 | Ae05g05580 | |
| Medtr8g078300 | Nod-factor receptor 5 | LYR1 | nfp | Signaling | | | |
| Medtr8g078320 | Nod-factor receptor 5 | | nfp | Signaling | | | |
| Medtr8g078340 | Nod factor receptor | | nfp | Signaling | | | |
| Medtr8g078360 | Nod-factor receptor 5 | | nfp | Signaling | OG0046535 | - | |

#### Supplementary Table 6 following. Analysis of the list of 138 AM conserved genes.

| Mt Gene v4[a] | Annotation | Gene | Family | Functional category | OrthoGroup[b] | Ae Gene[c] | Gene analysis in legumes[d] |
|---|---|---|---|---|---|---|---|
| Medtr2g017750 | nitrate transporter | | ntr1 | Transporter | OG0010428 | Ae04g04070 | |
| Medtr8g087780 | nitrate transporter | | ntr1 | Transporter | | Ae03g26550 | |
| Medtr8g087810 | nitrate transporter | | ntr1 | Transporter | | | |
| Medtr3g051230 | cytochrome P450 | | p450 | Secondary metabolism | - | - | |
| Medtr5g092150 | cytochrome P450 | | p450 | Secondary metabolism | OG0012010 | Ae07g02620 | |
| Medtr3g064080 | protein kinase | | kinBpair | Signaling | OG0000618 | Ae01g16400 | Legume gene cluster |
| Medtr3g064090 | protein kinase domain protein | | kinBpair | Signaling | | Ae01g16410, Ae01g16430, Ae01g16440 | |
| Medtr3g064110 | protein kinase domain protein | | kinBpair | Signaling | | | |
| Medtr2g038675 | protein kinase | | kinCpair | Signaling | OG0004144 | - | Nod-independent *Aeschynomene* tandem gene duplication |
| Medtr4g126930 | protein kinase | | kinCpair | Signaling | | Ae01g26600 - AeRLCK1 & AeRLCK2 | |
| Medtr0262s0020 | plastocyanin-like domain protein | | pcl | Redox enzyme | - | - | |
| Medtr1g105120 | plastocyanin-like domain protein | | pcl | Redox enzyme | OG0002550 | Ae01g02930, Ae06g15560, Ae06g15570, Ae06g15580, Ae07g23550 | Legume gene cluster |
| Medtr1g105130 | plastocyanin-like domain protein | | pcl | Redox enzyme | | | |
| Medtr7g086090 | plastocyanin-like domain protein | | pcl | Redox enzyme | | | |
| Medtr7g086100 | plastocyanin-like domain protein | | pcl | Redox enzyme | | | |
| Medtr7g086140 | plastocyanin-like domain protein | | pcl | Redox enzyme | | | |
| Medtr7g086190 | plastocyanin-like domain protein | | pcl | Redox enzyme | | | |
| Medtr7g086200 | plastocyanin-like domain protein | | pcl | Redox enzyme | | | |
| Medtr7g086220 | plastocyanin-like domain protein | | pcl | Redox enzyme | | | |
| Medtr7g086230 | plastocyanin-like domain protein | | pcl | Redox enzyme | | | |
| Medtr7g086280 | plastocyanin-like domain protein | | pcl | Redox enzyme | | | |
| Medtr7te086160 | plastocyanin-like domain protein | | pcl | Redox enzyme | | | |
| Medtr0027s0260 | class III chitinase | | qi | Defense | OG0000300 | Ae03g04310, Ae03g04340, Ae06g09870, Ae06g09910, Ae09g21530, Ae09g21550 | Legume gene clusters |
| Medtr5g043550 | class III chitinase | | qi | Defense | | | |
| Medtr7g116350 | class III chitinase | | qi | Defense | | | |
| Medtr8g055940 | class III chitinase | | qi | Defense | | | |
| Medtr8g467650 | class III chitinase | | qi | Defense | | | |
| Medtr1g023770 | rhicadhesin receptor | | rhi | Redox enzyme | - | | |
| Medtr2g030855 | rhicadhesin receptor | | rhi | Redox enzyme | - | | |
| Medtr2g030865 | rhicadhesin receptor | | rhi | Redox enzyme | - | | |
| Medtr2g030895 | rhicadhesin receptor | | rhi | Redox enzyme | OG0014679 | Ae04g09850 | |
| Medtr2g031270 | rhicadhesin receptor | | rhi | Redox enzyme | | | |
| Medtr2g031300 | rhicadhesin receptor | | rhi | Redox enzyme | | | |
| Medtr3g020800 | subtilisin inhibitor 1 | | sin | Defense | OG0014420 | Ae08g16310, Ae08g16320, Ae08g16360, Ae08g16380, Ae08g16390 | Legume gene cluster |
| Medtr3g020880 | subtilisin inhibitor 1 | | sin | Defense | | | |
| Medtr3g020930 | subtilisin inhibitor | | sin | Defense | | | |
| Medtr3g020970 | subtilisin inhibitor | | sin | Defense | | | |

[a] The list of AMS conserved genes was initially opbtained by Bravo et al. (2016)

[b] The groups of orthologous genes were obtained by ORTHOFINDER2 in Quilbé_2021

[c] When several genes are present in *A. evenia,* orthologs are in bold.

[d] Gene analysis was conducted by eximining ORTHIFINDER-derived phylogenic trees and through genome examination in AeschynomeneBase.

Mt: *M. truncatula*, Ae: *A. evenia*.

**Supplementary Table 7. Summary of Illumina transcriptome sequencing for *Aeschynomene evenia* CIAT22838**

| Samples | Total reads | Total length | Total mapped | % hit | Total genes | Gene coverage |
|---|---|---|---|---|---|---|
| Ae-myco-6wpi-NI-1 | 33,013,991 | 9,772,141,336 | 28,499,856 | 86,33 | 22,716 | 69,5% |
| Ae-myco-6wpi-NI-2 | 29,184,895 | 8,609,544,025 | 25,143,995 | 86,16 | 22,505 | 68,9% |
| Ae-myco-6wpi-NI-3 | 26,924,662 | 7,996,624,614 | 25,121,936 | 93,31 | 22,436 | 68,7% |
| Ae-myco-6wpi-I-1 | 26,927,650 | 7,970,584,400 | 22,396,078 | 83,17 | 22,601 | 69,2% |
| Ae-myco-6wpi-I-2 | 24,752,428 | 7,326,718,688 | 19,515,864 | 78,85 | 22,435 | 68,7% |
| Ae-myco-6wpi-I-3 | 23,086,500 | 6,833,604,000 | 19,752,697 | 85,56 | 22,314 | 68,3% |
| Total | 163,890,126 | 48,509,217,063 | 140,430,426 | - | - | - |

**Supplementary Table 8. Nodulation data for the cross-complementation tests of *rlck2-11* mutant.**

| Genotype | Construct | Transformed plants | Nodulated plants | Major nodulation phenotype | Nodules/nodulated plant* |
|---|---|---|---|---|---|
| *rlck2-11* | empty vector | 21 | 14 | 11/14: BN | 1.6 ± 2.0 |
| *rlck2-11* | pAeRLCK2:RLCK_O | 38 | 35 | 27/35 : WT like nodules | 27.0 ± 8.8 |
| *rlck2-11* | pAeRLCK2:RLCK1 | 37 | 24 | 20/26: BN | 2.2 ± 1.9 |
| *rlck2-11* | pAeRLCK2:RLCK2 | 44 | 41 | 30/41 : WT like nodules | 18.7 ± 7.5 |

| Genotype | Construct | Transformed plants | Nodulated plants | Major nodulation phenotype | Nodules/nodulated plant* |
|---|---|---|---|---|---|
| *rlck2-11* | empty vector | 43 | 7/43 | 36/43: Nod$^-$ roots | 1.4 ± 0.8 |
| *rlck2-11* | pLjUb:RLCK_O | 47 | 44/47 | 32/47: WT like nodules | 73.9 ± 23.2 |
| *rlck2-11* | pLjUb:RLCK1 | 49 | 30/49 | 22/30: few nodules | 4.5 ± 4.5 |
| *rlck2-11* | pLjUb:RLCK2 | 40 | 34/40 | 21/40: WT like nodues | 28.2 ± 11.4 |

| Genotype | Construct | Transformed plants | Nodulated plants | Major nodulation phenotype | Nodules/nodulated plant |
|---|---|---|---|---|---|
| WT | empty vector | 17 | 17 | 17/17: WT like nodules | 33.1 ± 13.5 |

BN: Big Nodule.

*: mean ± sd

**Supplementary Table 9.** Primer sequences used for PCR amplification, cloning and site-directed mutagenesis.

| Purpose | Approach | Primer name | Sequence (5'-3') |
|---|---|---|---|
| Mutation sequencing in *AeRLCK2* | Sanger sequencing | AeRLCK2-F1 | CACTGTTGGTCTCTGTATC |
| | | AeRLCK2-F2 | AGCAAGGAGATAAAGAATTTC |
| | | AeRLCK2-R1 | ATAAACATGATGGTGAATGG |
| | | AeRLCK2-R2 | AGGACATAGCTTCTAGTTTG |
| *RLCK* gene detection in *Aeschynomene* sp. | Sanger sequencing | RLCK-F1 | TGGAGCATTGTTCTTTSTCCT |
| | | RLCK-F2 | ATTGCTGATYCTTGTGTTCATA |
| | | RLCK-F3 | AGCATTGTTCTTTGTCCTTGG |
| | | RLCK-F4 | TGGAGTTCATTAGTGGGAACCT |
| | | RLCK-F5 | TGATTTCCAGACAATGAGAAGG |
| | | RLCK-F6 | GAATTTCTGGCAGAGGTGAGA |
| | | RLCK-R1 | GTTTTTGCGCCTCCTATCAA |
| | | RLCK-R2 | GGCCTCATTGGTGTYGAAA |
| | | RLCK-R3 | GCCTCCTATCAAGGAATGCT |
| *AeRLCK2* cloning in pUB-GW-GFP | GateWay | AeRLCK2-Fc | ATGGCATTACGACATTGTTCT |
| | | AeRLCK2-Rc | CTAGGATATGGCTTCCCAAG |
| *AeRLCK2* promoter | GoldenGate | Supplementary File 3 | |
| *AeCRK* promoter | GoldenGate | Supplementary File 3 | |
| AeCRK CDS (no stop codon) | GoldenGate | Supplementary File 3 | |
| *AaRLCK_O* CDS (no stop codon) | GoldenGate | B-RLCK0-F | GGTCTCACAAAATGCCTCGAATACTACTCC |
| | | C-RLCK0-R | GGTCTCGCACCTTGGCCTTTAGATACTGGT |
| *AaRLCK-O* CDS (stop codon) | GoldenGate | BSAI-RLCK0-bdSTOP-F | AAGGTCTCCCAAAATGCCTCGAATACTACTCCAAG |
| | | BSAI-RLCK0-bdSTOP-R | ACGGTCTCGCGTATCATTGGCCTTTAGATACTG |
| *AeRLCK1* CDS (no stop codon) | GoldenGate | B-RLCK1-F | AAGGTCTCCCAAAATGCCTCGAATACTA |
| | | C-RLCK1-R | GGTCTCGCACCGAACACATTAGCATTAGAA |
| *AeRLCK1* CDS (stop codon) | GoldenGate | BSAI-RLCK1-bdSTOP-F | AAGGTCTCCCAAAATGCCTCGAATACTATTCCA |
| | | BSAI-RLCK1-bdSTOP-R2 | ACGGTCTCGCGTATTAGAACACATTAGCATTA |
| *AeRLCK2* CDS (no stop codon) | GoldenGate | AeRLCK2-F | ACGGTCTCGCAAAATGGCATTACGAGCATTGT |
| | | AeRLCKnoSTOP-R | ACGGTCTCTCACCGGATATGGCTTCCCAAGG |
| *AeRLCK2* CDS (stop codon) | GoldenGate | BSAI-RLCK2-bdSTOP-F | AAGGTCTCCCAAAATGGCATTACGAGCATTGTT |
| | | BSAI-RLCK2-bdSTOP-R | ACGGTCTCGCGTACTAGGATATGGCTTCCCAAG |
| AeRLCK2 Kinase | | AeRLCK2-kinEcoF | CCCGAATTCGGGTTCATTAGTGGCAATCT |
| | | AeRLCK2-kinNotR | ATTTGCGGCCGCCTAGGATATGGCTTCCCAAG |
| AeRLCK2 Kin$^{G110E}$ | Site-directed mutagenesis | AeRLCK2-mutL42F | GAAACCAAGCTTGAAGAAGGTGGCTTC |
| | | AeRLCK2-mutL42R | GAAGCCACCTTCTTCAAGCTTGGTTTC |
| AeCRK Kinase | | CRKkin_F | CTTCGGACCCGTTTTCAAG |
| | | CRKkin_R | CAGCATCCCAAGAGCCTAAC |
| AeCRK KinG110E | Site-directed mutagenesis | AeCRKkin_Mut_G359E_F | GGTGGCTTCGaACCCGTTTTC |
| | | AeCRKkin_Mut_G359E_R | TTCCCCAAGCTTGGTTTC |

**Supplementary Table 10. List of genes with the primers used for RT-qPCR analysis.**

| Gene | Ae name | Primer name | Sequence | Reference |
|---|---|---|---|---|
| *AeCRK* | Ae05g12380 | AeCRK-F<br>AeCRK-R | CCCCAATAGTTCTGAGCCAACC<br>AGGAAGAGGCCATAGAGCCTTG | Quilbé et al. (2022) |
| *AeEF1a* | Ae09g20140 | AeEF1-F<br>AeEF1-R | TGCTGGTATGGTTAAGATGGTTCC<br>TTCTTCTTCTGTGCTGCCTTGG | Quilbé et al. (2022) |
| *AeENOD40* | not annotaed | AeNOD40-F<br>AeNOD40-R | CACACTTCTCCTCCATTCACTTTTC<br>TTGCCATACTTGTAGCCAAAAGC | Quilbé et al. (2022) |
| *AeNIN* | Ae07g00100 | AeNIN-F<br>AeNIN-R | CAACAGAACAAGGGGAAAGGGG<br>TAATGAGGCAGAGGCGGAAGTG | Quilbé et al. (2022) |
| *AeRAM1* | Ae06g18380 | AeRAM1-F<br>AeRAM1-R | GTAATTCCCTCGGTAATCTGTT<br>TGGCTTCCTGCTCAACTA | Quilbé et al. (2022) |
| *AeSBT* | Ae05g09230 | AeSBT-F<br>AeSBT-R | ATGAAGGAATGCCACCTCCACC<br>TGTGTGTGCCGTGTCCATTATC | Quilbé et al. (2022) |
| *AeSBTM1* | Ae05g09240 | AeSBTM1-F<br>AeSBTM1-R | TGATATAGGCGGAGGAGT<br>ACATGAACAGCAGGAAGTA | Quilbé et al. (2022) |
| *AeSTR* | Ae05g35200 | AeSTR-F<br>AeSTR-R | TTCCTCGTATGTCATATCTTCA<br>GCATTGGTTGTGATAAGTGA | Quilbé et al. (2022) |
| *AeSYMREM1* | Ae03g30480 | AeSYMREM1-F<br>AeSYMREM1-R | TGATGATGCTGCTGATGA<br>TTTGGGTTGTGAAGAAGTG | Quilbé et al. (2022) |
| *AeVPY* | Ae05g16930 | AeVPY-F<br>AeVPY-R | CTGAGGCTTCTCTTGCTTA<br>CTATGGCTGCTGCTATGT | Quilbé et al. (2022) |
| *AeUbi* | Ae10g10900 | AeUbi-F<br>AeUbi-R | TCAAAGTGAAGACTCTAACCG<br>CAAGTGAAGCACGGAACC | Quilbé et al. (2022) |
| *RiGADPH* | - | RiGADPH-F<br>RiGADPH-R | GACGTCTCAGTTGTTGATTTA<br>TTTGGCATCAAAAATACTAGA | Buendia et al. (2016) |
| *RiLSU* | - | RiLSU-F<br>RiLSU-R | GCATATCAATAAGCGGAGGA<br>ACTCCTCACGCTCCACAGA | Xue et al. (2015) |
