## Supplementary File 1 for "A Receptor Like Cytoplasmic Kinase evolved in *Aeschynomene* legumes to mediate Nod-independent rhizobial symbiosis"

**Supplementary File 1.** ***RLCK* sequences used for the phylogeny presented in Fig. 5a.**

**Papilionoideae**

- *Aeschynomene afraspera*

>AaRLCK_O

MPRILLQENSPASPSSMTLHRSSSLSYGALFFLLGGIVVLIILLILVFIYWMHIKQNADAKDKENEAPPTSHQQVEKYNESVEVVKMIVPTQQQRGSMEFISGNLRTISHFDFQTLRRATKNFHRKNLLGTGGFGPVYQGNLADGRLVAVKQLSLEKSQQGEREFLAEVRLITSIQHKNLVRLLGCCTDGSQRILVYEFMKNRSLDLIIYGKSDQFLNWSSRFQIILGVARGLQYLHEDSHLRIVHRDIKASNILLDEKFQPRIGDFGLARFFPEDQAYLSTQFAGTLGYTAPEYAIRGELSEKADIYSFGVLVLEIISCRKNTDLTLPSEMQYLPEYAWKLYEKSMVMDLVDPKLKEHGFVEKDVMQAFHVAFLCLQPHADLRPAMSEIVALLTFKIEMVSTPMRPAFLDRRRKKHDEENNSWEVISETFSTPKASDSNAPVSKGQ

>AaRLCK_P

MTRATLQESVIKNAVKPPSNDEPKYSPQRHKSDAFLFCLGALVVLIIVLVVILVFWKRLKGSAEPMRKKTPTKGKQGSMELFSGNLRTISYFDYQILRKATKNFSPANLLGSGGFGPVYRGKLVDGRLIALKKLSLNKSQQGEKEFLAEVRLITSIQHKNLVRLLGCCIDGPQRILVYEYMNNRSLDLFIYGNSDQFLNWSTRFQIILGVARGLQYLHEDSHLRIVHRDIKASNILLDEKFFPKIGDFGLARFFPEDQAYLSTQFAGTLGYTAPEYAIRGELSEKADIYSFGVLVLEIICCRKNTDLTLSSEMQYLPEYAWKLYESSKIMDIVDPKLKEKGFVEKDVMQAFHVAFMCLQPLPHLRPPMSEIVALLTFKIEMVTTPMRPAFLDRRRKKDYYNNDDDDVIDDYPETISQVFPSPMMSHSQ

- *Aeschynomene evenia*

> AeRLCK1 Ae1g26600

MALQSSSSWSSGALFFLLGGIVLVLILLLILVFIYWIRVKQHAETNEKEKEAPNSPQQVDKYNESIEVAKMIVPPQQQRGSMEFISGNLRTISYFDFQTIRRATKNFHRKNLLGAGGFGPVYQGKLADGRLVAVKQLSLEKSQQGEREFLAEVRLITSIQHKNLVRLLGCCTDGSQRILVYEFMKNRSLDLIIYGKSDQFLNWSTRFQIILGVARGLQYLHEDSHLRIVHRDIKASNILLDEKFQPRIGDFGLARFFPEDQAYLSTQFAGTLGYTAPEYALRGELSEKADIYSFGVLVLEIISCRKNTDLTLPSEMQYLPEYAWKLYERSMVMNLVDPKLREHGFVEKDVMQAFHVAFLCLQPHADLRPAMSEIVALLTFKIEMVSTPMRPAFLDRRRKNHDEENNSWEAISETFSTPKASDSNANVF

> AeRLCK2 Ae1g26600

MALRALFFVLGGIVVLITLLFLVFVYWIRIKQHSEVNKKEKEAPTSQQKVEEYNESLEVVKMVAPTQQQRGSMGFISNLRIISYFDFQTMRRATLNFHQKNLLGTGGFGPVYQGKLEDGRLVAVKQLSLEQSQQGDKEFLAEVRLITSIQHRNLVRLLGCCTDGSQRILVYEFMKNRSLDLIIYGKSDQFLNWSTRFQIILGVARGLQYLHEGSHLRIIHRDIKASNILLDEKFQPRIGDFGLARFFPEDQAYLSTQFAGTLGYTAPEYALRGELSEKADIYSFGVLVLEIISCRKNTDLTLPSEMQYLPEYAWKLYERSMVMNLVDPKLREHGFVEKDVMQAFHVAFLCLQPHADLRPAMSEIVALLTFKIEMVSTPMRPAFLDRRRKNHDEENNPWEAIS

- *Arachis duranensis*

>Aradu.CHL1Y

MPRNDIFYDSPGLVDPPPPASNEEPSPKKTDTTLFFLGGIVVFIILLIILVVFWKRIKGT

KNTTPPKTTKGKQGKLTETAEVMDMMFSSKQQQGSMELLSGNLRTISYFDYQILRRATKN

FSPTNLLGSGGFGPVYRQGEREFLAEVRLITSIQHKNLVRLLGCCMDGPQRILVYEYMKN

RSLDLFIYGNNADRFLNWSTRFQIVLGVARGLQYLHEDSHLRIVHRDIKASNILLDDKFH

PKIGDFGLARFFPEDQAYLSTQFAGTLGYTAPEYALRGELSEKADIYSFGVLLLEIICCR

KNTDLTLSLEMQYLPEYAWKLYENSKIMELVDPKLKEEGFVEKDVMQAFHVAFLCLQPLA

NLRPPMSEIVALLTFKIDMVTTPMRPAFLDRRRKTDFQTPSPEVISMLSHLP

>Aradu.UR751

MEVRLRSVVTYNNPESSMGAQNSSWSHHTSGALFFFLGGIIVLIILLILLFVYWMVIKRR

SPQEAEEKMEKKPSIMSQHHGSMDFFSGSLGTISHFNLNTLRRATRNFHSKNLLGSGGFG

PVYQGKLADGRLVAVKQLSAEKSQQGEKEFLAEVRLITSIQHKNLVRLLGCCTDGSQRIL

VYEYMKNKSLDRILYGKSDQFLDWSTRFQIILGVARGLQYLHEDSHLRIVHRDIKASNIL

LDEKFQPKIGDFGLARFFPEDQAYLSTQFAGTLGYTAPEYAIRGELSEKADIYSFGVLVL

EIISCRKNTDLTLPSEMQYLPEYAWKLYEKSMVMGLVDPKLREHGLVEKDVMRAFHVAFL

CLQPHAGLRPAMSEIVALLTFKVEMVSTPMRPSFFDRRPKKHDEDMPSWEAISDTFSVPI

ITDSNLPTSEDR

- *Arachis hypogaea*

>Arahy.8HXU4G

MPRNDIFYDSPGLVDPPPPASNGEPSPKKTDTTLFFLGGIVMFIILLIVLVVFWKRIKGT

KNTTPPKTTKGKQGKLTETAEVMDMMFSSKQQQGSMELLSGNLRTISYFDYQILRRATKN

FSPTNLLGSGGFGPVYRGKLMDGKLIALKKLSLNKSQQGEREFLAEVRLITSIQHKNLVR

LLGCCMDGPQRILVYEYMKNRSLDLFIYGNNADRFLNWSTRFQIVLGVARGLQYLHEDSH

LRIVHRDIKASNILLDDNFHSKIGDFGLARFFPEDQAYLSTQFAGTFYYKSQAWKLYENS

KIMELVDPKLKEEGFVEKDVMQAFHVAFLCLQPLANLRPPMSEIVALLTFKIDMVTTPMR

PAFLDRRRKTDFQTPSPEVISMLSHLP

>Arahy.HJ0ARX

MPRNDIFYDFPGLVDPPPPASNEEPSPKKTDTTLFFLGGIVMFIVLLIILVVFWKRIKGS

KNTTPPKTTKGKQGKLTETAEVMDMMFSSKQQQGSMELLSGNLRTISYFDYQILRRATKN

FSPTNLLGSGGFGPVYRGKLMDGRLIALKKLSLNKSQQGEREFLAEVRLITSIQHKNLVR

LLGCCIDGPQRILVYEYMKNRSLDFFIYGNNADRFLNWSTRFQIVLGVARGLQYLHEDSH

LRIVHRDIKASNILLDDKFHPKIGDFGLARFFPEDQAYLSTQFAGTLGYTAPEYALRGEL

SEKADIYSFGVLLLEIICCRKNTDLTLSLEMQYLPEYAWKLYENSKIMELVDPKLKEEGF

VEKDVMQAFHVAFLCLQPLANLRPPMSEIVALLTFKIDMVTTPMRPAFLDRRRKTDFQTP

SPEVISMLSHLP

>Arahy.MKUM1Y

MGAQNSSLFFFLGGIIVLIILLILLFVYWMIIKRRSPQEAEEKMEKEPSIMSQHQDEVVK

VTFSSKQQPGSMDFFSGSLGTISHFNLNTLRRATRNFHSKNLLGSGGFGPVYQGKLADGR

LVAVKQLSAEKSQQGEKEFLAEVRLITSIQHKNLVRLLGKSDQFLDWSTRFQIILGVARG

LQYLHEDSHLRIVHRDIKASNILLDEKFQPKIGDFGLARFFPEDQAYLSTQFAGTLKNTD

LTLPSEMQYLPEYAWKLYEKSMVMGLVDPKLREHGLVEKDVMRALHVAFLCLQPHAGLRP

AMSEIVALLTFKVEMVSTPMRPSFFDRRPKKHDEDMPSWEAISDTFSVPIISDSNPPTSE

DH

- *Arachis ipaensis*

>Araip.I6EYC

MGAQNSSLFFFLGGIIVLIILLILLFVYWMIIKRRSPQEAEEKMEKEPSIMSQHQDEVVK

VTFSSKQQPGSMDFFSGSLGTISHFNLNTLRRATRNFHSKNLLGSGGFGPVYQGKLADGR

LVAVKQLSAEKSQQGEKEFLAEVRLITSIQHKNLVRLLGCCTDGSQRILVYEYMKNTSLD

RILYGKSDQFLDWSTRFQIILGVARGLQYLHEDSHLRIVHRDIKASNILLDEKFQPKIGD

FGLARFFPEDQAYLSTQFAGTLGYTAPEYAIRGELSEKADIYSFGVLVLEIISCRKNTDL

TLPSEMQYLPEYAWKLYEKSMVMGLVDPKLREHGLVEKDVMRALHVAFLCLQPHAGLRPA

MSEIVALLTFKVEMVSTPMRPSFFDRRPKKHDEDMPSWEAISDTFSVPIISDSNPPTSED

H

>Araip.L2JGI

MITKLRLIQKLSLNKSQQGEREFLAEVRLITSIQHKNLVRLLGCCIDGPQRILVYEYMKN

RSLDFFIYGNNADRFLNWSTRFQIVLGVARGLQYLHEDSHLRIVHRDIKASNILLDDKFH

PKIGDFGLARFFPEDQAYLSTQFAGTLGYTAPEYALRGELSEKADIYSFGVLLLEIICCR

KNTDLTLSLEMQYLPEYAWKLYENSKIMELVDPKLKEEGFVEKDVMQAFHVAFLCLQPLA

NLRPPMSEIVALLTFKIDMVTTPMRPAFLDRRRKTDFQTPSPEVISMLSHLP

- *Cajanus cajan*

>C.cajan_44473

MFLFLGVIAVIIMLLILVFIFWRRVKRPAKEMENTAVASKQHEVMKMIVPNIQQSGPMGF

FSGNLRTISYFDFRTLRRATTNFHPRNLLGSGGFGPVYQGKLADGRLIAVKTLSLDKSQQ

QGEKEFLAEVRMITSIQHKNLVRLLGCCTDGSQRILVYEYMKNRSLDLIIHGKSDQFLNW

RTRFQIILGVARGLQYLHEDSHLRIVHRDIKASNILLDEKFQPRIGDFGLARFFPEDQAY

LSTQFAGTLGYTAPEYAIRGELSEKADIYSFGVLVLEIISCRKNTDLTLPSEMQYLPEYA

WKLYEKSMLMEIVDPKLREHGIVEKDVMQAFHVALLCLQPHADLRPAMSDIVAMLTFKVE

MISTPMRPAFLDRRRVMDDEHHSWEAVSEAFTTAVASDSASPPKAPIL

- *Cicer arietinum*

>Ca_20635 gnm1

MPRRVLQGETSNEHKHETIFFILGGIVVITILVILWIVFWKRAKRSSKLLEKKVTTEGEPVEAMKVVFSS

TQQQPSGSMELFSASLRSTSYFNYHILKKATNNFFHANLLGTGGFGPVYQGKLEDGRMIVVKTLSLNKSH

QGEKEFLAEVKLITSVQHKNLVRLLGCCIDGPQRILVYEYIKNKSLDLFIYGNSDQFINWRTRFQIILGV

ARGLQYLHEDSHLRIVHGDIKASNILLDDKFLPRIGDFGLARFFPGDQAYHLSTQFAGTLGYTAPEYAIR

GELSEKTDIYSFGVLLLEIISSKKNTDHTLPSDMQYLPEYAWKLYEKSSLLDLVDPKLREDGFVEKDVMQ

TTHVALLCLQPHAHLRPPMSEIVALLTFKIEMVTTPMRPAFLDRRRRKDEDNHSFEASLPSH

- *Glycine max*

>Glyma.09G062500

LRVLQATSPSNESHAPPHQSESLFYILGGIVVLAIVLIFLYVVRKRIKRPAQTMEFGGHNESAEVMKMIFSSNQHSGSKEFFSGNLRTISCFDYQTLKKATRNFHPDNLLGSGGFGPVYQGKLVDERLVAVKKLALNKSQQGEKEFLVEVRTITSIQHKNLVRLLGCCLDGPQRLLVYEYMKNRSLDLFIHGNSDQFLNWSTRFQIILGVARGLQYLHEDSHPRIVHRDIKASNILLDDKFHPRIGDFGLARFFPEDQAYLSTQFAGTLGYTAPEYAIRGELSEKADIYSFGVLVLEIICCRKNTEHTLPSEMQYLPEYAWKLYENARILDIVDPKLRQHGFVEKDVMQAIHVAFLCLQPHAHLRPPMSEIVALLTFKIEMVTTPMRPAFLDQRPREDGENHPLEALSQGFTSPIYVK

>Glyma.15G16900

LRVLQATSPSNEHAPQHKSGSSLFYILGGLVVLAIVLIFLYVVWKRIKRPAQTMEFGKHNESAEVMKMIFSSNQQSGSKEFFSGNLRTISCFDYQTLKKATENFHPDNLLGSGGFGPVYQGKLVDGRLVAVKKLALNKSQQGEKEFLVEVRTITSIQHKNLVRLLGCCVDGPQRLLVYEYMKNRSLDLFIHGNSDQFLNWSTRFQIILGVARGLQYLHEDSHQRIVHRDIKASNILLDDKFHPRIGDFGLARFFPEDQAYLSTQFAGTLGYTAPEYAIRGELSEKADIYSFGVLVLEIICCRKNTEHTLPSEMQYLPEYAWKLYENARILDIVDPKLREHGFVEKDVMQANHVAFLCLQPHAHLRPPMSEIVALLTFKIEMVTTPMRPAFLDRRPRKGDENHPLEALSQGFTSPIYL

>Glyma.17G055900

PSHHDPGVFFFLGGIVVLVILLILVFIFWRRIKRPAKVMQHGKYIEPSEVMKMIVPNIKQPGPMEFISGNLRTISYFDFRTLRRATKNFHPRNLLGSGGFGPVYQGKLADGRLIAVKTLSLDKSQQGEKEFLAEVRMITSIQHKNLVRLIGCCTDGPQRILVYEYMKNRSLDLIIYGKSDQFLNWSTRFQIILGVARGLQYLHEDSHLRIVHRDIKASNILLDEKFQPRIGDFGLARFFPEDQAYLSTQFAGTLGYTAPEYAIRGELSEKADIYSFGVLVLEIISCRKNTDLTLASEKQYLPEYAWKLYEKSMLMEIVDPKLQEQGIEEKDVMQAFHVALLCLQPHADLRPAMSEIVAMLTFKVEMVTAPMRPIFVDRRRVMDDEHHSWETIYEAFTTAVAS

- *Lotus japonicus*

>LjAMK8 Lj4g3v2140260

MSILLTKTFEELELQSQPECLLLSPKMHRRILQGKSPSSSSSPHNSGALFFFLGGTVVLIILLILLFVFWRRIKRPAERTTPTNEQHGRITMAASTTKAQQTGFMEFVSGNLRTISYFDFQTLRKATKNFHRTNLLGSGGFGPVYQGKLADGRLIAVKQLSLDKSQQGDKEFLAEVRMITSIQHKNLVRLMGCCTDGPQRILVYEYMKNRSLDPFVYGNSDQFLNWSTRFQIILGVARGLQYLHEDSHIRIVHRDIKASNILLDEKFRPRIGDFGLARFFPEDQAYLSTQFAGTLGYTAPEYAIRGELSEKADIYSFGVLVLEIICCRKNTDLTLPSQMQYLPEYGWKLYEKSMVMDLVDPKLREDGFVEKDVMQAFHVAFLCLQPLPDMRPAMSEIVALLTFKIDMVTTPMRPAFLDRRRKMDEEHHSWEAISKSFKSPGASDYSP

>LjAMK24 Lj6g3v1270820

MPSRVLQGEAPLPSKESQIPSSRYKHETLFFVLGGIVVVAILFILWFVFRKRIKQPTKPKGKTAPSKEHKEVMKMVFPSKQQSGSKSMSMEFFSGNLQSICFFDYQTLRKATHNFFPGNLLGSGGYGPVYRGKLVDGRMIAVKTLSHNKSQQGEREFLAEVKMITSIQHKNLVRLLGCCIDGPQRILVYEYMKNRSLELFIYGNGDQFLNWRTRFQIILGVARGLQYPHEDSHLRIVHRDIKASNILLDDKFQPRIGDFGLARFFPEDQDYLSTQFAGT

- *Medicago truncatula*

>MtrunA17_Chr4g0071371

MSIPTKTAQLQLQLQQEYILLTMPRRLLQLQETPPSSPHHVSGALYFFLGVIVVLVILLIILFVFWKRFRRGSGKTEELPEETAPPPPSQTQKQEEVMKRIIPTNQQSGFMEFISGNLRTISYFDFQTLRKATKNFHRRYLLGSGGFGPVYQGKLADGRLVACKKLSLDKSHQGEREFLAEVRMITSIQHKNLVRLLGCCSDGPQRILVYEYMKNRSLDFFIHGKSDEFLNWSTRFQIILGVARGLQYLHEDSHVRIVHRDIKASNILLDEKFQPRIGDFGLARFFPEDQAYLSTQFAGTLGYTAPEYAIRGELSEKADIYSFGVLLLEIISCRKNTDLTLPSDMQYLPEYAWKLYEKSMVMELIDPKLIEKGYVEKDVMQAFHVAFLCLQPHPDLRPAMSQIVALLTFKIDMVTTPMRPAFLDRRRVMDDENHSWEVISEVLQTPAASDSTL

>MtrunA17_Chr2g0299041

MPLRVLQGEASLPSNDESKKSSEHKRETMFFVLGGIVVVTIILILGWIVFRKRVKRSPNPVGKTVPNEAEPTEVMKAIFPSKQQSSGSMEFFSGSLRSISYFDYQTLRKATNNFFHGNLLGSGGFGPVYKGKLEDGRIIAVKALSLNKSQQGEREFLAEVKLITSIQHKNLVRLLGSCIDGPQRILIYEYMKNRSLDLFIYGNNDRFLNWSTRYQIILGVARGLQYLHEDSHLRIVHRDIKASNILLDDKFLPRIGDFGLARFFPEDQAYLSTQFAGTLGYTAPEYAIRGELSEKADIYSFGVLLLEIICCRKNTDHTLPPDMQYLPEYAWKLYEKSSLLDLVDPKLKQDGFVEKDVMQATHVALLCLQPHAHLRPRMSEIVALLTFKIEMVTTPMRPAFLGLRSRKDEDNHSFEVTSMANH

- *Phaseolus vulgaris*

>Phvul.003G137700g

MTLLTNTFESECLLLFPRMPRRILQVTAEEPPSSSVALKTSASRYAPGVFFFLGGTVLLI

ILLILLFIFWRRTKGPAKVTENTTLTGKYIEPSEVMKMIVPNIQQGQMEFISGNLRTISY

FDFRTLRRATKNFHPRNLLGSGGFGPVYQGKLADGRLVAVKTLSLDKSQQGEKEFLAEVR

MITSIQHKNLVRLIGCCTDGPQKILVYEYMKNRSLDLIIYGKSDQFLNWNTRFQIILGVA

RGLQYLHEDSQLRIVHRDIKASNILLDEKFQPRIGDFGLARFFPEDQAYLSTQFAGTLGY

TAPEYAIRGELSEKADIYSFGVLVLEIISCRKNTDLTLPSEMQYLPEYAWKLYEKSMLME

IVDPKLREHGMEEKDVMQAFQVALSCLQPHADLRPAMSEIVALLTFKVEMVTAPMRPTFF

HRRRVMDDENHSWGAISSEGSTTAVTSS

> Phvul.009G233300g

MTLRVLQEASPSNRTTQHKSEALFYILGGIIVLAIVLIFLYVFRKRIKRQPQNMTVTTTEKQGIMKHNESADMMKMIFSSNPQSGSMEFFSGNLRTINCFDYQTLKNATTNFHADNFLGSGGFGPVYKGKLVDGRLIAVKKLSLNKSQQGEKEFLVEVRTITSIQHKNLVRLLGCCIDGPQRIIVYEYMKNRSLDLFIHENSDQFLKWGTRFQIILGVARGLQYLHEDSHQRIVHRDIKASNILLDDKLQPRIGDFGLARFFPEDQAYLSTQFAGTLGYTAPEYAIRGELSEKVDIYSFGVLLLEIICCRKNTDQTLPSEMQYLPEYAWKLYENARILDIVDPKLRQDGFVEKDVMQAIHVAFLCLQPDPQLRPPMSEIVALLTFKIEMVTTPMRPAFLDRRPGKDDENHCLRGLSEGLTSPI

- *Vigna angularis*

>Vang11g13150

MKSFDTVTAQEPPSSSVALKASVSRYDPGIFFFLGGTVLLIILLILLFIFWRRTKGPAKV

NTTLTCQQHGQMEFISGNLRTISYFDFRTLSRATKNFHPRNLLGSGGFGPVYQGKLADGR

LVAVKTLSLDKSQQGEKEFLAEVRMITSIQHKNLVRLLGCCTDGPQKILVYEYMKNRSLD

LIIYGGSDQFLNWNTRFQIILGVARGLQYLHEDSHLRIVHRDIKASNILLDEKFQPRIGD

FGLARFFPEDQAYLSTQFAGTLGYTAPEYAIRGELSEKADIYSFGVLVLEIICCRKNTDL

TLPSEMQYLPEYAWKVYEKSMLMEIVDPRLREHGMEEKDVMQAFHVALSCLQPHADLRPA

MSEIVALLTFKVEMVAKPIRPTFVHRRRVMDDENHSWGAISSSEPSTTAVASL

> Vang 09g06880

MSPRTLQEASLSDMTQTPQHKSEGTLFYILGGIVVLLIVLIFLYVLRKRLKWSPHNMTEQQEIRKHIESADMMKMIFSSNQQSGSKEFFSGNLRTINCFDYQTLKNATMNFHADNFLGSGGFGPVYKGKLADGRVVAVKKLSLNKSQQGEKEFLVEVRTITSIQHKNLVRLLGCCIDGPQRIIVYEYMKNRSLDLFIHENSDQFLNWRTRFQIILGVARGLQYLHEDSHQKIVHRDIKASNILLDEKFQPRIGDFGLARFFPEDQAYLSTQFAGTLGYTAPEYAIRGELSEKADIYSFGVLLLEIICCRKNTDHTLPSEMQYLPEYAWKLYENARILDIVDPKLQEHGLVEKDVMQAIHVAFLCLQPDAHLRPPMSEIVALLTFKIEMVTTPMRPAFLDRAAKKDDEKQSLGTIYHGLTSPI

- *Vigna radiata*

>Vradi07g22830

MKSFDTVTAQEPPSSSVASKAPTSRYDPGMFFFLGGTVLLIILLILLFIFWRRTKGPAKV

NTTLTCQQNGQMEFISGNLRTISYFDFRTLRRATKNFHPRNLLGSGGFGPVYQGKLADGR

LVAVKTLSLDKSQQGEKEFLAEVRMITSIQHKNLVRLLGCCTDGPQKILVYEYMKNRSLD

LIIYGGSDQFLNWNTRFQIILGVARGLQYLHEDSHLRIVHRDIKASNILLDEKFQPRIGD

FGLARFFPEDQAYLSTQFAGTLGYTAPEYAIRGELSEKADIYSFGVLVLEIICCRKNTDL

TLPSEMQYLPEYAWKLYEKSMLMEIVDPRLREDGMEEKDVMQAFHVALSCLQPHADLRPA

MSEIVALLTFKVEMVAKPVRPTFVHRRRVMDDENHSWGAISSSEPSTTALASL

>Vradi05g02500

MSPRTLQEIGKHTESADIMQMIFSSNQQSGSKELCSGNFRTINCFDYQTLKNATMNFHAD

NFLGSGGFGPVYKGKLADGRVVAVKKLSLNKSQQGEKEFLVEVRTITSIQHKNLVRLLGC

CVDGPQRILVYEYMKNRSLDLFIHENSDQFLNWRTRFQIILGVARGLQYLHEDSQQRIVH

RDIKASNILLDDKFQPRIGDFGLARFFPEDQAYLSTQFAGTLGYTAPEYAIRGELSEKAD

IYSFGVLLLEIICCRKNTDHTLPSDMQYLPEYAWKLYENAKILEIVDPKLQEHGLVEKDV

MQAIHVAFLCLQPDAHLRPPMSEIVALLTFKIEMVTTPMRPAFLYRRAKKDDEKQHLGAI

YQDLTSPI

- *Vigna unguiculata*

>Vigun03g3908001

MTLFTNTVESECFLLPQIMSRRVLQVTAQEPPSSSAALKTSASRYDPGVFFFLGGTVLLIIVLILLFIFW

RRTKGPAKVTENTTLTCQQHGKYIEPSEVMKMMAPNIQPGQMEFISGNLRTISHFDFRTLKRATKNFHPR

NLLGSGGFGPVYQGKLADGRLVAVKTLSLDKSQQGEKEFLAEVRMITSIQHKNLVRLLGCCTDGPQKILV

YEYMKNRSLDLIIYGESDQFLNWNTRFQIILGVARGLQYLHEDSHLRIVHRDIKASNILLDEKFQPRIGD

FGLARFFPEDQAYLSTQFAGTLGYTAPEYAIRGELSEKADIYSFGVLVLEIISCRKNTDLTLPSEMQYLP

EYAWKLYEKSMLMEIVDPRLRQHGMEEKDVMQAFHVALSCLQPHADLRPAMSEIVALLTFKVEMVTKPIR

PTFVHRRRVMDDENHSWGAISSGTSTTAVASSSS

>Vigun09g032500

MSPRILQEASPSTKPEHKSGGTLFYFLGGIVVLAIVLIFFYVLVLRKRIKRSPQGMTEQKEIRKHIESAD

MMKMIFSSNQQSGSKEFFSGNLRTINCFDYQTLKNATMNFHADNFLGSGGFGPVYKGKLADGRVIAVKKL

SLNKSQQGEKEFLVEVRTITSIQHKNLVRLLGYCIDGPQRILVYEYMKNRSLDLFIHENSDQFLNWRTRF

QIILGVARGLQYLHEDSHQRIVHRDIKASNILLDDKFQPKIGDFGLARFFPEDQAYLSTQFAGTLGYTAP

EYAIRGELSEKADIYSFGVLLLEIICCRKNTDHTLASEMQYLPEYAWKLYENERILDIVDPKLREHGLVE

KDVMQAIHVAFLCLQPDAHLRPPMSEIVALLTFKIEMVTTPMRPAFLDRGARNDDDDEKEHKGLTSPI

**Caesalpinoideae**

- *Mimosa pudica*

> Mimpu109S05362

MELMSGSLRTIIYFDYSTLKMATKNFHSGNLLGSGGFGPVYRGKLANGRLIAVKKLSINKSKQGEKEFLAEVKMITSIQHKNLVRLIGCCTEGIERILVYEYLKNKSLDLYIYGNSVQFLDWRTRFQIILGIARGLQYLHEDSHIRIVHRDIKASNILLDEKFHPRIGDFGLARFFPEDEDYLSTQFAGTLGYTAPEYAIRGELSEKADIYSFGVLVLEIVCGRKNTDLTLPPEMQYLPDYAWKLFDKSRLMDLVDPKLQELGFTEKDVMQAIHVAFLCIQSQANLRPPMSEIVALLTFKIGMVRTPMRSSFIGRRQRKDDENHSWEALSESSSPFPSDFSSTPK

- *Chamaecrista fasciculata*

>Chafa1000S13389

MARLVLEVQPPTSQGNHGSSSLQISEGLFFLLGGVVMLIILVILLYVFRRHIIPENLMKMRAPTQKHLGKQIVESAEVVEMIVPIKQHLGSMELLSGNLRTIIYFDFQTLKKATMNFHSRNLLGSGGFGPVYRGKLADGRLIAVKKLSLNKSHQGEKEFLAEVRMITSIQHKNLVGLIGCCTDGLQRILVYEYMKNRSLDLFIHGTRDQFLNWSTRFQIILGIARGLQYLHEDSQLRIVHRDIKASNILLDEKFQPRIGDFGLARFFPEDQDYLSTQFAGTLGYTAPEYAIRGELSEKADIYSFGVLVLEIISNRKNTDLTLPSEMQYLPEYAWKLREKSRWMDLVDPKLQEHGFVEKDVMRTIHVALLCLQTQANLRPPMSEVVALLTFKIDTVKTPMRPAFLDRRRNKDDENLSWEAASGPMSGDSASIAKPPN

**Non-legume species**

- *Prunus persica*

>Prupe1G068800

MEMNSTSKPTSPALFFFLGGIVMLIILLVLIFVFRKLIKPEELKKLVARARRQPESKDLFSGNLRTISYFDFRTLKMATKNFHPGNLLGVGGFGPVYRGKLGDGRLIAAKKLCLDKSQQGESEFLTEVKLITSVQHRNLVRLIGCCSDGPQRLLVYEYMKNRSLDLIVYGKSDRFLNWSTRFQIIVGIARGLQYLHEDSPLRIIHRDIKASNILLDEKYQPKIGDFGLARFFPEDQAYLSTTFAGTLGYTAPEYAIRGELSEKADIYSFGVLVLEIISGRKNTDLTLPSEMQYLPEYAWKLFETSNVIELVDPKLHENGFVERDVLQAIQVAFLCLQPHANLRPPMSEVVAMLTCKVEMIGTPMKPAFLARRRTKDQNLSWDTISEVFPSPFQSESTSLPKPPT

- *Oryza sativa*

> OsRLCK171 Os04g56360

MANPNASAVFLAFIVILIIVIFILLGICWKFLRPDIMRRLMRPKRAPSEVPEYFSGNMSGNLRTITYFDYATLKKATRDFHQKNQLGRGGFGPVYLGKLDDGRKVAVKQLSVGKSGQGESEFFVEVNMITSIQHKNLVRLVGCCSEGQQRLLVYEYMKNKSLDKILFGVDGAPFLNWKTRHQIIIGIARGLQYLHEESNLRIVHRDIKASNILLDDKFQPKISDFGLARFFPEDQTYLSTAFAGTLGYTAPEYAIRGELTVKADTYSFGVLVLEIVSSRKNTDLSLPNEMQYLPEHAWRLYEQSKILELVDAKLQADGFDEKEVMQVCQIALLCVQPFPNLRPAMSEVVLMLTMKTTEQSVIPAPVRPAFLDRKSLKDKNNGGGSDTAAEMRSTAYWLGTPSPMVDRPYDMSCGI
