## Supplementary File 2 for "A Receptor Like Cytoplasmic Kinase evolved in *Aeschynomene* legumes to mediate Nod-independent rhizobial symbiosis"

**Supplementary File 2. *RLCK* sequences used for the analysis presented in Fig. 5b.**

>A.afraspera_RLCK_O (RNASeq)

MPRILLQENSPASPSSMTLHRSSSLSYGALFFLLGGIVVLIILLILVFIYWMHIKQNADAKDKENEAPPTSHQQVEKYNESVEVVKMIVPTQQQRGSMEFISGNLRTISHFDFQTLRRATKNFHRKNLLGTGGFGPVYQGNLADGRLVAVKQLSLEKSQQGEREFLAEVRLITSIQHKNLVRLLGCCTDGSQRILVYEFMKNRSLDLIIYGKSDQFLNWSSRFQIILGVARGLQYLHEDSHLRIVHRDIKASNILLDEKFQPRIGDFGLARFFPEDQAYLSTQFAGTLGYTAPEYAIRGELSEKADIYSFGVLVLEIISCRKNTDLTLPSEMQYLPEYAWKLYEKSMVMDLVDPKLKEHGFVEKDVMQAFHVAFLCLQPHADLRPAMSEIVALLTFKIEMVSTPMRPAFLDRRRKKHDEENNSWEVISETFSTPKASDSNAPVSKGQ

>A.americana_RLCK_O (PCR)

GKLADGRMVAVKQLSVEKSQQGEREFLAEVRLITSIQHKNLVRLLGCCTDGSQRILVYEFMKNRSLDLIIYGKSDQFLNWSTRFQIILGVARGLQYLHEDSHLRIVHRDIKASNILLDEKFHPKIGDFGL

>A.ciliata_RLCK1 (PCR)

NERKKEAPNSPQQVEKYNESIEVVKMIVPTQQQRGSMEFISGNLRTISYFDFQTIRRATKNFHRKNLLGAGGFGPVYQGKLADGRLVAVKQLSLEKSQQGEREFLAEVRLITSIQHKNLVRLLGCCTDGSQRILVYEFMKNRSLDLIIYGKSDQFLNWSTRFQIILGVARGLQYLHEDSHLRIVHRDIKASNILLDEKFQPRIGDFGLARFFPEDQAYLSTQFAGTLGYTAPEYALRGELSEKADIYSFGVLVLEIISCRENTDLTLPSEMQYLPEYAWKLYERSMVMNLVDPKLREHGFVEKDVMQAFHVAFLCLQPHAD

>A.ciliata_RLCK2 (RNAseq)

MALRALFFVLGGIVVLITLLFLVFIYWICIKQRSEANKKEKGAPTSQQKVEEYNETIEVVKMVAPTQQQRGSMGFISGNLRTISYFDFQTMRRATLNFHHKNLLGTGGFGPVYQGKLEDGRLVAVKQLSLEQSQQGDKEFLAEVRLITSIQHRNLVRLLGCCTDGSQRILVYEFMKNRSLDLIIYGKSDQFLNWSTRFQIILGVARGLQYLHEGSHLRIVHRDIKASNILLDEKFQPRIGDFGLARFFPEDQAYLSTQFAGTLGYTAPEYALRGELSEKADIYSFGVLLLEIISCRKNTDLTLPSEMQYLPEYAWKLYERSMVMNLVDQKLREHGFVEKDVMQAFHVAFLCLQPHADLRPAMSEIVALLTFKIEMVSTPMRPAFLDRRRKKHDEENNSWEAIS

>A.deamii_RLCK1 (PCR)

MVAFASYSISDICAWYFQVEKYNKSIEEVKMIVPTQQQRGSMEFISGNLRTISYFDFQTIRRATKNFHQKNLLGSGGFGPVYQAWKLYERSMVMDLVDPKLREHGFVEKDVMQAFHVAFLCLQPHADLRPAMSEIVALLTFKIEM

>A.deamii_RLCK2 (RNAseq)

MALGALFFVLGGVVVLIILLILVFIYWIRINHHAEANKKEKEAPTNQQQVEKHNESIEVVKMVVPAQQQRGSMEFISGNLRTISYFDFQTMRRATLNFHPKNLLGTGGFGPVYQGKLADGRLVAVKQLSLEQSQQGDKEFLAEVRLITSIQHRNLVRLLGCCTDGSQRILVYEFMKNRSLDLVIYGKSDQFLNWSTRFQIILGVARGLQYLHEGSHLRIIHRDIKASNILLDEKFQPRIGDFGLARFFPDDQAYLSTQFAGTLGYTAPEYALRGELSEKADIYSFGVLVLEIISCRKNTDLTLPSEMQYLPEYAWKLYERSMVMDLVDPKLREHGFVEKNVMQAFHVAFLCLQPHADLRPAMSEVVALLTFKIEMVSTPMRPAFLDRRH

>A.denticulata_RLCK1 (PCR)

YWMRIKQHAHTNEKEKEAPNSPQQVEKYNESIEVVKMIVPPQQQRGSMEFISGNLRTISYFDFQTIRRATKNFHRKNLLGAGGFGPVYQGKLADGRLVAVKQLSLEKSQQGEREFLAEVLITSIQHKNLVRLLGCCTDGSQRILVYEFMKNRSLDLIIYGKSDQFLDWSTRFQIILGVARGLQYLHEDSHLRIVHRDIKASNILLDEKFQPRIGDFGLARFFPEDQAYLSTQFAGTLGYTAPEYALRGELSEKADIYSFGVLVLEIISCRENTDLTLPSEMQYLPEYAWKLYERSMVMDLVDPKLREHGFVEKDVMQAFHVAFLCLQPHADLRPAMSEIVAL

>A.denticulata_RLCK2 (RNAseq)

MALRALFFVLGGIVVLITLLFLVFIYWIRIKQHSEANKKEKGAPTSQQKVEEYNESLEVVKMVAPTQQQRGSMGFISGNLRIISYFDFQTMRRATLNFHHKNLLGTGGFGPVYQGKLEDGRLVAVKQLSLEQSQQGDKEFLAEVRLITSIQHRNLVRLLGCCTDGSQRILVYEFMKNRSLDLIIYGKSDQFLNWSTRFQIILGVARGLQYLHEGSHLRIIHRDIKASNILLDEKFQPRIGDFGLARFFPEDQAYLSTQFAGTLGYTAPEYALRGELSEKADIYSFGVLVLEIISCRKNTDLTLPSEMQYLPEYAWKLYERSMVMNLVDPKLREHGFVEKDVMQAFHVAFLCLQPHADLRPAMSEIVALLTFKIEMVSTPMRPAFLDRRRKNHDEENNSWEAIS

>A.evenia_RLCK1 (RNAseq)

MALQSSSSWSSGALFFLLGGIVLVLILLLILVFIYWIRVKQHAETNEKEKEAPNSPQQVDKYNESIEVAKMIVPPQQQRGSMEFISGNLRTISYFDFQTIRRATKNFHRKNLLGAGGFGPVYQGKLADGRLVAVKQLSLEKSQQGEREFLAEVRLITSIQHKNLVRLLGCCTDGSQRILVYEFMKNRSLDLIIYGKSDQFLNWSTRFQIILGVARGLQYLHEDSHLRIVHRDIKASNILLDEKFQPRIGDFGLARFFPEDQAYLSTQFAGTLGYTAPEYALRGELSEKADIYSFGVLVLEIISCRKNTDLTLPSEMQYLPEYAWKLYERSMVMNLVDPKLREHGFVEKDVMQAFHVAFLCLQPHADLRPAMSEIVALLTFKIEMVSTPMRPAFLDRRRKNHDEENNSWEAISETFSTPKASDSNANVF

>A.evenia_RLCK2 (RNAseq)

MALRALFFVLGGIVVLITLLFLVFVYWIRIKQHSEVNKKEKEAPTSQQKVEEYNESLEVVKMVAPTQQQRGSMGFISNLRIISYFDFQTMRRATLNFHQKNLLGTGGFGPVYQGKLEDGRLVAVKQLSLEQSQQGDKEFLAEVRLITSIQHRNLVRLLGCCTDGSQRILVYEFMKNRSLDLIIYGKSDQFLNWSTRFQIILGVARGLQYLHEGSHLRIIHRDIKASNILLDEKFQPRIGDFGLARFFPEDQAYLSTQFAGTLGYTAPEYALRGELSEKADIYSFGVLVLEIISCRKNTDLTLPSEMQYLPEYAWKLYERSMVMNLVDPKLREHGFVEKDVMQAFHVAFLCLQPHADLRPAMSEIVALLTFKIEMVSTPMRPAFLDRRRKNHDEENNPWEAIS

>A.filosa-RLCK1 (PCR)

EKYNESIEVVKMIVPTQQQRGSMEFISGNLRTISYFDFQTMRRATKNFHRKNLLGTGGFGPVYQGKLADGRLVAVKQLCLEKSQQGEKEFLAEVRLITSIQHKNLVALLGCCTDGSQRILVYEFMKNRSLDLIIYGNTDRFLNWSTRFQIILGVARGLQYLHEDSHLRIVHRDIKASNILLDEKFQPRIGDFGLARFFPEDQAFLSTQFAGT

>A.filosa_RLCK2 (RNASeq)

MALRALFFVLGGIVVLFILLILVFIYWIRIKQHAEEVAPTSHQQVEKYNNESIEVVKMVVPTQQQRGSMEFISGNLRTISYFDFQTMRRATLNFHHKNLLGTGGFGPVYQGKLADGRLVAVKQLSLEESQQGDKEFLAEVRLITSIQHRNLVRLLGCCTDGSQRILVYEFMKNRSLDLIIYGKSDQFLNWSTRFQIILGVARGLQYLHEGSHLRIVHRDIKASNILLDEKFQPRIGDFGLARFFPEDQAYLSTQFAGTLGYTAPEYALRGELSEKADIYSFGVLVLEIISCRKNTDLTLPSEMQYLPEFAWKLYERSMVMDLVDPKLREHGFVEKDVMQAFHVAFLCLQSHADLRPAMSEIVALLTFKIEMVSTPMRPAFLDRRRKKHDEENYSWEAIS

>A.fluminensis_RLCK_O (PCR)

LLIYWMRIKLHAEANEKEKEKEKEKEAPISQQQVEKYNESIEVVKMVVPTQQQPGSTEFISGNLRTISYFDFQTIRRVTKNFHRKNLLGTGGFGPVYQGKLADGRMVAVKQLSLEKSQQGEREFLAEVRLITSIQHKNLVRLLGCCTDGSQRILVYEFMKNRSLDLIIYGKSDQFLNWSTRFQIILGVARGLQYLHEDSHLRIVHRDIKASNILLDEKFHPRIGDFGLARFFPEDQAYLSTQFAGTLGYTAPEYAIRGELSEKADIYSFGVLVLEIISCRKNTDLTLPSEMQYLPEYAWKLYEKSMVMDLVDPKLREHGFVEKDVMQAFHVAILCLQPHADLRPAMSEIVALLTFK

>A.montevidensis_RLCK_O (PCR)

LVFIYWMRIKQYAEANENEKEAPRSHRQLEKYNESIEVVKMIVPTQQQRGSMEFISGSLRTISYFDFQTMRRATKNFHRKNLLGTGGFGPVYQGKLADGRLVAVKQLSLEKSQQGEREFLAEVRLITSIQHKNLVRLLGCCTDGSQRILVYEFMKNRSLDLIIYGKSDQFLNWSTRFQIILGVARGLQYLHEDSHLRIVHRDIKASNILLDEKFHPRIGDFGLARFFPEDQAYLSTQFAGTLGYTAPEYAIRGELSEKADIYSFGVLVLEIISCRKNTDLTLPSEMQYLPEYAWKLYEKSMVMDLVDPKLKEHGFVEKDVMQAFHVAFLCLQPHADLRPAMSEIVALL

>A.patula_RLCK_O (PCR)

LILVFIYWMRIKQHPDANVKEKEAPTSHQQVEKYNESIEVVKMAVPTQQQRGSLEFISANLRTVSHFDFQTMRRATKNFHRKNLLGTGGFGPVYQGKLADGRLVAVKQLSLEKSQQGEREFLAEVRLITSIQHKNLVRLLGCCTDGSQRILVYEFMKNRSLDLILYGKSDQFLNWSTRFQIILGVARGLQYLHEDSHLRIVHRDIKASNILLDENFHPRIGDFGLARFFPEDQAYLSTQFAGTLGYTAPEYAIRGELSEKADIYSFGVLVLEIISCRKNTDLTLPSEMQYLPEYAWKLYEKSMVMDLMDPKLKEHGFVEKDVLQAFQVAFLCLQPHADLRPAMSEIVALLTFK

>A.rudis_RLCK1 (PCR)

EKEAPNSPQQVEKYNESIEVVKMIVPTQQQRGSMEFISGNLRTISYFDFQTIRRATKNFHRKNLLGAGGFGPVYQGKLADGRLVAVKQLSLEKSQQGEREFLAEVRLITSIQHKNLVRLLGCCTDGSQRIVYEFMKNRSLDLIIYGKSDQFLNWSTRFQIILGVARGLQYLHEDSHLRIVHRDIKASNILLDEKFQPRIGDFGLARFFPEDQAYLSTQFAGTLGYTAPEYALRGELSEKADIYSFGVLVLEIISCRENTDLTLPSEMQYLPEYAWKLYERSMVMNLVDPKLREHGFVEKDVMQAFHVAFLCLQPHADLRPA

>A.rudis_RLCK2 (RNAseq)

MALRALFFVLGGIVVLITLLFLVFIYWIRIKQRSEANKKEKGAPTSQQKVEEYNETIEVVKMVAPTQQQRGSMGFISGNLRTISYFDFQTMRRATLNFHHKNLLGTGGFGPVYQGKLEDGRLVAVKQLSLEQSQQGDKEFLAEVRLITSIQHRNLVRLLGCCTDGSQRILVYEFMKNRSLDLIIYGKSDQFLNWSTRFQIILGVARGLQYLHEGSHLRIVHRDIKASNILLDEKFQPRIGDFGLARFFPEDQAYLSTQFAGTLGYTAPEYALRGELSEKADIYSFGVLLLEIISCRKNTDLTLPSEMQYLPEYAWKLYERSMVMNLVDPKLREHGFVEKDVMQAFHVAFLCLQPHADLRPAMSEIVALLTFKIEMVSTPMRPAFLDRRRKNHDEENNSWEAIS

>A.scabra_RLCK1 (RNAseq)

MPRILFQFPRVDLVTDYCRSENPAASSSMALQSSSSWSSGALFFLLGGIVVLILLLILVFIYWMRIKQHADTNEKEAPNSPQQVEKYNESIEVVKMIVPPQQQRGSMEFISGNLRTISYFDFQTIRRATKNFHRKNLLGAGGFGPVYQGKLADGRLVAVKQLSLEKSQQGEREFLAEVRLITSIQHKNLVRLLGCCTDGSQRILVYEFMKNRSLDLIIYGKSDQFLNWSTRFQIILGVARGLQYLHEDSHLRIVHRDIKASNILLDEKFQPRIGDFGLARFFPEDQAYLSTQFAGTLGYTAPEYALRGELSEKADIYSFGVLVLEIISCRENTDLTLPSEMQYLPEYAWKLYERSMVMDLVDPKLREHGFVEKDVMQAFHVAFLCLQPHADLRPAMSEIVALLTFKIEMVSTPMRPAFLDRRRKNHDVENNSWEAISETFSTPKASDSNAPVPKGQ

>A.scabra_RLCK2 (RNAseq)

MALRALFFVLGGIVVLITLLFLVFIYWIRIKQHSEANKKEKGAPTSQQKVEEYNESLEVVKMVAPTQQQRGSMGFISGNLRIISYFDFQTMRRATLNFHHKNLLGTGGFGPVYQGKLEDGRLVAVKQLSLEQSQQGDKEFLAEVRLITSIQHRNLVRLLGCCTDGSQRILVYEFMKNRSLDLIIYGKSDQFLNWSTRFQIILGVARGLQYLHEGSHLRIIHRDIKASNILLDEKFQPRIGDFGLARFFPEDQAYLSTQFAGTLGYTAPEYALRGELSEKADIYSFGVLVLEIISCRKNTDLTLPSEMQYLPEYAWKLYERSMVMNLVDPKLREHGFVEKDVMQAFHVAFLCLQPHADLRPAMSEIVALLTFKIEMVSTPMRPAFLDRRRKNHDEENNSWEAIS

>A.selloi_RLCK1 (PCR)

ATNNFHRKNLLGAGGFGPVYQGKLADGRLVAVKQLSLEKSQQGEREFLAEVRLITSIRHKNLVRLLGCCTDGSQRILVYEFMKNRSLDLIIYGKSDQFLNWSTRFQIILGVARGLQYLHEDSHLRIVHRDIKASNILLDEKFQPRIGDFGLARFFPEDQAYLSTQFAGTLGYTAPEYALRGELSEKADIYSFGVLVLEIISCRENTDLTLPSEMQYLPEY

>Aselloi_RLCK2 (RNAseq)

MALRALFFVIGGIVVLITLLFLVFIYWIRIKQHSEANKKEKGAPTSQQKVEEYNESIEGVKMVAPTKQQRGSTGFISGNLRTISYFDFQTMRRATLNFHHKNLLGTGGFGPVYQGRLEDGRLVAVKQLSLEQSQQGDKEFLAEVRLITSIQHRNLVRLLGCCTDGSQRILVYEFMKNRSLDLIIYGKSDQFLNWSTRFQIILGVARGLQYLHEGSHLRIVHRDIKASNILLDEKFHPRIGDFGLARFFPDDQAYLSTQFAGTLGYTAPEYALRGELSEKADIYSFGVLVLEIISCRKNTDLTLPSEMQYLPEYAWKLYERSMVMNLVDPKLREHGFVEKDVMQAFHVAFLCLQPHANLRPAMSEIVALLTFKIEMVSTPMRPAFLDRRRNNHDGENNSWEAIS

>Asensitiva_RLCK1 (PCR)

NIKRATNNFHRKNLLGAGGFGPVYQGKLADGRLVAVKQLSLEKSQQGEREFLAEVRLITSIQHKNLVRLLGCCTDGSQRILVYEFMKNRSLDLIIYGKNDQFLNWSTRFQIILGVARGLQYLHEDSHLRIVHRDIKASNILLDEKFQPRIGDFGLARFFPEDQAYLSTQFAGT

>A.sensitiva_RLCK2 (RNAseq)

MALRALFFVLGGIVVLFTLLFLVFIYWIRIKQNSEANKKEKGAPTSQQKVEEYNESIEGVKMVAPTKQQRGSMGFISGNLRTISYFDFQTMRRATLNFHHKNLLGTGGFGPVYQGKLEDGRLVAVKQLSLEQSQQGDKEFLAEVRLITSIQHRNLVRLLGCCTDGSQRILVYEFMKNRSLDLIIYGKSDQFLNWSTRFQIILGVARGLQYLHEGSHLRIIHRDIKASNILLDEKFQPRIGDFGLARFFPEDQAYLSTQFAGTLGYTAPEYALRGELSEKADIYSFGVLVLEIISCRKNTDLTLPSEMQYLPEYAWKLYERSMVMNLVDPELRENGFVEKDVMQAFHVAFLCLQPHADLRPAMSEIVALLTFKIEMVSTPMRPAFLDRRRKNHDEENNSWEAIS

>A.serrulata_RLCK1 (PCR)

FLVFIYWMRIKQHADTNEKEKEAPNSHQQVEKYNESIEVVKMIVPPQQQRGSMEFISGNLRTISYFDFQTIRRATKNFHRKNLLGAGGFGPVYQGKLADGRLVAVKQLSLEKSQQGEREFLAEVRLITSIQHKNLVRLLGCCTDGSQRILVYEFMKNRSLDLIIYGKSDQFLNWSTRFQIILGVARGLQYLHEDSHLRIVHRDIKASNILLDEKFHPRIGDFGLARFFPEDQAYLSTQFAGTLGYTAPEYALRGELSEKADIYSFGVLVLEIISCRENTDLTLPSEMQYLPEYAWKLYERSMVMNLVDPKLREHGFVEKDVMQAFHVAFLCLQPHADLRPAMSEIVALLTSK

>A.serrulata_RLCK2 (RNASeq)

MALRALFFVLGGIVVLITLLFLVFVYWIRIKQHSEANKKEKEAPTSQQKVEEYNESLEVVKMVAPTQQQRGSMGFISGNLRIISYFDFQTMRRATLNFHHKNLLGTGGFGPVYQGKLEDGRLVAVKQLSLGQSQQGDKEFLAEVRLITSIQHRNLVRLLGCCTDGSQRILVYEFMKNRSLDLIIYGKSDQFLNWSTRFQIILGVARGLQYLHEGSHLRIIHRDIKASNILLDEKFQPRIGDFGLARFFPEDQAYLSTQFAGTLGYTAPEYALRGELSEKADIYSFGVLVLEIISCRKNTDLTLPSEMQYLPEYAWKLYERSMVMNLVDPKLREHGFVEKDVMQAFHVAFLCLQPHADLRPAMSEIVALLTFKIEMVSTPMRPAFLDRRRNNHDEENNSWEAIS

>A.sp328_RLCK1 (PCR)

FHRKNLLGAGGFGPVYQGKLADGRLVAVKQLSLEKSQQGEREFLAEVRLITSIRHKNLVRLLGCCTDGSQRILVYEFMKNRSLDLIIYGKSDQFLNWSTRFQIILGVARGLQYLHEDSHLRIVHRDIKASNILLDEKF

>A.sp_RLCK2 (PCR)

MALRALFFVLGGIVVLFTLLFLVFIYWIRIKQNSEANKKENGAPTSQQKVEEYNESIEGVKMVATTKQQRGTMGFISGNLRTISYFDFQTMRRATLNFHHKNLLGTGGFGPVYQGKLEDGRLVAVKQLSLEQSQQGDKEFLAEVRLITSIQHRNLVRLLGCCTDGSQRILVYEFMKNRSLDLIIYGKSDQFLNWSTRFQIILGVARGLQYLHEGSHLRIIHRDIKASNILLDEKFQPRIGDFGLARFFPEDQAYLSTQFAGTLGYTAPEYALRGELSEKADIYSFGVLVLEIISCRKNTDLTLPSEMQYLPEYAWKLYERSMVMDLVDPELRENGFVEKDVMQAFHVAFLCLQPHADLRPAMSEIVALLTFKIEMVSTPMRPAFLDRRRKNHDEENNSWEAIS

>A.tambacoundensis_RLCK1 (PCR)

ERKKEAPTSHQQVEKYNESIEVVKMIVPTQQQRGSMEFISGNLRTISYFDFQTMRRATKNFHRKNLLGTGGFGPVYQGKLADGRLVAVKQLSLEKSQQGEREFLAEVRLITSIQHKNLVGLLGCCTDGSQRILVYEFMKNRSLDLIIYGNSDQFLNWSIRFQIILGIARGLQYLHEDSHLRIVHRDIKASNILLDEKFQPRIGDFGLARFFPEDQAYLSTQFAGTLGYTAPEYALRGELSEKADIYSFGVLVLEIISCRKNTDLTLPSEMQYLPEYAWKLYERSIVMDLVDPKLREHGFVENDVMQAFHVAFLCLQPNADLRPAMSEIVALLTSK

>A.tambacoundensis_RLCK2 (RNAseq)

MALQALFFVLGGIVVLIVLLILVFIYWIRIKQHAEEGAPTSHQQVEKYNESIEVVKMVVPAQQQRGSMEFISGNLRTISYFDFQTMRRATLNFHHKNLLGTGGFGPVYQGKLGDGRLVAVKQLSLEQSQQGDKEFLAEVRLITSIQHRNLVRLLGCCADGSQRILVYEFMKNRSLDLIIYGKSDQFLNWSTRFQIILGVARGLQYLHEGSHLRIVHRDIKASNILLDEKFQPRIGDFGLARFFPEDQAYLSTQFAGTLGYTAPEYALRGELSEKADIYSFGVLVLEIISCRKNTDLTLPSEMQYLPEFAWKLYERSMVMDLVDPKLREHGFVENDVMQAFHVAFLCLQPNADLRPAMSEIVALLTFKIEMVSTPMRPAFLDRRRKKHDEESNSWEAIS

>S.semperflorens_RLCK_O (PCR)

ILAFIYWMRTKKHAEANNENKKEAPASHQQDKKYNESIEVVKMTVPTQQQRGSMDIISGSLRTISYFDFQTMRKATKNFHRKNLLGTGGFGPVYQGKLADGRLVAVKQLSLEKSHQGEREFLAEVRLITSIQHKNLVRLIGCCTDGSQRMLVYEFMKNRSLDLIIYGKSDQFLKWNTRFQIILGVARGLQYLHEDSHVRIVHRDIKASNILLDEKFQPRIGDFGLARFFPEDQAYLSTQFAGTLGYTAPEYAIRGELSEKADIYSFGVLVLEIICCRKNTDLTLPSEMQYLPEYAWKLYEKSMVMDLVDPKLREHGFVEKDVMQAFHVAFLCLQPHADLRPAMSEIVALLTFK
