## Supplementary File 3 for "A Receptor Like Cytoplasmic Kinase evolved in *Aeschynomene* legumes to mediate Nod-independent rhizobial symbiosis"

**Supplementary File 3. Sequences synthesized for *AeRLCK2* and A*eCRK*.**

>AB_ProAeRLCK

ACGGTCTCGAAATAGAAAGGTTTAGAGGTTTGAACGTTCGGCTATAAAGTCTGGATTTTCTTTGCTTTAGGCATATATCCTCGGTAGTTATGTATATAGAGTTATGCTATTTTTATATAAGTTCTATGAAATATAGTACAGAATTACAGATTCACACTTTATTTATGTTTAGATGAAAGGTCGTTTCTACCTAATTGGGACTCACATTTTTTAATAGGACGAGGAAAGATTGATTGATCAGTCCAAAAATTAAGTCCTTAATAATACTATATATACAATTTGCAGTTCTTTCTTTTTTACCACCATATGAAAAATAAATTATTTGTGAACATCAATATATGAAAGTATGAGGTAAGAGGTGATTATATAGAGATCAGAAAAGTAATTTTAAAATATTGTAACATAATAATCTTTTTTGTTTTGAAATTTTCTAAGAAGAGATTTTCAGAGTTCAAGGTATAACTATTATTTGTTTTTTTTATTTCCATAGACGAATAGAAGATGTAGAACATAGATAAACATAGGGATTAAGATTAATAAATAAATAAATAAAGTGGGCTGGACTGATAGTTGATAGCGATTTGGGAAAGCAACCATTGTCATCATCCTCTCTACTTCTCTCGCTTTTCGAAAGAGCAACGAATTTCACTGAACAGCTTGCTTCTTCTTCCTCCCACAAGAACTGTTCCAATGTCTCTACGCTTTTCTTCATCTCACTGTTCTTTCTCTCCGTCTTCATCTTCAGAATCTTTCCCTTCACACAGATCTCGCCCTCCACCATTCTACTCTACCTATCTGGCTATCTCTATCCCCGATAATAACACTATCACCCAAACCCAACAACAACAACCCTTAGAACCCGCAGAACCGTAGCCCGCCACCGTGCCCACCGACACCGACACCTCCGTCGATGACGATGTAGGCATTCAAGCAGCTTCTTTTACCAAAGCGGTGGTTGATAACGCTTATGCTGATGCTAATGATGGCAATGGTGATGTAGCGCTGAAGCAGCAGCAGTTAGAGTGCGATTTGTACAGGGGAACGTGAGTGAAAGACGAGGATTACCCGATTTATCAACCGGGTTATTGCCCTTACGTAGCAGAGGCTTATGACTGCCAAATTAATGGAAGAAACGATTCCGAGTACACCAAATGGCGCTTGAAGCCCGACGGCTGTGATCTTCCCAGGTATCTATCCAATTTCCTTCTCTTTTTTTGTTAGGTCAAACTATCCAATTTCCTTAAATCAATTCCTTTACTTCCATGCTACATTGGGGCTTCACTGTGTCAATTTCCAGCAATTAAAATGTAGTGATGTTTTGTATGTGATAGGCATTGATTCCAGAATGTATGATAGGGATTTTAGTTACTTTGGTTGTTCCACGTATTTGTGGACTTATTAAATCACCAATTCATCATGAATGATTTGAAACTCTTGGGAGTGGTGTGCTTTTTTGTTCATTATTTCCACATTTCATTGTAAAGAAGTATTCATTGCTAATTCCAGAAGATGTGCTCTTTCTTTGGTAGATGAGAGGTGCAAAAATCAAATTTGGATCTTTCTATTGGACGGCATCTTTATGTGTCCCTTTAGCTTTAATTGGATCGTATGACAATATTTACATTGTTCCGATTAATTCTATAGGACACCCTTGGAATAATTCTCCCATCTCTCCCAAGTTTAATTTTTGGTTTTTCAATGTAACAAATTGACAACAGCAACAAAAAAGAAATAAAAATGAAAACTGAAAAAAAAAATAGTCGGAAAATGGTTGTTTTTGCCAGTTGTTGAATGAACTTTGTAAGTTGTAACTCTTATAAATGGCCATGGCAATATTGAGTTGACCAAGTCTTTCCAACTTCAACTTCAACCAGAATGCCTCAAATACTATTCCAAGGTTTTATATTTTTATCAGATTGTTTTTTCCACTCTAGGCTTCATAGTGTAAGGATCGAGAATTCAATTCCATGCATTGTGTGCACCATTCATTTTGAGTTTTGAATTGGGGTTACTATGGTACAGGCCTAAACAAGCTAAGTATCCATTTGGTAACTCTCTGAAAATCGAGGTGTTGGTAAATAACATGTAAATAAGGGATCTTTTATAGCTTTTACAGATTTTTTTCATACTATTATTTTGAATCATGTTCAAACAGGTTTCATGGCAGAGTCTTGTGGCTGATTGCAGAGAATCTGATGCAGGTGCAAAATATAATTAGTAAGTTAAGTGTGTTGAAGAATTGTATCAAGGGAATAAAAGTCCCACAATGCTTAGGTTAAATAACTTATCCCTTAGAAGTCCCACATTGCTTAGTTTTCAAAGGTATCCCCTTAGTACTAGTATAAATAACCCCATATCTTTGTTTGACAATTTGAATTGAATGGCTACTTGCTTCAAATTAAGTGTATGCTTATTTCCTCTAAGCTCTTATTAAAGAGTAAGAGGTCTAGTTAAATTTCCTTGGCTTGGCCAACCAAGGTAGTTTAATTGGCTTTTTTCACTGTAGTGAAGTTATTACCCTTTCCACTGTTCGTCTCTGTATCCTCCTCTCAAAGGAGACCGT

>AB_ProAeCRK

ACGGTCTCGAAATTCTACCATATCCTCAATCAATGACATGTCAAAGAGTCTTTTTTTCCCTCGATAGTATATATTTATATAAAGAGTTTGATCTTTATTCTGTGAAATTAATTATTATGTACTAATTTATTCGTTTTTATAAAGTTTCATAACAAATCATTCAGTTTTGTGTGTGTTGTGTATCCAACTTGCCTTTTAATGGATGAAAAAAGCTTACAAAATACCAATGCACTCATTCTAAGGAATTTAATTTAGACTTAGTTTACGCTTTCGATAAGGTCTATGAAAAATTCTTAGATGGTCTTGGATCACATTTAATGCATGACTCAGGAATATTTATAATCACATATTTTGATTTATAATATTCAAAATTTATATAAAAAAATATAAAAAATGTGGTTGTAAATATTTTTAGACCATTTATTAAATGTGATCCAGAATTGTATAAAAATTTCTCTAAGATCTAGTGTAACAAAAATAAATAAATAAATAAATTGACAGGTGATGAGGAGGACTAAGTATGATATTTGTTTAAAAACAAAATGTCATATTCATCTTCATCATAAGCACAAGTTGCTATTTTGCAGATTAAACAAGAAATGTGCTAGCTGGTAAGTGTACAATAAAATAAGGGTTAAAGCTACGTTTGATGAAATTATACATGAATCCCTCCTAGTTTCAAAATTTGATTAGAGTAGGTCAGTTTAATAAAAATATATACTGAACTCCTCTCTAAGTGTTTAATTGTTACTCAAGTTAACAATGAAATTGACAATAAAAAATACACTGATAAATATGAAATGACTAAATGATCATTTTATTAAGTTATTAAATTATTAATTAATAAATATTAAGGAGATGATTTCTCTCAACTTAATTTAGTGTAGGATAACACTGAAAGTTTTATATTTATATTTTTTTTATTTACTAAAATGTCTATCAATTATTTTATTAACTTGACCAACAATCAAATGATTAACCAAAATGTCAGTGTATATTTTTATCAAATTGAAAGAAGTTTAAATGTATATTTTAAAATTAGAGAAAAAATCTGGTGTAGTTTGGTTAAACTTCATGTGTAATTGTAATTTACTTAACTTTTGTCCTAATCGAAATTCCAAGAGTTAGCCTTAATGGTAAATAGTAAACTCTTAGAACACAAAAAGAGTATAATAATCAACACTACAGTGAGGTCCATTTAGATTTTCTCAGTGATTTTTGGTATGAAGCAATGGTTGAGAAGACACTTGTATCCTATAAACAATTACACAATAACCTTTCAAGACATCATTGAGCTGAAAGAGACATATATTCCCTCACTATGTACTCTCTCAATGAAAAGTTTTTCATATCTCAGTAGCAAATTGATCAAAGAAACCATGCAGTAATAAACACAAAGGAGACCGT

>BC_AeCRK_nostop

ACGGTCTCGCAAAATGTATCTCTTCCACAAGAAGTACAGAGCTTTTGCTTTGGTTTTGATCTC

AATCATTGTTGTTCTAGTCAACCCCAATAGTTCTGAGCCAACCTACAACT

ACCATATATGTGAAGATCAATTAGATAATGCAACCAATGCTAATAATTTC

CAATCTGATCTAACTTCCCTCTTGGATTCTTTATCTCTTAAAGCTTCACT

TCACAGCTTCTTCAATGATAGCATCAGTACAAGGCTCTATGGCCTCTTCC

TCTGTAGGGGTGATGTCTCTATTGACACTTGCCGAAACTGCACCAAAACT

GCAAGCCAAGAAATCAAAACCCTTTGTGTATCAAAGTCCAGTGCAATCAT

CTGGTATGACGAGTGCATGCTTCGCTACTCCGACAAACAATTCTTTGCAC

AAGCACAGACACTTCCTATGGCATTTGCTTGGAACATTGAGAATAGAACA

TCAGCTGCTGAAACAGATATTGATGCACAAGCTCTGATGTACCAGTTAAT

AAAAGAAGCCTCAAACACGGATATACTGTTCAAAACAAAGAAATCTGAGG

GGAAAGATGAGTCTGAGAATAGGTATGGACTTGTGCAGTGCACAAGAGAT

ATAAATGACAGTTTGTGCAGCTCTTGTTTGTCACAGTTGATGAACAAGGC

GGAGCGATGTTGCCAACGGACCGTTGGGTGGCGCATATTGGCTCCAAGCT

GCAATATTAGGTATGAGAACTATACCTTCTATCAGCAAACTTCGGAGCCA

CCTTCTGAAGGGAAGCGAGGAAGCAATAGAGCAAGGATTATATTCCACAC

AGTTTTTCTGATAGCAGTTATTTTGGCCTTTTTAAGTTTAGGGTGGTGTG

CATGTCCTATCTCAAAAATAAGAAGAAGATTAAAAGGGGGTGGGACAAGT

GATGAAATTCAATTCAAGAATTTGATGAGTTCAAGTAGGTCAGACTTTGA

GGAACAAAGGAAGAGTTCAGTACATAAAGACAACAGTGGAGAAGTGCAAT

ACTTTAGTTTAGGCATGATTAGGGTTGCTACCAACAATTTTTCCAATGAA

ACCAAGCTTGGGGAAGGTGGCTTCGGACCCGTTTTCAAGGGGAAGCTGAT

TAATGGGAAGGAAATAGCAGTTAAAAGGCTTTCATTCAGATCAAGGCAAG

GCCTTGAAGAATTCAAGAATGAAGTGATGTTAATTGCAAAGCTTCAACAC

AGAAACCTTGTTAGGCTCTTGGGATGCTGCCTTGATAAAAATGAAAAGCT

TCTTGTCTATGAGTACATGGCCAACACTAGTCTTGATGCTTTCCTCTTTG

ATCCAACCAAGCGTAGAAAGCTTGATTGGCCTAATCGGGCGAAGATTATA

AATGGAATTGCTAAAGGTCTTCTATATTTACATGAGGACTCAAGGCTGAA

AATCATCCATAGGGATCTAAAAGCTAGCAATGTATTGCTAGATGAAGAGA

TGAATCCAAAGATATCAGACTTTGGAACTGCTAGGATTTTTTGTACTAAT

CAAATTGAAGCAAGCACTGAAATAGTTGTTGGCACATATGGATATATGGC

ACCAGAATATGCCTTGGAAGGAATATTTTCCATCAAGTCTGATGTTTACA

GCTTTGGGGTACTGATGCTGGAAATTTTGAGTGGGAGAAAGAACAATGGC

TTCTTCCAAAAGGACCGTGGTCGTGATCAAACCCTTCTATCCTATGCATG

GCGACTCTGGAGTGAAGGGAAGGAAGTGGAATTTCTGGACCCGGTATTGG

TGAATGCATGCCCTATACATGTGGCGTTGAGATGGATCCATATAGGGTTG

TTGTGTGTTCAAGAAAGGCCCAATGACAGGCCCACTATGTCCTCCGTTAC

TCTCATGCTTGGAAGTGCTATAAATCTTCCTCGGCCTTATGCTCCTCCCT

TTTCCTTAAGCATATATCACTTTTCATCTGAGGAAGAATCTTCAACAATA

GGAGATGATGATATTCAATTTCAACCCTCTTCCAGCTCTGTTTCTACTTT

CTATGGTGAGAGACCGT
